# Single-Cell Omics to Uncover Cell-Type-Specific Gene Regulatory Networks Driving Mucilage Production in Specialized Roots

**DOI:** 10.64898/2026.09.29.755433

**Authors:** Rafael E. Venado, Marina Kotvanova, Chandrani Kumari, Prakriti Garg, Saptarshi Pyne, Rubens Diogo, Tess Kern, Christopher Dervinis, Wendell J. Pereira, Matias Kirst, Sushmita Roy, Jean-Michel Ané

## Abstract

Aerial roots of certain maize (*Zea mays*) and sorghum (*Sorghum bicolor*) genotypes secrete a carbohydrate-rich mucilage that sustains nitrogen-fixing bacteria, and the underground roots of most plants secrete mucilage at their tips. In both settings, border cells released from the root cap produce this mucilage, yet the genes controlling their formation remain largely unknown. We sequenced the RNA of about 23,000 nuclei from mucilage-secreting roots of maize, sorghum, and *Medicago truncatula*, sampled before and after secretion. Each species yielded 10 to 15 clusters, which we annotated with published markers and with new markers that we validated by *in situ* hybridization in the two cereals and by promoter-reporter fusions in *M. truncatula*. We then inferred modular and cell type-specific gene regulatory networks. Transcriptional programs diverged across the three species except in the quiescent center and the meristem, and in the clusters containing border cells, where a shared set of regulators emerged. Silencing three of five candidate regulators reduced border cell number, while silencing a partially overlapping set of three regulators reduced root diameter in *M. truncatula*. Silencing the C2H2 zinc finger protein MtrunA17_Chr1g0148421 also reduced border cell viability. A maize insertion mutant of *bzip89* released fewer border cells than the wild type. These networks offer entry points for tuning mucilage production in cereal and legume crops.

## INTRODUCTION

Cereals are the most widely cultivated crops worldwide, providing approximately 50% of the calories in the human diet (Awika 2011). Maize (*Zea mays*), wheat (*Triticum aestivum*), rice (*Oryza sativa*), sorghum (*Sorghum bicolor*) and millets collectively account for 90% of total cereal production (FAO Cereal Supply and Demand Brief). These cereals require substantial inputs, particularly synthetic nitrogen fertilizer, whose production is energy-intensive and whose overuse poses serious environmental risks. Remarkably, certain maize landraces and sorghum accessions can acquire a significant portion of their nitrogen through biological nitrogen fixation (BNF), supported by diazotrophic bacteria colonizing carbohydrate-rich mucilage secreted by their adventitious aerial roots (hereafter referred to as aerial roots) (Van Deynze et al. 2018; Venado et al. 2023). The mucilage is released when aerial roots are exposed to moisture and creates a specialized microenvironment that sustains nitrogen-fixing communities, with root diameter serving as a positive predictor of mucilage output and BNF capacity (Van Deynze et al. 2018; Bennett et al. 2020; Pankievicz et al. 2022; Wolf et al. 2024; Venado et al. 2025a, 2025b).

Mucilage secretion is not unique to aerial roots. It also occurs at the tips of underground roots across phylogenetically diverse plant species (Hawes and Pueppke 1986; Hawes et al. 2002). In both contexts, border cells (BCs) produce and release mucilage; these specialized cells originate in the outermost layer of the root cap and detach from it (Hawes and Pueppke 1986). Once released, BCs remain metabolically active and form, together with their mucilage a structure now termed the root extracellular trap (RET), a matrix embedding cells in a network of polysaccharides, defense-related proteins, extracellular DNA (eDNA), histones, and secondary metabolites (Hawes et al. 2000; Hawes et al. 2012). RETs function as a frontline defense against soilborne pathogens by physically trapping and immobilizing invaders in a manner functionally analogous to neutrophil extracellular traps (NETs) in mammalian innate immunity (Brinkmann et al. 2004; Hawes et al. 2016). In pea (*Pisum sativum*), for example, the RET traps zoospores of *Aphanomyces euteiches*, induces nematode quiescence, and prevents fungal penetration of the root cap (Cannesan et al. 2011; Fortier et al. 2023), with eDNA playing both structural and antimicrobial roles (Wen et al. 2009; Tran et al. 2016).

Beyond pathogen defense, BCs are also major secretory cells whose chemical output shapes the broader rhizosphere environment. Spatial metabolomics profiling of *Medicago truncatula* (hereafter referred to as Medicago) roots has revealed that, despite having lower overall internal metabolite diversity than other root tissues, BCs are enriched in secreted compounds with clear roles in microbe recruitment (Kranawetter and Sumner 2025). These include tryptophan, a chemoattractant for the rhizobium *Sinorhizobium meliloti* (Webb et al. 2017); 7,4′-dihydroxyflavone (DHF), which induces rhizobial *nod* genes while also functioning as an antimicrobial agent (Maxwell et al. 1989; Watson et al. 2015); and organic acids such as malic and citric acid, which recruit beneficial bacteria and mobilize soil-bound nutrients (Rudrappa et al. 2008; Kranawetter et al. 2021). This dual capacity of BCs, pathogen exclusion and symbiont attraction, makes them uniquely positioned at the intersection of plant immunity and rhizosphere ecology.

The number and composition of BCs vary substantially across species. *Arabidopsis thaliana* and its close relatives in the Brassicaceae do not produce true BCs but instead release BC-like (BCL) structures that remain clustered and share only a subset of BC functions (Driouich et al. 2010). These differences limit the use of *Arabidopsis* as a model for BC biology, making species such as Medicago, pea, maize, and sorghum much more informative systems. Despite their biological and agronomic importance, the gene regulatory mechanisms governing BC development and mucilage production remain poorly understood, largely because root tissues are underground, composed of heterogeneous cell populations with distinct transcriptional states, and difficult to resolve using conventional bulk transcriptomic approaches. Furthermore, the Gene Regulatory Networks (GRNs) that specify cell type- and context-specific expression are largely unknown for BCs.

Single-cell RNA sequencing (scRNAseq) and single-nucleus RNA sequencing (snRNAseq) now make it possible to capture cell-type-specific transcriptional profiles at scale, and have been applied successfully to roots of *Arabidopsis* (Denyer et al. 2019; Shulse et al. 2019), Medicago *(Cervantes-Pérez et al. 2022; Pereira et al. 2024)*, *Lotus japonicus* (Frank et al. 2023), and major cereal crops. However, no study has yet applied this technology to resolve the transcriptional programs and the GRNs governing BC development and mucilage production across phylogenetically diverse species. Although BCs occur across different root types and species, mucilage formation appears to be an evolutionarily conserved strategy for plant-microbe interactions that depend on environmental cues such as humidity, mechanical stimuli, and microbial signals, suggesting integration with local environmental conditions. The availability of scRNA-seq data also enables GRN inference in a cell type-specific manner, enabling the identification of key transcription factors and their target genes within specific cell populations (Kim et al. 2023). Experimental validation remains essential, and Medicago’s relative tractability for *Agrobacterium rhizogenes*-based transformation makes it an attractive experimental system for functional testing of conserved regulators before translation to legume and even cereal crops.

In this study, we analyzed the transcriptional profiles of mucilage-producing roots, both aerial (maize and sorghum) and underground (Medicago), to identify regulators associated with BCs. Using a comprehensive multi-species analysis pipeline, we defined cell clusters and novel cell markers across these species. In each species, we identified 10–15 distinct clusters further annotated using known cellular markers. We validated additional novel markers by cell type using *in situ* hybridization in maize and sorghum and applied a promoter-reporter system in Medicago. We then constructed GRNs using Modular Regulatory Network Learning with Per Gene Information (MERLIN) to define gene expression modules and regulators for individual genes and modules, and Single-Cell Multi-Task Network Inference (scMTNI) to identify cell type-specific GRNs. We identified and functionally validated key transcription factors whose loss reduces BC number and viability. Our results reveal a conserved transcriptional program controlling BC development across monocots and dicots and provide a foundation for engineering mucilage production and rhizosphere chemistry in both legume and cereal crops.

## RESULTS

### Multi-species single-nucleus transcriptome on mucilage-secreting roots

We characterized the cell type-specific transcriptional profiles of mucilage-secreting roots from three plant species: maize, sorghum, and Medicago (**Figure 1A**). These profiles examined transcriptional changes before and after mucilage production (hereafter, dry and wet conditions). For the cereals, we collected root tissue from the aerial roots of the maize landrace Oaxa524 (Venado et al. 2025) and the sorghum accession IS23992 (Venado et al. 2025). In contrast, for Medicago, we used underground roots of the genotype Jemalong A17. Due to the tough nature of aerial roots, we optimized the nuclei isolation from our previously reported protocol (Conde et al. 2021) to effectively remove debris and other organelles while preserving the nuclei integrity (**Supplemental Figure 1A**). We then used Fluorescence-Activated Nuclei Sorting (FANS) to select single nuclei based on forward and side scatter, aiming to capture at least 40,000 nuclei per sample (**Supplemental Figure 1B**). To confirm the presence of BC, we exposed root samples to water and verified their release. BCs from maize and sorghum are similarly enlarged in shape, with no apparent differences between them, whereas those from Medicago are smaller and exhibit a distinctive ‘bean-like’ morphology (**Figure 1C**). We processed high-quality nuclei using the 10X Genomics droplet-based assay and prepared and sequenced the libraries (**Figure 1B**).

**Figure 1.**
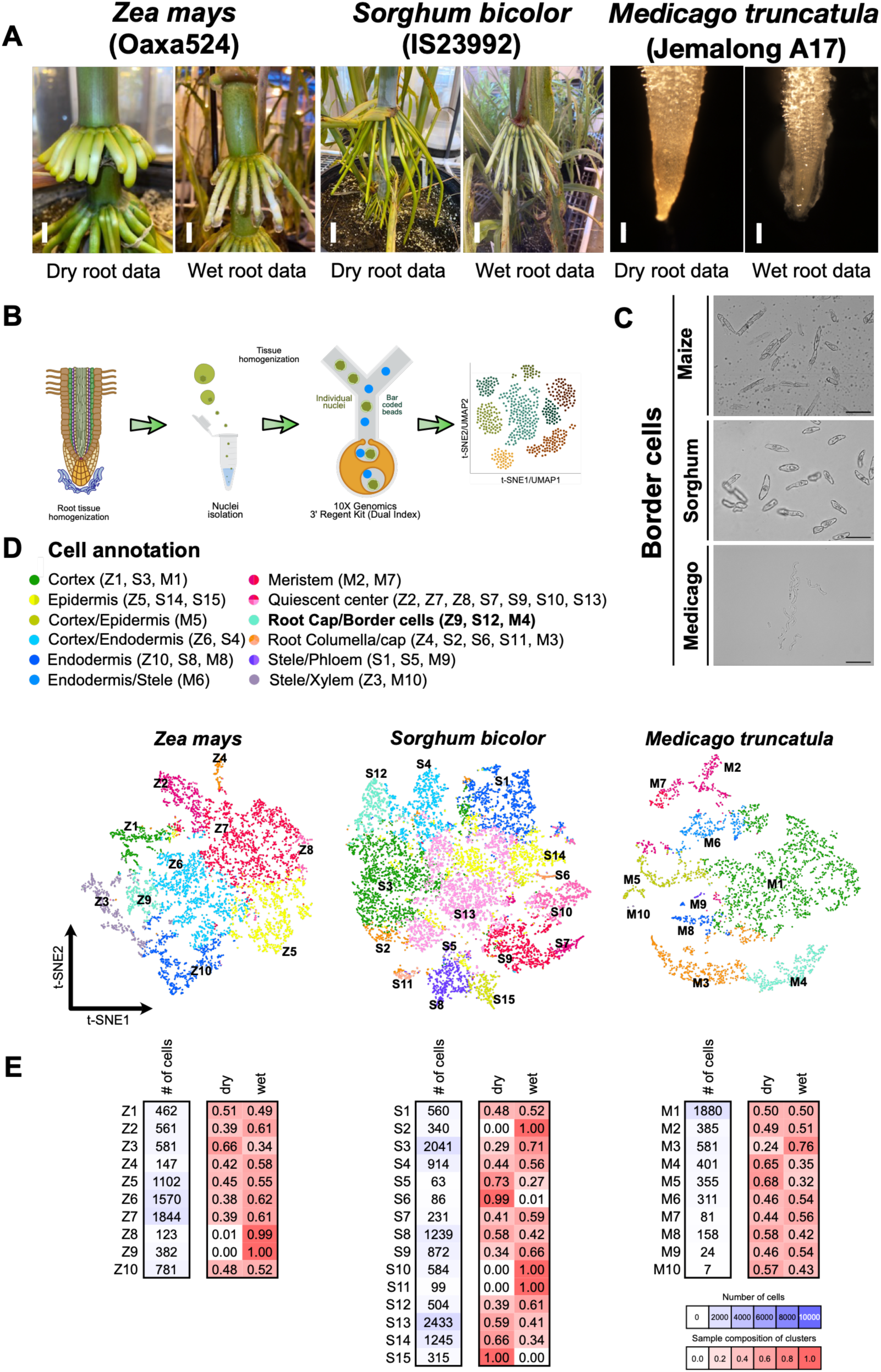
Single-nucleus RNA-seq of mucilage-producing roots in maize, sorghum, and Medicago. **A)** Adventitious nitrogen-fixing aerial roots of maize and sorghum shown before and after mucilage release. Underground roots of Medicago shown before and after mucilage release at the root tip. Scale bars = 1 cm maize and sorghum and 1 mm for Medicago. **B)** Schematic diagram illustrating the isolation of nuclei using 10x Genomics technology and the downstream analysis pipeline. **C)** BCs from the three species. Maize and sorghum BCs are elongated and irregular in shape, whereas Medicago BCs are smaller and bean shaped. Scale bar = 100 µm. **D)** Single-nucleus RNA-seq (snRNA-seq) data visualized using a t-SNE plot of the combined dataset. Final cluster annotations were based on differential gene expression (DEG) analyses and known cell-specific markers. Clusters are labeled with standard color codes and species-specific identifiers: maize (Z1 - Z10), sorghum (S1 - S15), and Medicago (M1 - M10). **E)** Sample composition of inferred clusters. For each cluster shown in panel (D), the total number of cells and the fractions of cells contributed by the dry and wet samples are displayed. Rows correspond to clusters, the blue color scale indicates cluster size, and the red color scale indicates sample proportion.

After sequencing and quality control, we mapped reads from the dry and wet samples to each species reference genome using CellRanger. Reads mapped at a high rate of 46-90% to reference genomes, with only maize dry and wet sample reads at 46% and 63% (**Supplemental Table 1**) and all other samples at 87% or higher. After filtering cells and genes in each sample for depth, we merged the sample-specific matrices to generate a single species-specific expression dataset of 4,183, 7,553, and 11,526 nuclei in Medicago, maize, and sorghum, respectively, totaling 23,262 nuclei across all datasets. These datasets included 25,126, 26,131 and 21,224 genes in Medicago, maize, and sorghum, respectively. We used these matrices for cell clustering, annotation, GRN inference, and downstream analysis.

### Mucilage-secreting roots exhibit a diversity of cell types validated with promoter assays and *in-situ* hybridization that exhibit conserved and diverged evolutionary profiles

To characterize root cellular diversity before and after mucilage production, we applied Non-negative Matrix Factorization (NMF), a popular dimensionality reduction and analysis tool for single-cell omics datasets (Stein-O’Brien et al. 2018; Iyer et al. 2022). Using NMF, we identified 10 distinct clusters for both maize and Medicago, and 15 clusters for sorghum (**Figure 1D**, **Supplemental Figure 2**). Most clusters contained cells from both wet and dry samples. However, Z9, Z8, S2, S10, and S11 contained only cells from wet samples, whereas S6 and S15 contained only cells from dry samples **(Figure 1E**).

To annotate each cell cluster, we identified differentially expressed genes (DEGs) for each cluster (**Methods**). In parallel, we created literature-based cell type-specific gene sets in Medicago and maize and used orthology to transfer annotation to sorghum. We assigned a cell type to each cluster based on significant overlap between cluster-specific DEGs and literature-curated gene sets (**Figure 1D**). Given that aerial roots may contain a greater diversity of cell populations, we simplified the annotation by defining 12 distinct cell populations. We identified these populations based on previously reported marker lists. The defined cell types include the cortex, epidermis, cortex/epidermis, cortex/endodermis, endodermis/stele, endodermis, meristem, quiescent center, root columella/cap, stele/phloem, xylem, and root cap/BCs (**Figure 1D**).

We next used orthologous gene groups (orthogroups) to assess how cell-type-specific gene expression was conserved across species. We selected 4,181 orthogroups with no more than one gene duplication in the corresponding gene tree. These included orthogroups with one-to-one mapping across all species, as well as orthogroups in which a gene was absent from one species, consistent with a single gene-loss event. We compared cell-cluster pseudo-bulk profiles across Medicago, maize, and sorghum to assess conservation across the three species using all-by-all Pearson correlation of the pseudo-bulk profiles (**Supplemental Figure 3A, B**). When clusters were grouped by tissue annotation, the QC/meristem block emerged as the only strong, coherent diagonal pattern across species, reaching the highest cross-species correlation scores, indicating a clearly conserved transcriptional program. Other tissues (cortex + endodermis, stele/xylem, epidermis, BCs/cap/columella) showed weaker, more fragmented diagonal block structure, with maize-sorghum correlations generally higher than either species with Medicago. However, these signals were patchy and rarely formed continuous blocks. Without tissue grouping, heatmaps revealed broad interspecies similarity, forming a maize-sorghum block (reflecting grass-specific conservation) and lower overall similarity to Medicago. Against this background, the QC/meristem relationship remained visible as an off-diagonal band, reinforcing it as the most conserved program among the three species. Relaxing orthogroup filters from 1-1 ortho to include 8533 orthogroups with one duplication and/or loss decreased correlation dispersion and further fragmented non-QC tissue structure, but the QC/meristem signal remained robust (**Supplemental Figure 3C, 3D**).

To complement these global patterns, we examined per-orthogroup pseudo-bulk profiles by clustering genes based on their expression value across species (**Methods**). One cluster of orthogroups displayed QC/meristem-restricted expression across all three species, including known QC markers *Zm00001eb086340*, *Zm00001eb307330*, and *Zm00001eb336890*, confirming a conserved meristem signature (**Supplemental Figure 4A**). Another orthogroup cluster (**Supplemental Figure 4B**) showed broadly elevated expression across multiple cluster types but consistently higher levels in maize and sorghum compared with Medicago, suggesting a grass-biased regulatory shift rather than strict tissue specificity. A third orthogroup (**Supplemental Figure 4C**) exhibited sharp, tissue- and species-restricted enrichment confined to sorghum root cap/columella, indicating divergence of this program in sorghum. Conversely, another orthogroup (**Supplemental Figure 4D**) was enriched in BCs across species while remaining low in sorghum root cap, suggesting that BC-associated expression can remain conserved even when the related root cap module diverges.

Taken together, these results show that root cell-type transcriptional programs are conserved across species for several gene sets, with the strongest conservation in QC/meristem cell populations, while other tissues show weaker and more fragmented conservation patterns.

### Validation of the cluster annotation using promoter assays and *in situ* hybridization

After computational cluster annotation and evolutionary analysis, we experimentally validated our results by selecting differentially expressed genes (DEGs; **Supplemental data set 1**) unique to specific clusters (**Supplemental Figure 5**) and confirming their annotation with special emphasis on root cap and BC markers across all three species. In Medicago, we generated transgenic Medicago roots expressing a 2 kb promoter region from each marker gene fused to a nuclear-localized YFP (NLS-2xYFP) reporter. These constructs showed strong fluorescence in the corresponding cell layers (**Figure 2A, B**). As expected, the BC marker was also detected in the root cap, reflecting the origin of BCs from this region (**Figure 2C, D**).

**Figure 2.**
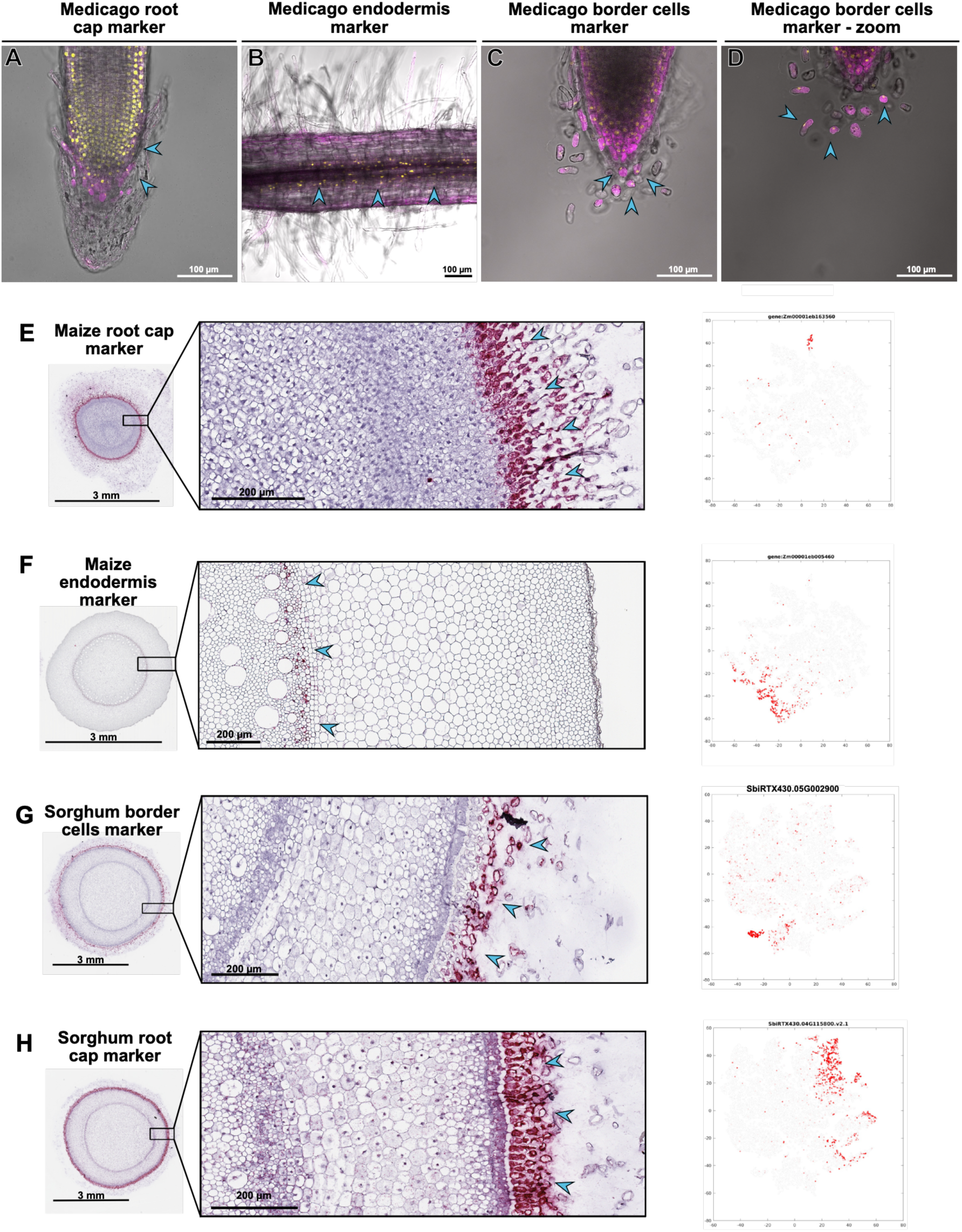
Marker Validation Across Species. Marker genes from distinct expression clusters were selected and validated in the three plant species: maize, sorghum, and Medicago. In Medicago, promoter-reporter assays were performed by expressing the promoter region of selected marker genes driving a nuclear-localized fluorescent reporter (NLS-2xYFP, yellow). Transgenic roots were identified using a transformation marker (NES-2xmCherry, magenta). In maize and sorghum, *in situ* hybridization was used to localize marker-gene expression. Signals were visualized (red), with hematoxylin used as a counterstain (blue). **A)** Representative image of a 2 kb promoter from a root cap marker (MtrunA17_Chr8g0379831). Blue arrows indicate root cap cells. **B)** Representative image of a 2 kb promoter from an endodermis marker (MtrunA17_Chr2g0321181). Blue arrows indicate endodermal cells. **C–D)** Representative images of a 2 kb promoter from a BC marker (MtrunA17_Chr4g0011791). Blue arrows indicate BCs. **E)** Overview and magnified region showing in situ hybridization of the maize root cap marker (Zm00001eb163560), with corresponding t-SNE plot indicating cluster expression in maize snRNA-seq data. Blue arrows highlight red hybridization signal in the root cap. **F)** Overview and zoom of in situ hybridization for the maize endodermis marker (Zm00001eb005460), with t-SNE plot showing marker expression in snRNA-seq clusters. Blue arrows indicate red signal in the endodermis. **G)** Overview and zoom of in situ hybridization for the sorghum BC marker (SbiRTX430.05G002900), along with t-SNE plot of expression in snRNA-seq clusters. Blue arrows point to red signal in BCs H) Overview and zoom of in situ hybridization for the sorghum root cap marker (SbiRTX430.04G115800), with t-SNE plot showing expression pattern. Blue arrows indicate red signal in the root cap. For the promoter assays, five independent transgenic plants were analyzed. For the in-situ hybridization experiments, three aerial roots were examined

For the cereal species, marker validation was initially performed using RT-qPCR on two tissue types, the root tip and a section from the elongated root zone (**Supplemental Figure 6A**). In maize, we validated markers for the root cap (*Zm00001eb163560* – Nucleotide sugar transporter-KT1), cortex (*Zm00001eb027760* – Pollen-specific protein), and endodermis (*Zm00001eb005460* – Glutaredoxin family protein). In sorghum, the selected markers corresponded to the root cap (SbiRTX430.04G115800 – callose synthase), BCs (SbiRTX430.05G002900 – Protein Kinase Domain-Containing Protein), and cortex (SbiRTX430.01G385000 – PTHR11977:SF40 – Villin). Markers associated with root cap and BCs were expected to show high expression in the root tip, whereas markers for the endodermis, epidermis, and cortex were expected to be highly expressed in the elongated root zone. Our results confirmed these expression patterns in both cereal species, with one exception: the cortex marker in sorghum showed high expression in both tissue types (**Supplemental Figure 6B and C**).

To further support our findings and demonstrate spatial specificity, we performed in situ hybridization. Aerial roots were sectioned approximately every five mm to ensure proper tissue fixation and to distinguish developmental zones. In maize, we tested an endodermal marker and a root cap marker. The root cap marker was strongly expressed in the young root cap tissue, with minor diffusion into adjacent BCs (**Figure 2E and Supplemental Figure. 7)**, while the endodermal marker was localized as in the elongated root zone and absent in young sections (**Figure 2F and Supplemental Figure. 8**). In sorghum, the BC marker exhibited strong signal in the mucilage zone of the root tip but was absent in later sections (**Figure 2G and Supplemental Figure 9**). Similarly, root cap markers were highly localized to the outer cells of the initial root sections, with no signal detected in more mature regions (**Figure 2H and Supplemental Figure 10**). Together, the promoter-reporter assays and in situ hybridization analyses validated our annotations, confirming the cluster identities in all three species.

### Examining coarse and fine-grained cell lineage structure

To characterize relationships among the annotated cell types, we first constructed a Minimum Spanning Tree (MST) based on each cluster’s pseudo-bulk expression profile. We then integrated the cell type annotations described above with the inferred MST, refining it based on prior literature of root cell-type organization and functional relationships, to obtain the final cell lineage trees **(Figure 3A, Supplemental data set 2**).

**Figure 3.**
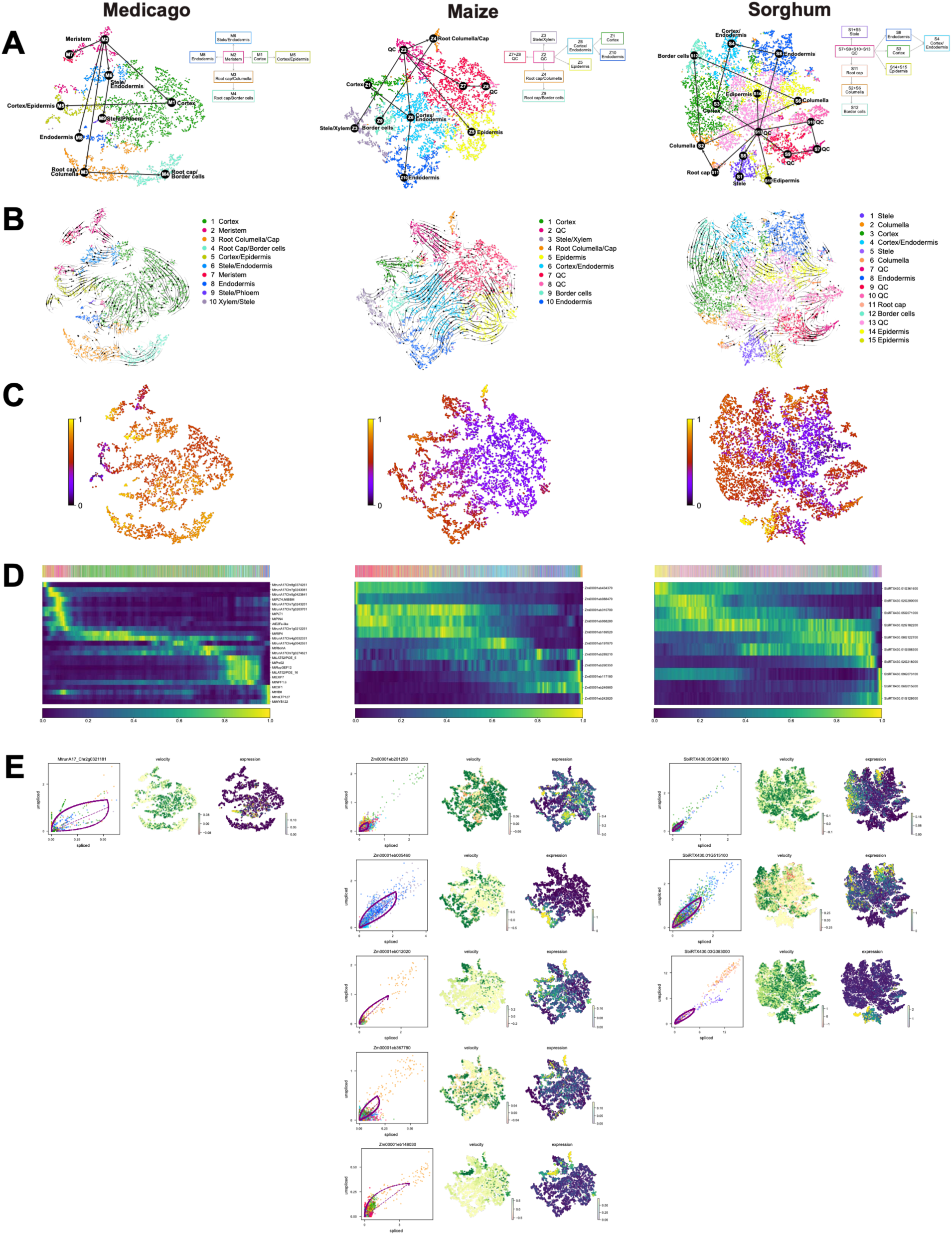
RNA velocity analysis across Medicago, maize and sorghum root cell populations. Columns correspond to Medicago (left), maize (middle) and sorghum (right). **A)** Two-dimensional t-SNE embeddings colored by cell cluster annotations together with schematic lineage trees summarizing the inferred relationships among clusters. Lineage trees were constructed based on cluster identities and the developmental assumption that the meristem or quiescent center gives rise to most differentiated root cell layers. **B)** RNA velocity stream plots projected onto the same t-SNE embeddings, showing the inferred direction of cell-state transitions within each species. **C)** Latent time inferred from the dynamical model and visualized on the same embeddings, highlighting the progression of cells along developmental trajectories. **D)** Heatmaps showing expression dynamics of selected marker genes ordered by latent time for each species; columns represent cells arranged from early to late latent time and rows represent genes. The color bar above each heatmap indicates cluster identity. **E)** Phase portraits for selected orthologous genes across the three species, with corresponding velocity and expression patterns projected onto the t-SNE embeddings. Phase portraits show spliced versus unspliced transcript abundances used for dynamical modeling; individual cells are colored by cluster identity, and the purple curve indicates the fitted dynamical model.

We complemented this coarse cell cluster level analysis with RNA velocity analysis for all three species using scVelo (Bergen et al. 2020) (**Figure 3B**-**E**). Across Medicago, maize, and sorghum, the proportions of spliced versus unspliced reads were consistent within species and across clusters, indicating sufficient kinetic resolution for dynamical modeling (**Methods, Supplemental Figure 11**). Projection of velocity vectors onto the NMF-based tSNE embedding (**Figure 1D**), displayed as streamlines, revealed directional transcriptional flows originating in meristematic populations and terminating in tissue-specific clusters (**Figure 3B**, **left**). In Medicago, arrows originated from the Meristem clusters (M2 and M7) and swept toward Cortex (M1) and Cortex/Epidermis (M5) on the periphery, Endodermis (M8) and Stele/Endodermis (M6) along the central axis, and Stele Phloem (M9) at the tip. Root cap/columella (M3) and Root cap/BCs (M4) resolved as distinct terminal branches. The per-cell arrow map reinforces this view: within meristem clusters and at interfaces between clusters, arrows are often longer and consistently oriented, indicating faster transcriptional change and transitional populations migrating toward neighboring tissues; by contrast, arrows inside cortex, endodermis, stele, and root-cap compartments are typically shorter and less variable, consistent with more stable or terminal states. These directional flows aligned with the MST-based lineage tree (**Figure 3A**), confirming that RNA velocity captured biologically meaningful progression among root cell states. We observed similar trends in maize as well with streamlines radiated from the quiescent center (QC) toward Epidermis (Z5) and Cortex (Z1), with coherent arrows extending across the QC–outer boundary. Some flows converged inward from Cortex/Endodermis (Z6), BCs (Z9), and Endodermis (Z10) back toward the QC. Endodermis (Z10) and Stele/Xylem (Z3) were largely velocity-poor, consistent with near-steady kinetics (**Figure 3B, middle**). In sorghum, epidermis (S14) acted as a local source, showing outward-oriented streamlines across Cortex (S3), Cortex–Endodermis (S4), and Endodermis (S8). QC-related clusters (S9, S10, S13) showed outgoing flows, while QC (S7) behaved as a sink with short, disordered arrows. Paths toward Columella (S2/S6), Root cap (S11), and BCs (S12) formed clear terminal trajectories (**Figure 3B, right**).

Latent time, which estimates a cell’s internal time with respect to the starting cellular state (QC in our case), provided a one-dimensional, biologically interpretable ordering of cells in each species. In Medicago, cluster-level aggregation of latent time recapitulated the lineage-like tree trajectory: Cortex/Epidermis → Meristem → Cortex → Root cap/Columella → Stele/Endodermis → Endodermis → Root cap/BCs → Stele Phloem (means 0.44, 0.55, 0.66, 0.74–0.75, 0.79, 0.80, 0.81, respectively) **(Figure 3C, Medicago; Supplemental data set 3).** For maize and sorghum, as automatic root detection was uncertain, we initialized latent time with QC-annotated clusters to set the origin and then computed latent time from those roots. In maize, the ordering localized QC2 earliest and increased through Epidermis → Cortex/Endodermis → QC1 → BCs → Stele/Xylem → Endodermis → Cortex → Columella/Cap (cluster means: 0.23, 0.28, 0.33, 0.41, 0.42, 0.51, 0.54, 0.56, 0.67) (**Figure 3C, Maize; Supplemental data set 3)**. In sorghum, early values mapped to Epidermis and QC, increasing through Endodermis and Cortex and reaching the highest values in BCs, Stele, and Root Cap (cluster means: Epidermis 0.26 → QC 0.37 → Endodermis 0.42 → Cortex/Endodermis 0.45 → Columella 0.51 → Cortex 0.52 → BCs 0.61 → Stele 0.63 → Root Cap 0.86) **(Figure 3C, Sorghum; Supplemental data set 3).**

Ordering velocity-supported marker genes by scVelo latent time revealed a clear sequence of transcriptional waves across the root program **(Figure 3D, Supplemental Figure 12)**. In Medicago, at the start of the latent-time axis, expression peaks for epidermis/cortex markers (e.g., *MtrunA17_Chr8g0374261*, *MtrunA17_Chr5g0423841*, and *MtrunA17_Chr7g0243081)*, followed closely by a surge in meristem-associated regulators (*MtPLT4*; *MtBBM*, *MtrunA17_Chr7g0243201*, *MtrunA17_Chr7g0263701*, *MtPLT1*, *MtPIN4*, and an *AtE2Fe-like* factor). This pattern suggests these genes act early in the program captured by the velocity model, i.e., during the initial steps of epidermal/cortical specification and meristem activity. As latent time increases, endodermis markers (*MtrunA17_Chr1g0212251*, *MtRIP4*, *MtrunA17_Chr4g0032331*, *MtrunA17_Chr4g0042551*, *MtRbohA*, *MtrunA17_Chr7g0274621*) show rising expression, consistent with progression toward inner tissue identities. Root-hair/border-cell genes (*MtLAT52/POE_5*, *MtPrx02*, *MtRopGEF12*, *MtLAT52/POE_16*, *MtEXP7*) exhibit transient activation, matching their specialized roles at the root surface, and the trajectory culminates with increased expression of stele/vascular markers (*MtNPF1.6*, *MtCIF1*, *MtHB8*, *MtnsLTP127*, *MtMYB122*) **(Figure 3D, Medicago).**

Similarly, in maize, QC (e.g. *Zm00001eb434370*, *Zm00001eb068280*) and epidermis markers (e.g. *Zm00001eb088470*, *Zm00001eb310700*) show elevated expression at the beginning of latent time trajectory, followed by cortex (e.g. *Zm00001eb169520*, *Zm00001eb197870)* and endodermis markers (e.g., *Zm00001eb289210*) and culminating with BC (e.g. *Zm00001eb260350*, *Zm00001eb117180)* and stele markers (e.g., *Zm00001eb246860*, *Zm00001eb242820),* (**Figure 3D, Maize)**. Sorghum markers too showed a similar pattern when ordered by latent time with epidermis (*SbiRTX430.01G361600*) and QC (*SbiRTX430.02G200000, SbiRTX430.05G071000*) high at earlier stages, followed by endodermis (e.g., *SbiRTX430.02G182200, SbiRTX430.08G122700*) and cortex (*SbiRTX430.01G508300*). The trajectory ultimately culminated in increased expression in the columella (*SbiRTX430.02G218000, SbiRTX430.09G073100*), BCs (*SbiRTX430.06G015600*), and cap (*SbiRTX430.01G129500*) **(Figure 3D, Sorghum),** showing overall consistency with the other two species.

Finally, we examined the phase portraits of cell type-specific markers, including the novel ones we predicted and validated in this study. Phase portraits are used to examine the transcriptional up or down-regulation of individual genes based on the relative unspliced (y-axis) vs spliced (x-axis) reads per cell, compared to their steady-state relationship. For each cell type, an upregulated gene has an excess of unspliced reads, while downregulated genes have an excess of spliced reads. Orthologous markers in maize and sorghum showed conserved pattern profiles, including those validated experimentally. The endodermis orthologous pair, maize *Zm00001eb005460* (validated in this study) and sorghum *SbiRTX430.01G515100*, displayed clean induction portraits and positive velocities within expressing cells. Root-cap orthologs were also concordant: sorghum SbiRTX430.04G115800 (validated in this study) and maize orthologs, *Zm00001eb012020*, *Zm00001eb148030*, and *Zm00001eb367780,* showed similar induction profiles and spatial localization, with species-specific amplitude differences. Similarly, the validated sorghum BC marker *SbiRTX430.05G002900* and the validated root cap maize marker *Zm00001eb163560* both showed characteristic induction profiles localized to those tissues (**Figure 3E**).

Together, these analyses reveal conserved transcriptional dynamics for endodermis and root-cap/border-cell genes across Medicago, maize, and sorghum. Despite minor differences in kinetic amplitude and coherence, the underlying developmental trajectories and regulatory programs (as we show next) appear conserved across species.

### Gene regulatory networks and modules underlying mucilage production in roots

To gain insight into regulatory mechanisms underlying mucilage production, we applied the GRN inference algorithms, Modular Regulatory Network Learning with Per-Gene Information (MERLIN) framework (Roy et al. 2013), and single-cell Multi-task Network Inference (scMTNI) (Zhang et al. 2023) to the species-specific gene expression matrices. MERLIN-P enables inference of regulatory interactions at single-gene resolution and at the module level, while scMTNI (discussed in more detail next) provides cell type-specific GRNs.

We first computationally validated the inferred GRNs using multiple metrics, including area under the precision-recall curve (AUPR), early precision, recall, and F-score, using gold-standard GRNs defined based on accessibility and chromatin-mark-based datasets **(Methods, Supplemental Table 2)**. For each dataset, we defined five regulatory windows centered on the transcription start site (TSS; ±1–5 kb) to create five gold-standard GRNs. We compared the metrics for the MERLIN GRNs with those from randomized edge-shuffled networks used as null baselines for fold-enrichment estimation.

MERLIN achieved high fold enrichments for F-score (6.9-11.6-fold), and predictable TFs (14-17-fold) across the different gold standards (**Supplemental Figures 13, 14, 17 and 18, Methods**) in Medicago. The AUPRs were not as informative with fold enrichments ∼1.0 due to the large ATAC-seq gold standard, which is also evident by the low recall at different network sizes (**Supplemental Figure 17**). We observed a similar behavior in maize and sorghum, with high fold enrichment of F-score (9.3-10.6-fold in maize, 7.5-8.4-fold in sorghum, **Supplemental Figures 15 - 17**) and predictable TFs (14-19-fold in maize, 2-2.7-fold in sorghum, **Supplemental Figures 19**), but uninformative AUPRs. The relatively lower fold enrichment in sorghum likely reflects our use of histone modification ChIP-seq data rather than ATAC-seq, which is more informative of physical TF binding.

After computationally validating the GRNs, we interpreted them in a species-specific manner. In Medicago, MERLIN inferred a network comprising 14,853 regulatory edges connecting 1,118 of the initial 2,435 candidate regulators to 6,483 target genes, with all interactions supported by at least 80% confidence (**Figure 4A**). The two most prominent regulators were MtrunA17_Chr5g0393721 and the histone demethylase MtJMJ20 (MtrunA17_Chr3g0132281), which regulated 267 and 197 target genes, respectively (**Supplemental Figure 20A**). MERLIN identified 52 regulatory modules, each containing ≥5 genes and together encompassing 1,013 genes (**Supplemental Figure 20B; Supplemental data set 4)**. Modules ranged from 5 to 145 genes (mean = 19). On average, six regulators were significantly enriched per module (range: 0–33), confirming the modular structure of the inferred network. GO enrichment analysis revealed that 27 of the 52 modules were enriched for at least one biological process **(Supplemental data set 4).**

**Figure 4.**
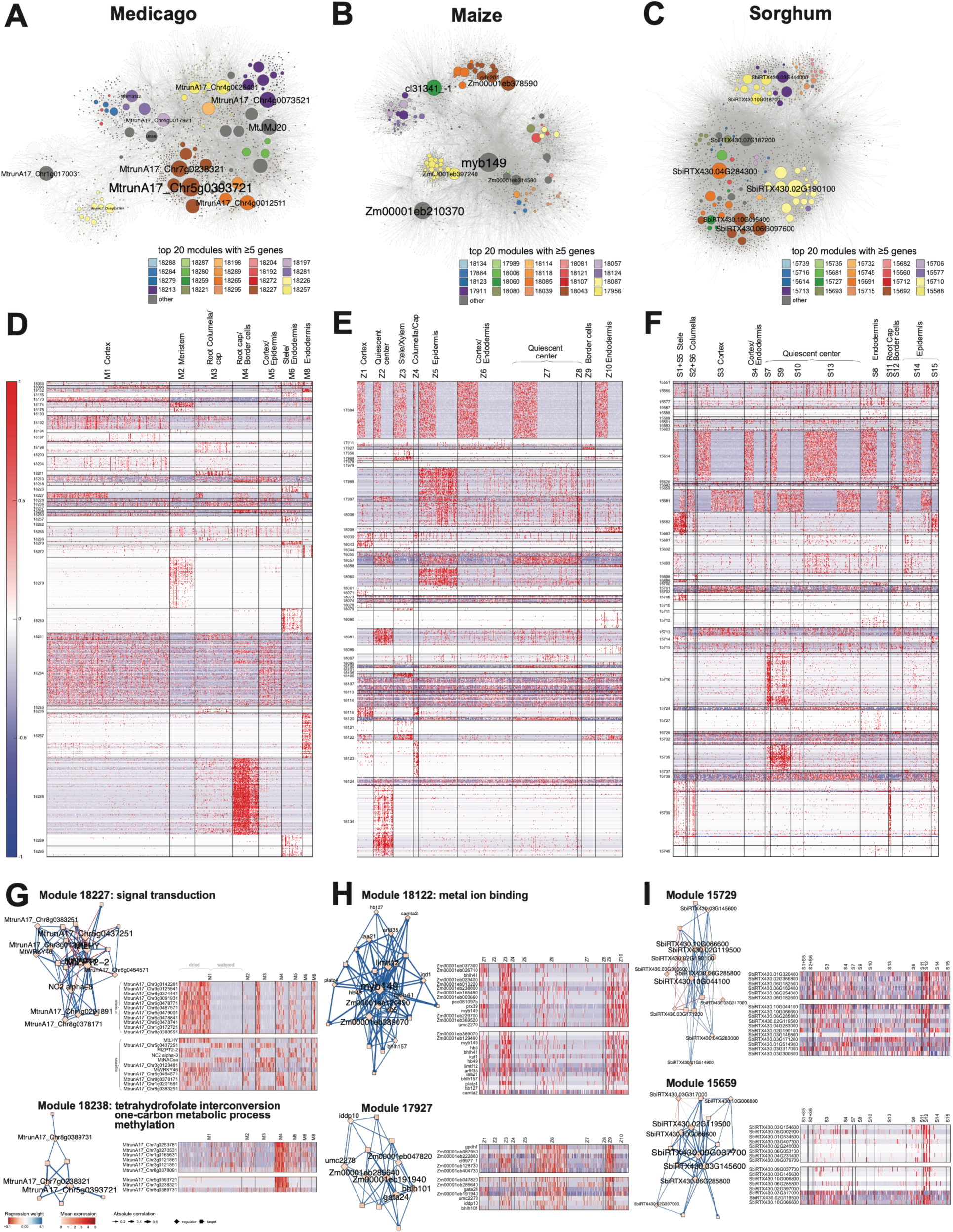
Gene regulatory networks inferred by MERLIN for Medicago, maize, and sorghum. **A–C)** Force-directed layouts of the inferred networks for **A)** Medicago, **B)** maize, and **C)** sorghum. Genes belonging to the same module are shown with the same color for the top 20 modules containing at least five genes. Node size is proportional to outdegree, and regulator names are shown for the largest nodes. Network visualization was performed using Cytoscape (v3.10.3). **D–F)** Expression patterns of MERLIN-predicted modules containing at least five genes across cell clusters for **D)** Medicago, **E)** maize, and **F)** sorghum. Each row represents the expression profile of a gene within the module, and columns represent individual cells grouped by cell cluster and ordered by sample (dry followed by wet). **G–I)** Detailed views of selected MERLIN modules showing regulator-target interactions, gene expression heatmaps, and associated Gene Ontology (GO) terms when available. Diamond-shaped nodes represent regulators, and square nodes represent gene targets. Node color indicated mean gene expression level. Edge color and width correspond to the learned regression weights and regulator-target correlation, respectively. For each module, the heatmap is divided into two sections: expression of module genes (top) and expression of their regulators (bottom). In Medicago **G)**, module 18227 displays elevated expression in dry-sample cells, whereas module 18238 is upregulated in cluster associated with root cap/BCs. For maize **H)** and sorghum **I)**, the selected modules show increased expression primarily in BC-associated clusters.

In maize, the final 0.8-confidence network included 19,987 regulatory edges connecting 1,160 regulators to 7,088 targets (**Figure 4B)**. The most connected regulators were Zm00001eb417620 and Zm00001eb191940, regulating 638 and 561 genes, respectively (**Supplemental Figure 20C**). MERLIN identified 51 modules, each with at least five genes and collectively spanning 1,184 genes (**Figure 4E**). Modules ranged from 5 to 172 genes (mean = 23), and on average, nine regulators were enriched per module (range: 0–29) (**Supplemental Figure 20D, Supplemental data set 4)**. Thirty-three of these modules showed enrichment for at least one GO term, indicating coherent biological organization **(Supplemental data set 4)**.

In sorghum, the final network contained 31,295 edges connecting 1,007 regulators to 8,437 targets (**Figure 4C**). The two top regulators, SbiRTX430.04G355300 and SbiRTX430.02G190100, regulated 1,651 and 684 targets, respectively (**Supplemental Figure 20E**). MERLIN identified 46 modules, each containing more than five genes and spanning 1,196 genes in total (**Figure 4F**). Module sizes ranged from 5 to 162 genes (mean = 26), with 14 enriched regulators per module on average (range: 0–50) (**Supplemental Figure 20F; Supplemental data set 4**). Twenty-nine of the 46 modules were significantly enriched for at least one GO term, confirming the functional structure of the inferred sorghum GRN **(Supplemental data set 4)**.

Across all three species, MERLIN modules captured clear transcriptional responses between dry and wet conditions across most tissue-specific clusters. For Medicago, Module 18227 showed strong induction under dry conditions and repression under wet conditions (**Figure 4G**). This pattern was consistent across clusters except in Cluster 2 (Meristem). The corresponding regulators exhibited parallel expression changes, and genes in this module were enriched for signal transduction. In contrast, Module 18096 showed the opposite trend, with repression under dry and upregulation under watered conditions. Similar responsive patterns were observed in sorghum (**Figure 4F**). Module 15614 was upregulated under dry conditions and enriched for mRNA processing, RNA, and nucleotide binding, whereas Module 15681 showed higher expression under watered conditions and was enriched for RNA binding, translation, and ribosomal structure. Finally, in maize, Module 17884 exhibited strong induction under dry conditions, with enrichment for chromatin organization, RNA helicase activity, translation, and vesicle trafficking functions (**Figure 4E**). In addition to stress responses, several modules showed tissue-specific expression corresponding to root cap and BC clusters. In Medicago, Modules 18211 and 18221 were upregulated in Cluster 3 (Root cap/Columella) and enriched for protein phosphorylation, defense response, and oxidation–reduction processes. Modules 18238 and 18288 were upregulated in Cluster 4 (Root cap/BCs) and enriched for cell wall modification and ion transport (**Figures 4D, G**). In maize, Module 18118 showed upregulation in Cluster 4 (Columella/Cap) and enrichment for transmembrane transport, while Modules 17927 and 18122 were upregulated in Cluster 9 (BCs) and enriched for plasma membrane, extracellular region, and oxidoreductase activity (**Figure 4H**). In sorghum, Modules 15659 and 15729 were upregulated in Cluster 12 (BCs) (**Figure 4I**).

Altogether, these results show that MERLIN effectively captured both species-specific and conserved regulatory patterns, revealing coordinated water responses and cell-type–specific regulatory programs in Medicago, maize, and sorghum roots.

### Cell type-specific GRN inference identifies key subnetworks rewired between BC and non-BC populations

While MERLIN outputs a single global GRN and modules across all cell types, examining how GRNs change across cell types in each species can prioritize key regulatory mechanisms underlying BC development. To this end, we applied single-cell Multi-Task Network Inference (scMTNI), a cell type-specific GRN inference algorithm that integrates cell type-specific scRNA-seq datasets with a cell lineage structure (Zhang et al. 2023) to each species’ clustered data. As the input tree, we used our refined cell lineage tree from previously defined cluster-averaged expression profiles (**Figure 5A**, **Methods**).

**Figure 5.**
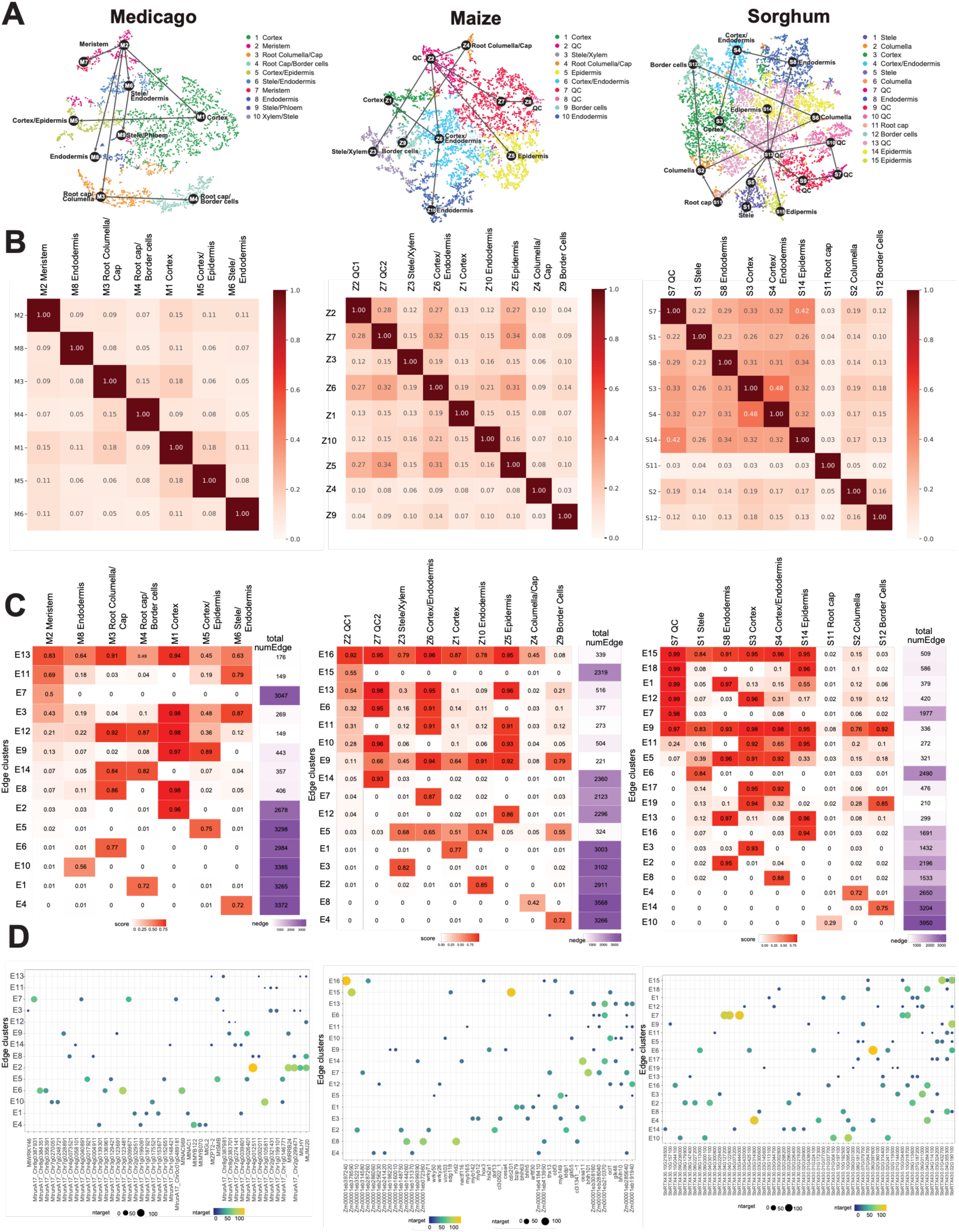
Cell-type–specific network inference for medicago, maize and sorghum using scMTNI. **(A)** tSNE visualization of cell clusters in Medicago, maize, and sorghum, with annotated cell types. Colors indicate distinct cell clusters identified from single-cell data. Similar cell types were merged in the final lineage tree shown in Figure 3A (schematic lineage trees). **(B)** Pairwise F-score comparison between inferred regulatory networks of cell clusters in each species. Color intensity indicates the degree of similarity between cell types across species **(C)** k-means edge clustering analysis of top 4k edges from inferred gene regulatory networks. Heatmaps show the mean edge confidence scores of edge clusters across cell clusters in each species. Each row corresponds to an edge cluster (E1–Ek), and each column corresponds to a cell cluster. Shades of purple on the right indicate the total number of edges within each edge cluster. The number of clusters used for k-means is k = 14 for Medicago, k = 16 for maize, and k = 19 for sorghum. **(D)** Top five regulators associated with each edge cluster. Circle size and color correspond to the number of target genes regulated by that transcription factor.

Similar to MERLIN, we first computationally validated the scMTNI GRNs using similar metrics computed for each cell cluster individually. These scMTNI networks showed similar trends, with inferred networks enriched by F-score, across all three species (**Supplemental Figures 13-17**), but varied across cell clusters. In Medicago, Cortex (M1) had the highest fold enrichment, followed by Cortex/Epidermis (M5) and Stele/Endodermis (M6), with F-score enrichment of ∼3–6-fold, while Root cap/BCs (M4) and Endodermis alone (M8) had lower enrichment (1.4-2-fold) (**Supplemental Figures 13, 14, 17**). We observed a similar behavior in maize, with scMTNI networks showing higher fold enrichment in Cortex (Z6), QC2 (Z7), and Epidermis (Z5). Here too, Columella/Cap (Z4) and BCs (Z9) were weaker in fold enrichment (∼1.7–2.0-fold) (**Supplemental Figures 15 and 17**). In sorghum, scMTNI networks for Endodermis (S8) had the highest fold enrichment (F-score enrichment ∼6-fold, precision 0.206–0.230), while Root cap (S11) and BCs (S12) were near baseline (1-1.2-fold) for most gold standards (**Supplemental Figures 16 and 17**). When using predictable TFs as a metric, Medicago and maize scMTNI GRNs for Cortex/Epidermis and QC2 (M1, Z6, Z7), were enriched (**Supplemental Figure 17-19**). As the same TF might have targets in multiple cell types, we also computed predictable TFs on the merged scMTNI GRNs and found substantially higher fold enrichment in Medicago and maize (**Supplemental Figure 17**). Taken together, these computational validation results indicate that MERLIN and scMTNI successfully captured regulatory relationships supported by physical binding data from ATAC-seq or histone ChIP-seq data.

Based on the distribution of edge confidence scores **(Supplemental data set 5)**, we selected the top 4,000 edges for each species using thresholds of 60% for Medicago, 70% for maize, and 80% for sorghum, yielding approximately 4,000 high-confidence regulatory interactions in each case. In Medicago, these edges linked 531 regulators and 2,589 target genes in M1 (cortex) to 1,116 regulators and 3,169 target genes in M6 (stele/endodermis). In maize, scMTNI GRNs linked 570 (Z4: columella/cap) and 1,095 (Z9: BCs) regulators to target genes ranging from 2,476 (Z6: cortex/endodermis) to 3,185 (Z9: BCs). In sorghum, regulator counts were lowest in S7 (quiescent center; 297) and highest in S12 (BCs; 970), while target genes ranged from 2,231 in S3: cortex to 3,142 in S12: BCs. Pairwise F-score comparisons of the inferred networks showed that the majority of TF–target interactions are cell-type specific (F-scores < 0.3) for most cell type-specific GRNS across all three species, but the extent of cell type-specificity varied between species **(Figure 5B)**. scMTNI GRNs were most dissimilar in Medicago, followed by maize and sorghum. Similar cell types tended to have more similar GRNs (e.g., Cortex, Epidermis, and Endodermis) across all three species. Furthermore, in maize and sorghum, QC cells were closer to Cortex, Epidermis and Endodermis cell types compared to Columella or BCs, reflecting their spatial organization.

We used the scMTNI GRNs to examine lineage-resolved regulatory dynamics using two complementary approaches: k-means edge clustering to identify dynamic regulators within subnetworks, and Latent Dirichlet Allocation (LDA) topic modeling to detect regulator rewiring and define dynamic subnetworks across cell clusters. The k-means analysis focuses on more cell-type-specific subnetworks for individual cell type/clusters, while the LDA analysis identifies subnetworks for a subset of cell types/clusters.

In Medicago, k-means clustering partitioned the edge confidence matrix across the different cell clusters into 14 patterns that captured both shared and cluster-specific regulatory relationships (**Figure 5C, left**). The largest edge clusters corresponded to specific cell types, including E7:M2 (meristem), E2:M1 (cortex), E5:M5 (cortex/epidermis), E6:M3 (root columella/cap), E10:M8 (endodermis), E1:M4 (Root cap/BCs), and E4:M6 (stele/endodermis). Several edge clusters were also shared across multiple lineages, including E14, which was shared between M3 (root columella/cap) and M4 (Root cap/BCs), and E9, which was shared among a larger set of clusters, including cortex and epidermis. In contrast, small edge clusters were shared across multiple cell types. To better understand these clusters, we examined the top regulators associated with each edge cluster (**Figure 5D, left**). Edge cluster E1, which is specific to BCs, has MtNAC1 as one of the top regulators, and D’haeseleer et al. (2011) showed that its post-transcriptional repression by miR164 leads to reduced protein levels and a decrease in nodule number, indicating an indirect role in nodule development. E2, corresponding to cortex cell types, has MtrunA17_Chr4g0012511 (137 targets), MtRRB24 (98), MtrunA17_Chr2g0299471 (97), MtLHY (63), and MtJMJ20 (80) as top regulators. We also examined the regulators linked to edge clusters shared across multiple lineages and found that MtLHY and MtJMJ20 were frequently associated with these clusters. MtLHY regulates circadian rhythms and enhances salt stress tolerance in Medicago (Lu et al. 2023), while MtJMJ20 is associated with chromatin modification and gene regulation, contributing to developmental processes through histone demethylation (Lopez et al. 2022). Edge cluster E14, shared between the root columella/cap (M3) and Root cap/BCs (M4) lineages, was dominated by *MtrunA17_Chr1g0148421* and *MtZPT2-2*, with 13 and 14 targets, respectively; both were experimentally validated by RNAi knockdown, where silencing *MtrunA17_Chr1g0148421* significantly reduced BC numbers (as described later).

To identify additional shared and cell type-specific subnetworks, we performed LDA analysis with the number of latent topics set to 10 (**Figure 6A, right)**, each topic corresponding to a GRN subnetwork that could reflect a pathway conserved or rewired across the cell clusters (**Supplemental Figures 21 and 22**; **Figure 6A**). Nearly all topics, except topic 6 (**Supplemental Figure 22A)**, showed rewiring of GRN topology across cell clusters, indicated by changes in the degree of regulators across cell clusters in the topic of interest. Several of these are associated with root cap, BC processes and BC lineage and enriched for processes related to development, hormone signaling and root morphology (**Supplemental Figure. 23**). Topic 5 along the BC lineage (M2–M4) identified *MtrunA17_Chr1g0201891, MtrunA17_Chr8g0378171,* and *MtZPT2-2* as regulatory hubs, while targets of MtLHY were lost in M4 and M5 (**Figure. 6A, left**). Topic 5 was enriched for signaling processes and included several regulators such as MtLBD18 with known roles in root development. In contrast, Topic 1 included regulators like MtPLT2, MtSMB, and *MtrunA17_Chr8g0339091* (**Supplemental Figure. 21B**) and was enriched for developmental processes such as root hair cell differentiation and cell wall organization. Other topics of interest were enriched for relevant GO processes and regulated by TFs with potential involvement in root morphogenesis, including topics 3 predicted to be regulated by MtBELL and MtFIE and involved in developmental processes (**Figure 6 (right), Supplemental Figure. 21D**).

**Figure 6.**
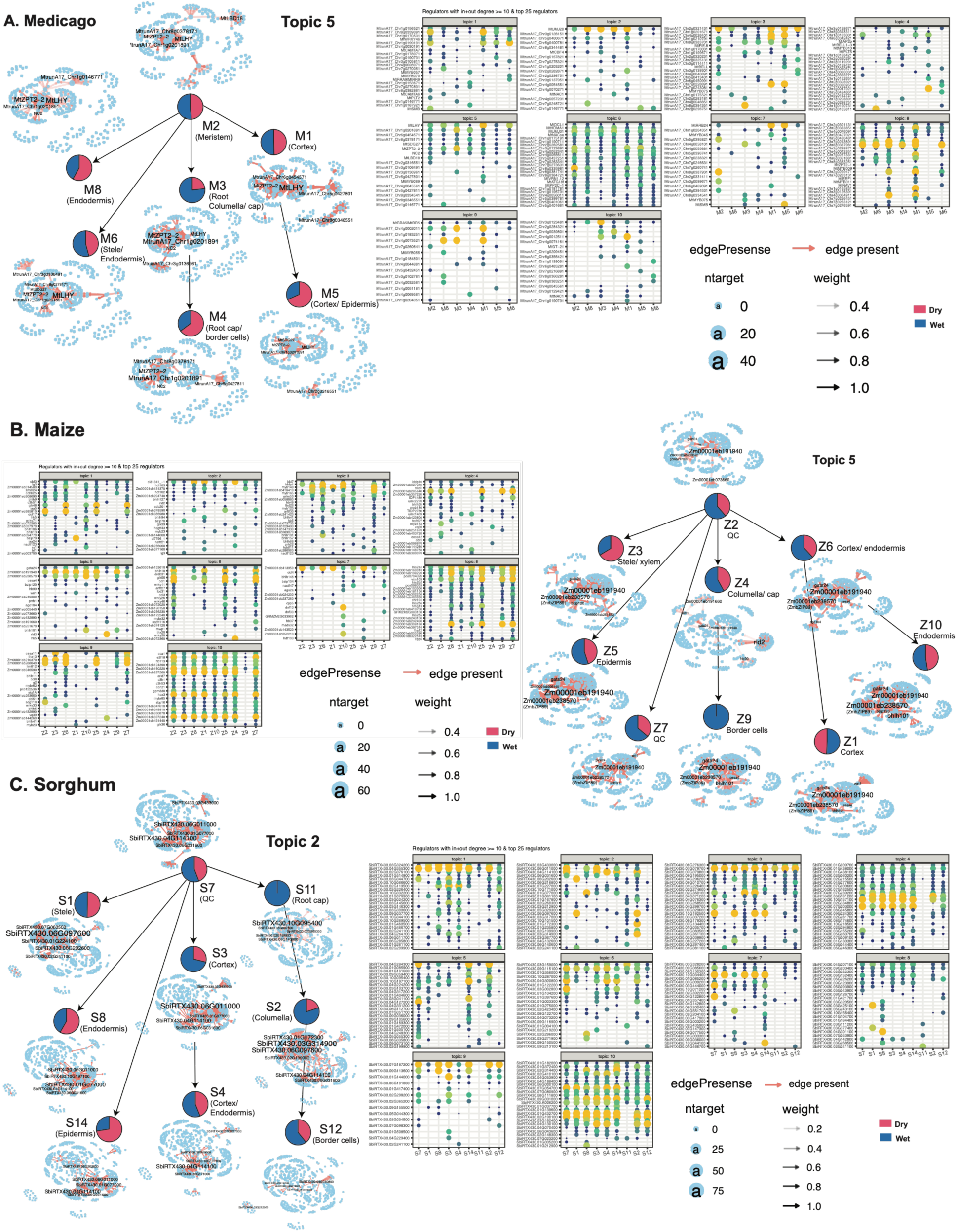
LDA topic analysis of inferred gene regulatory networks across three species: **A)** LDA topic 5 networks in Medicago along the cell lineage. In each cell cluster, nodes represent regulators, and edges are shown in red. Nodes labeled in the network represent regulators with a degree greater than 10. Plots on the right in each species show the cell cluster–specific top regulators that have at least 10 targets and are among the top 25 regulators in the topic. Circle size and yellow color intensity increase with the number of targets regulated by the regulator. **B)** Maize LDA topic 5 and **C)** Sorghum LDA topic 2 networks.

For the maize k-means edge clustering analysis, k = 16 provided high SI values and moderate cluster sizes **(Supplemental Figure 24, middle),** with the highest number of edges (∼92%) being specific to a single cell cluster (e.g., E15, E14, E7, E12, E1, E3, E2, E8 and E4), whereas the smaller edge clusters contain edges shared across multiple cell clusters (e.g., E16, E9, and E5) **(Figure 5C, middle).** Cluster 9 corresponds to the BC, where edge cluster E4 is specifically expressed. Top regulators associated with E4 are Zm00001eb286060 (29 targets), Zm00001eb241420 (23), Zm00001eb418410 (29), Zm00001eb131370 (32), and Zm00001eb285640 (27) **(Figure 5D, middle).** Edge cluster E8, specific to the columella/root cap cell cluster Z4, had several regulatory hubs, including rld2 (101 targets), Zm00001eb072360 (91), and Zm00001eb144260 (82). Of these, rld2 has been shown to be a key regulator of root development and morphogenesis in maize (Gautam et al. 2021). The edge clusters E10 shared between QC and epidermis had ocl1, glk48, and bhlh10 as top regulators. E13, also shared with QC and epidermis and additionally Cortex, included glk48 and other hubs such as Zm00001eb210370, Zm00001eb285640, and Zm00001eb191940. Finally, edge cluster E16, which included nearly all cell clusters (least for BCs), had obf3, Zm00001eb397240, and Zm00001eb287260 as top regulators. In contrast, the edge clusters (E9, E5), which had more shared edges from BCs, had his2a1 and cesa4 as top regulators (E9) and myb149 (E5). Of these, ocl1 has been specifically associated with epidermis in maize, while the others, like obf3, glk48, bhlh10, and myb149, are broad developmental regulators, which is consistent with their association across multiple cell types (Javelle et al. 2010; Chen et al. 2016; Zhang et al. 2018; Wei et al. 2012).

Similar to Medicago, we next applied LDA topic modeling analysis to examine subnetwork-level rewiring across the maize cell clusters (**Figure 6B)**. Except for topic 10 (**Supplemental Figure. 26F**), all other topics exhibited regulator rewiring based on the change in their degree across cell types. GO enrichment analysis identified several topics relevant for BC morphology and function for the BC cluster (Z9). This included topic 1, enriched for ABA signaling, topics 3 and 5 for auxin signaling and topic 6 for vesicle transport (**Figure 6B**, **Supplemental Figure 27)**. The regulators associated with these topics were involved in cell wall synthesis (topic5: cesa4), hormone signaling (topic1: obf3, iaa24 and iaa5) as well as TF families implicated in environmental stress response (topic5: BZIP family (bzip120, bzip89/Zm00001eb238570) (Ma et al. 2018), and BHLH family (bhlh101) (Heim et al. 2003)), **(Supplemental Figure 25B, Supplemental data set 6),** further supporting Z9 and its associated GRN to be involved in BC morphology, stress response and potentially mucilage production.

In sorghum, the k-means analysis, *k*=19 edge clusters were optimal based on SI (**Supplemental Figure 24**). Like Medicago and maize, the majority of the edges (84.73%) came from edge clusters that correspond to individual cell clusters (E7, E6, E16, E3, E2, E8, E4, E14, and E10), with E14 being exclusive to the BC cluster S12 (**Figure 5C, right**). The remaining edge clusters were shared across multiple cell clusters, of which E9 and E19 included edges from S12. We interpreted edge clusters E14, E9, and E19 based on available maize orthologs **(Supplemental data set 6)**. E19, which spanned Cortex and Columella in addition to BCs, included top regulators involved in meristem identity (knox5, Ig3) and secondary cell wall biosynthesis (myb52, myb149, mapped to SbiRTX430.03G145600, and cesa6, cesa8, mapped to SbiRTX430.02G119500). The top regulators of E9 tended to involve more general regulatory processes, consistent with their edges being conserved across more cell clusters. Finally, E14’s top regulators included maize orthologs for *knox5* and *Ig3* in addition to regulators Iddp1 and Iddp7 (mapped to SbiRTX430.07G187200) (**Figure 5D, right**), which are involved in developmental transitions though have not been associated with BCs.

Similar to Medicago and maize, we did LDA analysis with k=10 topics in sorghum and identified subnetworks associated with BC formation regulators (**Figure 6C, right**) and GO processes (**Supplemental Figure 28**). In particular, topics 2, 4, and 10 were associated with hormone signaling regulators (WRKY50, EIL4), root cap-related functions (NACTF11, 29, or 71), or enriched for GO processes such as cell wall biogenesis and signaling response (**Supplemental Figures 29 and 30**), indicative of their BC association. Specifically, topic 2’s NACTF11 (sorghum gene SbiRTX430.01G404600) is known to be a root cap regulator and exhibited a large number of connections in the root cap cell cluster S11.

Taken together, these results identified several developmental regulators for the sorghum cell clusters that are plausible candidates for further examination of BC formation.

### Cross-species analysis of inferred GRNs identifies conserved and species-specific mucilage GRN components

To assess conservation and divergence across species, we compared GRNs inferred with MERLIN with cell-type-specific GRNs inferred with scMTNI for matched cell types (**Supplemental Figure 31**). We quantified the similarity and differences using network comparison metrics, namely precision, recall, and F-score. MERLIN-inferred GRNs exhibited an F-score ranging from 2-12%, with the highest similarity for sorghum and maize, followed by their comparisons to Medicago, which is consistent with sorghum and maize being evolutionarily closer to each other than either is to Medicago. The conserved GRN between sorghum and maize had 296 regulator OGs and 1151 target OGs, while the conserved core across all three species included 28 regulators and 78 targets. The top regulators here included several generic developmental regulators such as gata4, secondary cell wall-related functions such as myb128, myb149, and some GRAS-family regulators (gras42, gras47, gras88) that have been implicated in root patterning and development (Jaiswal et al. 2022)

To gain insight into the conservation patterns from a cell-type-specific point of view, we compared all the scMTNI-inferred GRNs for one species to all scMTNI-inferred GRNs for another species. GRNs from different cell types in sorghum and maize were more similar to each other than to Medicago. Most notably, QC, epidermis, and cortex/endodermis had the greatest conservation, although the overall conservation was low (F-score 0.07-0.09). The conserved GRN between sorghum and maize included 7-10 TFs groups in these cell types. Between the three species, the cell type-specific GRNs did not exhibit a clear pattern except for Medicago’s meristem cell cluster, which had the largest overlap with the QC cell cluster in sorghum and maize and included 49 TFs. The top regulators between maize and sorghum QC included cdc201 and gras42, which we discovered before, but also ago1a, which is a generic developmental regulator of microRNAs, developmental regulators like idd7 and iddp1, and chromatin remodelers (hist2a, sgd11, nfd106) that indicate regulatory roles in cell fate and cell state transitions.

Finally, we intersected these regulators with our scMTNI LDA topics and found several interesting regulators. One such regulator set includes iddp1 (Zm00001eb039810, maize; topic3, **Supplemental Figure 25D**) and its sorghum ortholog SbiRTX430.07G187200 (sorghum; topic9, **Supplemental Figure 30E** and MtrunA17_Chr1g0163001 (Medicago; topic4) (**Supplemental Figure 21E**). Iddp1 is associated with epidermis, cortex/epidermis and QC-related clusters and shared several targets between sorghum and maize. In Medicago, this regulator was most connected in cortex/epidermis-associated clusters (M5) and meristem/cortex (M1, M2), which are also the two cell types exhibiting conservation across the three species. In maize, *iddp1* had targets in endodermis (Z10), cortex/endodermis (Z6), QC and root cap clusters (Z2, Z3, Z7, Z9), while in sorghum, its ortholog SbiRTX430.07G187200 had the highest number of targets in QC (S7), BCs (S12), and epidermis (S14), indicating a conserved epidermis-QC cell type across the three species.

Taken together, these comparative analyses show the dynamic nature of regulatory networks and inform conserved and diverged GRN circuits that could be probed further with experimental perturbations.

### Experimental validation of gene regulators associated with mucilage production

We used our network dynamics and evolutionary analyses on the MERLIN and scMTNI-inferred GRNs to identify candidate regulators important for BCs and mucilage production and tested them with RNAi knockdown. We tested these regulators in Medicago to assess their tractability, selecting candidates based on their reported regulatory roles and, where applicable, the presence of orthologs in maize and sorghum (**Supplemental Table 3**). We generated RNAi knockdown lines targeting five transcription factors from diverse TF families: Homeobox-WOX (MtrunA17_Chr5g0400781), TUBBY (MtrunA17_Chr1g0201891), MYB (MtrunA17_Chr3g0111941), Krüppel-like (MtrunA17_Chr1g0207451), and C2H2 zinc finger (MtrunA17_Chr1g0148421).

The Homeobox-WOX TF MtrunA17_Chr5g0400781 was associated with MERLIN module Module18181 and scMTNI topic 2 (**Figure 6A**; right). TUBBY (MtrunA17_Chr1g0201891) was a top regulator in topic 5 of Medicago (**Figure 6A**; left), and its ortholog Zm00001eb202420 (tubtf3) in maize was rewired in topic 7 within cluster 9, which corresponds to the BC type, and it regulates seven targets. The putative C2H2 zinc finger protein MtrunA17_Chr1g0148421 was a dominant TF of the edge cluster E14, which had many interactions in BCs and Columella. This also shares orthology with SbiRTX430.03G300600 in sorghum and Zm00001eb154580 (umc1489) in maize. In sorghum, this ortholog emerged as a top regulator within k-means cluster E8, dominated by cortex/endodermis (S4, **Figure 5C, D** right), and exhibited rewiring across cell clusters with high connectivity in S4 and S8 (endodermis) based on LDA analysis (topic 1, **Supplemental Figure 29B**), suggesting an association with the cortex/endodermis cell type. In maize, the ortholog displayed rewiring in topic 4 of cluster 10 (endodermis, **Figure 6B**, Right) and topic 9 of cluster 4 (columella/Root cap), indicating functional diversification across cell types. These findings show the dynamic nature of regulatory networks, where conserved genes adapt to species-specific developmental contexts.

Phase portraits of orthologs selected for functional validation by RNAi knockdown supported conserved induction kinetics with species-specific variation. In maize, *Zm00001eb025650* and *Zm00001eb154580* (orthologs of *MtrunA17_Chr5g0400781* and *MtrunA17_Chr1g0148421*) showed positive velocities and consistent induction geometries, whereas sorghum SbiRTX430.03G396900 (ortholog of MtrunA17_Chr1g0201891) exhibited broader residuals. Endodermis orthologs Zm00001eb201250 and SbiRTX430.05G061900 (orthologs of MtrunA17_Chr2g0321181) showed short, fitted segments followed by steady-state stabilization (**Figure 3E**).

Since BCs are essential for mucilage production (Pankievicz et al. 2022), we quantified their number as a proxy for mucilage output. Roots were hydrated to release BCs, and counts from RNAi lines were compared to an empty vector (EV) control expressing only the transformation marker. We used an automated cell counter calibrated specifically to detect BCs while excluding other cellular debris. All RNAi lines showed a general trend toward reduced BC numbers, but only three lines, Homeobox-WOX (MtrunA17_Chr5g0400781), TUBBY, and MtrunA17_Chr1g0148421, showed statistically significant reductions compared with EV (**Figure 7A**). Microscopic analysis of root tips confirmed fewer BCs in knockdown lines, and interestingly, some appeared collapsed and deformed (**Figure 7B**). We assessed this using cell viability with live/dead staining with trypan blue. Using the same automated cell counting system, we quantified the proportion of dead and live cells. The control EV line exhibited approximately 75% live and 25% dead cells, while only MtrunA17_Chr1g0148421 showed a significant drop in viability, with live cells reduced by around 50% (**Figure 7C**).

**Figure 7.**
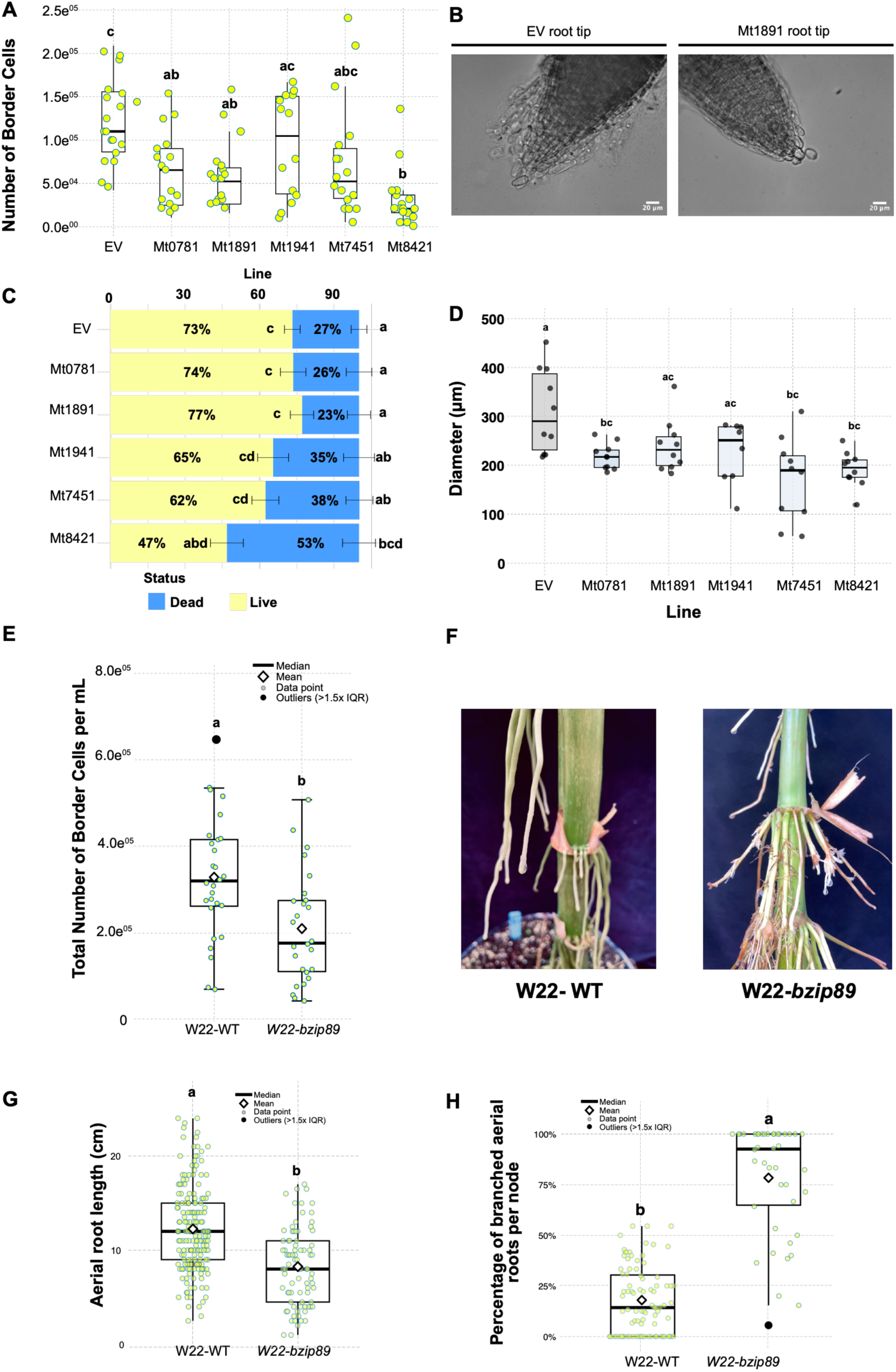
Experimental validation of the gene regulatory network (GRN) using RNAi. Five transcription factors were targeted by RNAi and compared to an empty vector (EV) control. These included a Homeobox-WOX (MtrunA17_Chr5g0400781 - Mt0781), TUBBY (MtrunA17_Chr1g0201891 - Mt1891), MYB (MtrunA17_Chr3g0111941 - Mt1941), Krüppel-like (MtrunA17_Chr1g0207451 - Mt7451), and C2H2 zinc finger (MtrunA17_Chr1g0148421 - Mt8421) transcription factor. A) Quantification of the total number of BCs. Three RNAi lines (MtrunA17_Chr5g0400781 - Mt0781, MtrunA17_Chr1g0201891 - Mt1891, and MtrunA17_Chr1g0148421 – Mt8421) showed a significantly lower number of BCs compared to the EV control. Two biological replicates were performed (n = 8–10 plants per replicate). B) Representative microscopy images of BCs from the EV and MtrunA17_Chr1g0201891 - Mt1891 RNAi lines. C) Percentage of live and dead BCs. The MtrunA17_Chr1g0148421 - Mt8421 line exhibited the most significant reduction in the proportion of live-dead BCs. Two biological replicates (n = 8–10 plants per replicate). D) Root diameter measurements. RNAi lines MtrunA17_Chr5g0400781 – Mt0781, MtrunA17_Chr1g0207451 - Mt7451, and MtrunA17_Chr1g0148421 - Mt8421 had significantly smaller root diameters compared to the EV control. (E) Total number of BCs in the Zmbzip89 mutant line. A significant difference was observed between the wild-type and mutant lines. (F) Representative aerial roots from the ZmbZIP89 both the mutant and wild-type lines. The mutant exhibited increased branching compared with the wild type. (G) Aerial root length in the Zmbzip89 mutant line. A significant difference was observed between the mutant and wild-type lines. (H) Percentage of branched aerial roots in the Zmbzip89mutant line. A significant difference was observed between the mutant and wild-type lines. Statistical analysis was performed using ANOVA, followed by Tukey’s Honest Significant Difference (HSD) test to assess significance between groups (Figure 7A, C - D) and Scott-Knott test (Figure 7E, G - H). All analyses were conducted in R (v4.4.0) using the multcompView package (v0.1-10) and ScottKnott (1.4-0).

Given its pronounced phenotype, evolutionary conservation, and functional relevance, we investigated the regulator MtrunA17_Chr1g0148421 further in the cereals, given that this is a less-characterized gene with a putative C2H2 domain. We constructed a neighbor-joining phylogenetic tree incorporating paralogs from Medicago, maize, and sorghum, which confirmed orthology relationships with strong bootstrap support. Conserved zinc finger domains were identified across three orthologs (**Supplemental Figure. 32**). The phylogenetic analysis and protein sequence alignment confirmed that the maize and sorghum homologs are more closely related, as expected given their shared lineage within the Andropogoneae tribe. However, we identified distinct sets of conserved amino acid residues, some shared exclusively between sorghum and Medicago, and others between maize and Medicago, suggesting possible functional conservation beyond strict phylogenetic relationships. Lastly, we measured root diameter, a trait positively correlated with mucilage production in aerial roots of maize and sorghum (Van Deynze et al. 2018; Pankievicz et al. 2022; Venado et al. 2025). The RNAi lines exhibited reduced root diameters compared to the control but only lines MtrunA17_Chr5g0400781, MtrunA17_Chr1g0207451 and MtrunA17_Chr1g0148421 had a significant reduction in diameter (**Figure 7D**), consistent with reduced BCs. Collectively, these findings support that BC regulators, particularly the novel regulator MtrunA17_Chr1g0148421, are functionally conserved across monocots and dicots, despite their evolutionary divergence.

To further support the GRNs identified in this study, we examined the regulators reported by Gao et al. (2023) to identify findings shared with our regulatory network analysis. Several high-confidence regulators were common to both studies. Notably, *ZmbZIP89*, which Gao et al. reported as a candidate gene, was among the highest degree regulators in topic 5 (enriched for hormone signaling, stress response), in the BC cluster Z9 **(Figure 6B, Supplemental Table 3)**. We investigated its potential role by obtaining a *Zmbzip89* mutant from the UniformMu insertion collection. We examined aerial root traits and mucilage production to determine whether disruption of *ZmbZIP89* affects mucilage synthesis. Quantification of BC production revealed that wild-type plants released significantly more BCs than the *Zmbzip89* mutant, with approximately 328,000 and 209,000 BCs mL⁻¹, respectively **(Figure 7E).** We did not identify significant differences in other aerial root phenotypes, including the number of nodes and the height of the uppermost node **(Supplemental Figure 32C).** However, *Zmbzip89* aerial roots were significantly shorter than those of wild-type plants (**Figure 7F - G**) and frequently exhibited multiple lateral branches (**Figure 7H**). These observations suggest that *ZmbZIP89* gene may have additional roles in aerial root development beyond the regulation of BC production. Collectively, these results provide experimental validation of our GRN predictions and demonstrate that integrating single-nuclei transcriptomics with genetic association analyses is an effective strategy for identifying previously unrecognized regulators of BC development. These findings establish *Zmbzip89* as a candidate regulator of BC production and provide a foundation for elucidating the molecular mechanisms controlling mucilage-producing cells in both monocot and dicot species.

## DISCUSSION

Understanding how plants regulate mucilage production and BC development has long been hampered by the complexity and cellular heterogeneity of root tissues. Here, we addressed this challenge by applying single-nucleus RNA sequencing across three phylogenetically diverse species, maize, sorghum, and Medicago, to capture transcriptional profiles both before and after mucilage secretion and defining gene regulatory networks (GRNs) at the global as well as cell type-specific levels. By integrating snRNAseq with GRN inference and functional validation, we identified conserved transcriptional programs governing BC development and uncovered key regulators whose loss reduces both BC number and viability.

### BCs and mucilage as a multifunctional rhizosphere interface with the environment

BCs and their associated mucilage form a root extracellular trap (RET), a specialized interface containing polysaccharides, defense-related proteins, extracellular DNA, histones, and secondary metabolites (Driouich et al. 2019, 2024). This matrix can contribute to root protection by trapping soilborne pathogens and shaping interactions with the surrounding microbial community (Hawes et al. 2016; Fortier et al. 2023). BC secretions can also contain metabolites that influence beneficial microbes, including compounds involved in rhizobial recruitment and antimicrobial responses (Watson et al. 2015; Kranawetter et al. 2021). In maize and sorghum aerial roots, mucilage additionally provides a microenvironment that supports diazotrophic communities and biological nitrogen fixation (Pankievicz et al. 2022; Venado et al. 2025b). Thus, BC number, viability, and mucilage secretion have the potential to influence several aspects of the root–microbe interface.

Our results provide a regulatory framework for understanding how this interface is established. Across maize, sorghum, and *Medicago*, we identified distinct BC populations and cell-type-specific regulatory networks, with BC-associated modules enriched for processes including cell-wall modification, extracellular functions, ion transport, and stress responses (**Figure 4E**). In maize, the BC-associated network was further connected to regulators involved in cell-wall synthesis and auxin, ethylene, and stress signaling, consistent with the cellular remodeling required for BC formation and mucilage production (**Figure 6B**). Importantly, functional validation demonstrated that perturbation of several predicted regulators altered BC number or viability, while the maize *bzip89* mutant reduced BC release (**Figure 7E**). These results indicate that the capacity of roots to establish this extracellular interface is genetically regulated and can be modulated through specific transcriptional regulators. Although we did not directly measure changes in mucilage chemistry, microbial recruitment, or pathogen resistance, our findings identify candidate regulatory entry points through which BC production could potentially be modified to alter rhizosphere functions such as biological nitrogen fixation and root protection.

### A conserved transcriptional program underlies BC formation from monocots to eudicots

One of the most striking findings of our study is that despite the evolutionary distance between the monocots and Medicago, a core set of transcription factors predicted to regulate BC development is conserved. This conservation is not simply a reflection of shared root cell-type identity; our cross-species pseudo-bulk correlation analyses showed that, outside the quiescent center and meristem, transcriptional programs are generally poorly conserved. The conservation of BC regulators in this broader context of divergence suggests a specific evolutionary constraint, consistent with the fundamental importance of RET across plant lineages (Hawes et al. 2016; Driouich et al. 2024). Our RNA velocity analyses further revealed that the BC lineage exhibits the sharpest and most coherent transcriptional dynamics across all three species, consistent with a tightly regulated developmental program rather than a plastic environmental response.

### Key transcription factors controlling BC number and integrity

Functional validation through RNAi knockdown in Medicago confirmed roles for five transcription factors in BC biology in at least one of the phenotypes measured. Among these, MtrunA17_Chr1g0148421, encoding a C2H2-type zinc finger transcription factor, emerged as the most consequential regulator, affecting not only BC abundance but also cell viability, critical since the RET’s defensive and symbiotic functions depend on metabolically active, living cells (Ropitaux et al. 2020; Driouich et al. 2024). The C2H2 zinc finger family has established roles in root cap development in *Arabidopsis*, where members, including AtZAT14, regulate programmed cell death of outer root cap cells (Fendrych et al. 2014). MtrunA17_Chr5g0400781 - Mt0781 (WOX family) and MtrunA17_Chr1g0201891 - Mt1891 (TUBBY-like) also reduced BC numbers when silenced, pointing to roles upstream of or parallel to MtrunA17_Chr1g0148421 - Mt8421. The absence of significant phenotypes in the MYB and Krüppel-like lines likely reflects redundancy, consistent with *Arabidopsis* findings where quintuple C2H2 mutants display defects absent in single knockouts

The absence of a significant phenotype in the MYB line may reflect functional redundancy within this transcription factor family. The reduction in root diameter in the Krüppel-like line suggests that this factor has a more specific or less redundant role, although functional overlap with related C2H2 zinc-finger proteins cannot be excluded. In Arabidopsis studies have shown that loss of a single C2H2 factor can produce little or no detectable phenotype, whereas higher-order mutants of related ZAT proteins reveal developmental defects, supporting functional redundancy within this family (Feng et al. 2023).

*ZmbZIP89* (*Zm00001eb238570*) is an epidermis- and cortex-enriched transcription factor that regulates root development through the coordination of auxin signaling and reactive oxygen species (ROS) homeostasis (Li et al. 2025; Gul et al. 2026; Zhu et al. 2026). Its role in promoting root cell elongation and architecture makes it a compelling candidate regulator of aerial root cap development. Consistent with this hypothesis, *bzip89* mutants exhibited a significant reduction in BC release but also in other aerial root trats such as length and branching (**Figures E - H**). Given that *ZmbZIP89* directly regulates genes involved in cell elongation and ROS homeostasis, loss of its function may impair the cellular remodeling required for efficient BC separation and release. Together, these findings identify ZmbZIP89 as a previously unrecognized regulator of BC production and demonstrate the predictive power of our gene regulatory network for prioritizing biologically relevant regulators of aerial root mucilage production.

### Implications for engineering nitrogen fixation and root protection

Our identification of regulators controlling BC number and mucilage production has direct implications for two distinct agronomic goals. First, if BC output and RET quality are amenable to genetic manipulation, it may be possible to tune the rhizosphere environment to favor diazotrophic colonization in cereal crops, reducing dependence on synthetic nitrogen fertilizers (Bennett et al. 2020; Pankievicz et al. 2022; Venado et al. 2025b). The metabolite composition of the mucilage, including the flavonoid and organic acid signals documented in Medicago by Kranawetter et al. (2021), is likely to be as important as mucilage quantity in determining whether the rhizosphere microenvironment supports nitrogen-fixing communities, suggesting that regulators of BC identity could influence both the volume and the chemical character of secretions. Second, because the RET is a primary barrier against soilborne pathogens, these same regulators are candidates for improving root disease resistance in legume crops where root rot caused by *Aphanomyces euteiches* remains a major uncontrolled threat (Fortier et al. 2023). The orthology of key BC regulators confirmed here across maize, sorghum, and Medicago suggests that findings in the tractable Medicago system will translate to both cereal and legume crops.

### Methodological considerations

The use of Medicago *Agrobacterium rhizogenes*-based transformation as a validation system, while rapid, introduces the caveat of chimeric root systems; validation in insertion mutants or via CRISPR knockouts in the cereal species will strengthen conclusions. GRN benchmarking confirmed that both MERLIN and scMTNI outperform random expectation, though BC and root cap clusters showed weaker chromatin-based recovery, possibly reflecting the specialized, rapidly changing transcriptional state of these populations during mucilage secretion. Incorporating single-cell ATAC-seq and spatial transcriptomics from the same tissues would substantially improve cell-type-specific GRN resolution in the future.

In summary, this study establishes a multi-species single-cell framework for dissecting the transcriptional control of mucilage production and BC development, identifies conserved regulators with functional roles across monocots and dicots, and connects BC biology to nitrogen fixation, plant immunity, and rhizosphere chemical ecology, where these cells play essential roles.

## MATERIALS AND METHODS

### Plant growth

Sorghum (accession IS23992) and maize (accession CB017456/Oaxa524) were grown in the Walnut Street Greenhouses at the University of Wisconsin-Madison in 2,800-sized pots (7 gallons) filled with Pro-Mix LP15 medium, with daytime temperatures maintained at 28°C and nighttime temperatures at 25°C. The light cycle was set to a 16-hour light photoperiod from 5 am to 9 pm and 8 hours of darkness from 9 pm to 5 am. Plants were irrigated with municipal water twice a week and fertilized weekly with NPK (20-10-20). The greenhouse room was conditioned with a humidifier system (Smart Fog, Reno, Nevada) to 100% relative humidity to induce mucilage production. *Medicago truncatula* seeds were scarified with sulfuric acid and sterilized with 12% sodium hypochlorite (Boisson-Dernier et al. 2001). All seeds were grown in 1.2% agar in the growth chamber at the University of Florida. The chamber was set up with 18 hours of light and 6 hours of darkness. Mucilage production was triggered by applying 10 µL of sterile water directly to the root tip.

### Nuclei isolation and fluorescence-activated cell sorting

Medicago root tips were collected six days post-germination. Nuclei isolation was performed following the protocol described by (Conde et al. 2021). Briefly, tissues were chopped with a blade in Nuclei Isolation Buffer (NIB) for two minutes, and the process was repeated twice. The samples were then filtered through a 40 µm strainer (Greiner Bio-One) placed over a 50 mL Falcon tube, followed by centrifugation at 600 g for 5 minutes at 4°C. The resulting pellet was resuspended in 750 µL of NIB Wash solution for cell sorting. Sorghum and maize aerial root tips, with and without mucilage, were collected at Stage 8 and V12, respectively. For each experiment, three roots from individual plants were pooled per condition. Nuclei isolation from these root tips followed the same procedure as for Medicago, with added double filtration and centrifugation steps to remove larger, more resistant debris from the plant material. Nuclei integrity was assessed using confocal microscopy. DAPI-stained nuclei (5 µg/mL) were visualized with a Leica TCS SP5 confocal microscope for Medicago and a Zeiss LSM780 confocal microscope for maize and sorghum, using laser excitation lines of 405 nm and 488 nm. Subsequently, nuclei were sorted using a BD FACSAria™ IIU/III with filter settings of 50 mW 488 nm (blue), 100 mW 350 nm (UV), 50 mW 635 nm (red), 100 mW 405 nm (violet), and 100 mW 531 nm (yellow/green), along with 15 detectors and FSC/SSC gating. A total of 40,000 nuclei were targeted to achieve a final concentration of 500 nuclei/µL for 10X Chromium technology.

### snRNAseq library preparation and sequencing

We followed the 10X Single Cell V3.1 Dual Index Gene Expression protocol (10X Genomics, Pleasanton, California). In brief, nuclei were loaded at a concentration of 500 nuclei/µL into the 10X Genomics microfluidic chip, along with the required reagents to generate Gel Beads-in-Emulsion (GEMs) using a Chromium X device. cDNA amplification was performed using pre-cleaned GEMs and commercial reagents. Quality control was assessed with an Agilent TapeStation to verify cDNA integrity. The final library was prepared using 10 µL of purified cDNA, followed by index PCR amplification with 13 cycles. Sequencing was conducted using an Illumina NovaSeq 6000 at the UF Interdisciplinary Center for Biotechnology Research (Gainesville, FL) for *Medicago* and at Novogene (Sacramento, California) for maize and sorghum. The sequencing protocol included 28 cycles for Read 1, 10 cycles for both Indexes (I5 and I7), and 90 cycles for Read 2, with a target sequencing depth of 200 million reads per sample.

### Promoter-reporter studies

A 2 kb promoter region from various differentially expressed genes (DEGs) was cloned based on cluster analysis. Amplification of the promoter region from *Medicago truncatula* A17 was performed using Q5 High-Fidelity DNA Polymerase (New England Biolabs) following the manufacturer’s protocol. The amplified promoter was initially cloned into the pJET vector (Thermo Fisher Scientific) using a blunt-end cloning approach and subsequently transferred via Esp3I into a Golden Gate expression vector containing an *NLS-2xYFP* reporter. Whole plasmid sequencing was conducted at Plasmidsaurus. A complete list of plasmids and primers is provided in **(Supplemental Table 4,5).**

### RNA interference

Gateway cloning was used to target BC regulators. RNAi constructs were designed to target a 300–400 bp specific fragment within the first exon of each regulator. Primers were designed using the pssRNAit tool (https://www.zhaolab.org/pssRNAit/). The resulting PCR fragments were cloned into the pENTR™/D-TOPO vector (Thermo-Fisher Scientific) according to the manufacturer’s instructions. The pENTR-Mt (Regulator) plasmid was then recombined into the pK7WIWG2(II): DsRED (Riely et al. 2011) binary vector, which carries a DsRED transformation marker. Whole-plasmid sequencing was performed at Plasmidsaurus. A complete list of plasmids and primers is provided in **Supplemental Tables 4 and 5**.

### *In situ* hybridization

Aerial roots from sorghum (at stage 4) and maize (at stage V12) were collected. Individual roots measuring 25–35 mm were cut into approximately 5 mm segments and fixed in 10% Neutral Buffered Formalin for 24 hours at room temperature, then stored in 70% ethanol for 48 hours before paraffin embedding (SurgPath Leica Paraplast Plus). Two molecular markers were tested in sorghum and maize, with both positive and negative controls included for each species. Probe design was based on the target sequences to ensure high specificity and minimal cross-reactivity **(Supplemental Table 5).** In situ hybridization was carried out using the RNAscope 2.5 HD RED Assay (Advanced Cell Diagnostics - ACD, https://www.bio-techne.com/), and all procedures followed the ACD standard protocol provided by the TRIP Lab at the University of Wisconsin (https://research.pathology.wisc.edu/trip-home/). Briefly, embedded tissues were deparaffinized, treated with hydrogen peroxide for pretreatment, and then hybridized with the target probes. Slides were counterstained with 50% hematoxylin using a HybEZ II oven (PN 321710/321720). For each marker, three independent aerial roots from different plants were analyzed. Slides were scanned with a Leica Aperio AT2 at 40×. Only representative images were included in the results. Image analysis and annotation were performed using Aperio ImageScope (V12.4.6, Leica Biosystems).

### Microscopy

BCs were examined from mucilage samples collected from the aerial roots of maize and sorghum, as well as the underground roots of Medicago, using the previously described genotypes. Approximately 1 µL of mucilage was mixed with 5 µL of distilled water, spread on a microscope slide, and imaged using a Leica DMi8 Model S inverted fluorescence microscope under light microscopy. All images were processed using ImageJ-Fiji (version 2.9.0) (Schindelin et al. 2012) to add scale bars.

Transformed roots were initially screened for the presence of the transformation marker (NES-2xmCherry) using a Leica DFC310FX stereo microscope. Subsequently, confocal imaging was performed with a Zeiss LSM 980 Airyscan 2 microscope. The transformation marker was detected using a DsRed filter (excitation 530–560 nm; emission 565–580 nm), while NLS-2xYFP was visualized with excitation 460–490 nm and emission 520–540 nm. All images were processed using ImageJ-Fiji (version 2.9.0) (Schindelin et al. 2012) to merge channels and add scale bars.

### Plant transformation

Medicago Jemalong A17 was transformed using the hairy root method (Boisson-Dernier et al. 2001). Seedlings were germinated on 1.2% agar, and 36 hours post-germination, root tips were excised and dipped in *Agrobacterium rhizogenes* Arqua1 carrying either reporter or RNAi constructs. The seedlings were then placed in 1.5% agar modified Fahraeus medium supplemented with 0.5 mM NH₄NO₃ and incubated at room temperature for two weeks. Subsequently, plants were transferred to CYG germination pouches (https://mega-international.com/) and grown for an additional two weeks before screening for transformation markers NES-2xmCherry for fluorescent reporters or DsRED for RNAi knockdown constructs using a DFC310 FX stereo microscope (Leica Biosystems). Transformed plants carrying reporters were grown for an additional week before confocal screening, while RNAi-transformed plants were maintained in 1.5% agar Fahraeus medium for one more week prior to phenotyping.

### RT-qPCR

Two tissues were collected from the aerial roots of sorghum and maize at stage 3, as described by Pankievicz et al. (Pankievicz et al. 2022). RNA was extracted from the root cap and a region 1 cm away from the stalk using the Direct-zol RNA MiniPrep Plus with TriReagent (Zymo Research, Irvine, CA). cDNA synthesis was performed using the iScript Advanced cDNA Synthesis Kit for RT-qPCR (Bio-Rad), following the manufacturer’s instructions, with 1 µg of RNA as the starting material. RT-qPCR was conducted with SsoAdvanced SYBR Green Supermix, using a 1:5 (vol/vol) cDNA dilution and 0.25 µM of each primer. Housekeeping genes used for normalization were *SbTubulin* and *SbGAPDH* for sorghum (Sudhakar Reddy et al. 2016) and *ZmUbiquitin* and *ZmActin* for maize(Lin et al. 2014). Reactions were carried out in a CFX Opus 96 Real-Time PCR System (Bio-Rad) under the following conditions: 95°C for 3 minutes, followed by 40 cycles of denaturation at 95°C for 10 s, annealing and extension at 60°C for 30 s. Melting curve analysis 65 to 95°C in 0.5°C increments. Primer sequences used for RT-qPCR are listed in **Supplemental Table 5**.

### BCs and root phenotyping

BCs were visualized using a Leica DMi8 model S inverted fluorescence microscope. Image post-processing and scale bar adjustments were performed with ImageJ - Fiji (version 2.9.0) (Schindelin et al. 2012). Knockdown roots expressing the transformation marker (DsRED) and the RNAi construct were phenotyped by quantifying the number of BC, the percentage of live and dead cells, and root diameter. BC counts were determined using a Countess 3 FL automated cell counter (Invitrogen). Root tips were submerged in 5 µL of distilled water for 60 minutes, then stained with 0.4% Trypan Blue (Invitrogen) for 5 minutes. A 10 µL suspension was loaded onto a Countess cell counting chamber slide (Invitrogen) and analyzed. Parameters were adjusted to exclude debris and non-root-derived cells, optimizing gate size and circularity to detect elongated cells. The total and live-dead cell counts were used to calculate viability percentages for each RNAi line. Root diameter was measured at a consistent position, 100 µm from the root tip, using a Leica DMi8 model S inverted fluorescence microscope. Measurements were taken with the internal ruler of Leica LAS X software (version 2.0.0.14332). Statistical analyses were conducted in R (v4.4.0) using multcompView package (v0.1-10)

### snRNAseq data pre-processing and analysis

Raw data from all species were processed using the CellRanger (V7.0.1) command-line tool developed by 10X Genomics. Briefly, paired-end FASTQ files were demultiplexed and aligned to custom reference genomes generated with the -mkref command. The reference genomes included Medicago Jemalong A17 v5.0 (https://medicago.toulouse.inra.fr/), maize Zm B73 REFERENCE NAM V5.0 (https://www.maizegdb.org/), and *Sorghum bicolor* RTX430 V2.1 (https://phytozome-next.jgi.doe.gov/). Reads from the dry and wet samples mapped to the *Medicago truncatula* A17 r5.0 reference genome at 87% and 90%. For maize, B73 V5 reference genome reads mapped at 46% and 63% for dry and wet samples, respectively. For sorghum, reads mapped at 87% and 90% for dry and wet samples, respectively **(Supplemental Table 1)**. For each species and condition (dry and wet), we excluded genes expressed in fewer than five cells per sample and removed cells with fewer than 1,000 reads. Expression values were depth-normalized to 1,000 reads per cell and variance-stabilized using an inhouse R implementation of the pagoda2package (Barkas et al. 2021). Dried and watered samples were then merged into a single expression matrix for each species by taking the union of genes across samples and assigning zeros to missing expression values. The resulting expression matrices comprised 25,126 genes × 4,183 cells for Medicago, 26,131 genes × 7,553 cells for maize, and 21,224 genes × 11,526 cells for sorghum. Processed output data were analyzed using various pipelines for cluster cell annotation, RNA velocity, and gene regulatory network construction with MERLIN (Roy et al. 2013) and scMTNI (Zhang et al. 2023)

### Clustering, cell type annotation and lineage tree inference

Nonnegative matrix factorization (NMF) was first applied using the ordinary least squares implementation (NMF-OLS) in MATLAB R2021b, separately for each species, to identify primary transcriptional clusters. Specifically, NMF decomposes the input expression matrix *X* ∈ *R^m^*^×*n*^ into lower-dimensional factors *U* ∈ *R^m^*^×*k*^ and *V* ∈ *R^k^*^×*n*^ by minimizing 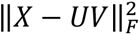, where *m* and *n* denote the numbers of cells and genes, respectively. NMF was performed with *k* values of 10, 15, and 20. To account for variation due to random initialization, each NMF run was repeated ten times with different random seeds, and clustering stability was evaluated using the Jaccard index (JI). For each *k*, the initialization yielding the highest cumulative similarity among clusters was retained. For maize and sorghum, NMF-OLS produced unbalanced cluster distributions, characterized by one or two dominant clusters and several small or empty ones (**Supplemental Figure. 33**). To obtain a more balanced and biologically meaningful clustering, NMF was applied based on alternating non-negativity constrained least squares (NMF-ANLS) using the block principal pivoting algorithm (Kim and Park, 2008), incorporating sparsity constraints in the gene space. The objective function included regularization parameters α and β controlling matrix growth and sparsity, respectively, both set to the mean of the input matrix values. The objective of constrained NMF-ANLS is defined as:

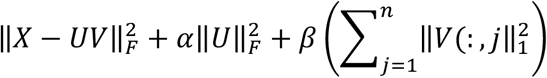

NMF-ANLS was performed with the same *k* values and stability selection protocol as NMF-OLS. The resulting clusters showed high stability (JI range: 0.42 − 0.95 for maize and 0.55 − 0.99 for sorghum). Cluster annotation was performed by identifying differentially expressed genes (DEGs) for each cluster using three statistical approaches. Genes were expressed in more than 50% of cells within a cluster and exhibited a ratio *E*(*x*|*Ini*)⁄*E*(*x*|*Outi*) greater than 1.25, where *E*(*x*|*Ini*) and *E*(*x*|*Outi*) were considered cluster specific. *E*(*x*|*Ini*) and *E*(*x*|*Outi*) denote the fractions of cells expressing gene *x* inside and outside cluster *i*, respectively. Statistical significance of overrepresentation was evaluated using a hypergeometric test with Benjamini–Hochberg correction. Welch’s *t*-test and Mann–Whitney (Wilcoxon rank-sum) tests were also applied to detect differential expression between cluster and non-cluster cells, using the *diffxpy* v0.7.4 package integrated in Scanpy v1.9.3 (Wolf et al. 2018). Only DEGs significant across all three tests (*P* < 0.05) were retained. To assign biological identities, each cluster’s DEG set was compared to known cell type-specific marker genes **(Supplemental data set 1)** using hypergeometric enrichment with *FDR* < 0.05. Additional enrichment was performed using the NMF maximal dimension (maxdim) gene sets, in which genes were assigned to their highest factor loading for each latent dimension. A consensus annotation for each cluster at each *k* value was then defined as the cell-type assignment supported by at least two of the four independent methods **(Supplemental data set 2).** Relationships among clusters were assessed using pairwise Euclidean distances between average expression profiles. A minimum spanning tree (MST) was constructed using the SciPy function *sparse. csgraph. minimum_spanning_tree*, identifying the minimal set of edges connecting clusters based on expression similarity. The MST was visualized as a hierarchical network, where each node was represented as a pie chart showing the proportions of cells from wet and dry conditions. Consensus annotations were integrated with the MST topology and prior biological knowledge to select the optimal *k* value for each species and assign final cell-type labels (**Figure 1D**, **Supplemental data set 2).** Finally, biologically informed lineage-like trees were constructed, leveraging the MST structure and literature on root cell-type organization and function, to illustrate relationships among cell types across environmental conditions (**Figure 3A**).

### Inference of gene regulatory networks with MERLIN

Gene regulatory networks (GRNs) were inferred separately for each species using the MERLIN algorithm (Roy et al. 2013), which reconstructs dependency networks through regularized regression within a probabilistic graphical model framework. MERLIN identifies both individual regulatory interactions and co-regulated gene modules. Candidate regulator lists were compiled from curated databases and genome annotations. For *Medicago truncatula* (Medicago), regulators were obtained from the IMGA 3.5v4 genome annotation (Larrainzar et al. 2015); for maize, transcription factors were included from (Burdo et al. 2014) Burdo et al. (2014) and PlantTFDB (https://planttfdb.gao-lab.org/); and for sorghum, from PlantTFDB. Gene identifiers were standardized across assemblies using MaizeMine, the Grassius Translation Tool, the JGI Genome Portal, and Ensembl Gramene. To expand these sets, orthologous regulators were identified across six plant species (*Sorghum bicolor*, *Zea mays*, *Brassica rapa*, *Arabidopsis thaliana*, *Medicago truncatula*, *Solanum lycopersicum*) using OrthoFinder-v2.5.5 (Emms and Kelly 2019), yielding final candidate regulator lists of 2,435 for Medicago, 2,280 for maize, and 1,924 for sorghum. Each expression matrix was divided into cluster-specific subsets, filtering out genes expressed in fewer than 20 cells per cluster and excluding poorly represented clusters. Filtered matrices were log-transformed before running MERLIN. For stability selection, 100 random subsamples (50% of cells each) were generated, and MERLIN was run independently on each subsample to produce 100 GRNs. These were integrated into a consensus network by retaining regulatory edges occurring in at least 80% of runs (confidence ≥0.8) Consensus gene modules were derived by constructing a gene–gene co-clustering matrix based on module co-occurrence across subsamples, followed by hierarchical clustering and optimal leaf ordering. The co-clustering threshold was selected based on maximal enrichment for GO terms, resulting in cut-offs of 0.3 for Medicago and sorghum, and 0.2 for maize. GO enrichment was performed using hypergeometric tests with *FDR* < 0.05. Regulator enrichment within modules was evaluated using the same statistical framework to identify key transcriptional regulators associated with each module.

### Cell cluster specific network inferences using scMTNI

Single-cell Multi-Task Network Inference (scMTNI) (Zhang et al. 2023) was applied to infer gene regulatory networks (GRNs) from single-cell RNA-seq data across Medicago, maize, and sorghum. scMTNI infers cluster-specific GRNs by modeling them as probabilistic graphical models and using multitask learning guided by cell lineage hierarchies as probabilistic priors. Filtered and normalized expression matrices were generated for each species by removing genes expressed in fewer than 20 cells per cluster and excluding clusters with low gene representation. We combined cluster-specific gene lists into a union gene set and appended identifiers to preserve cluster identity. The same transcriptional regulator lists used in the MERLIN analysis were applied. Cell lineage relationships were inferred from a minimum spanning tree constructed from cluster-averaged expression profiles, and lineage-like trees were used to define cluster order and biological context. For each cluster, we generated 100 subsampled, log-normalized expression datasets and ran scMTNI independently on each subsample. Consensus GRNs were obtained by aggregating edges across subsamples, assigning confidence scores based on edge frequency. For downstream analyses, the inferred networks were filtered using two complementary criteria, retaining either only the top edges or those above defined confidence thresholds. Summary statistics for these consensus networks, including the number of regulators, targets, and edges across thresholds, are provided for the three species and per cell type **(Supplemental data set 5).** The resulting cell type–specific GRNs were analyzed using k-means edge clustering and Latent Dirichlet Allocation (LDA) topic modeling to identify dynamic regulatory modules. GO enrichment for each topic was assessed using a hypergeometric test with false discovery rate (FDR) correction (*FDR* < 0.05).

### Evaluation of inferred GRNs using gold standards

Gold standard regulatory datasets were generated using bulk ATAC-seq from maize and *Medicago*, and ChIP-seq from sorghum. For maize, we used ATAC-seq data (GSE152046; Crisp et al. 2020); for Medicago, two ATAC-seq datasets (GSE101482 and GSE154845; Maher et al. 2018; Knaack et al. 2022); and for sorghum, a ChIP-seq–derived H3K27ac dataset (PRJNA952350; Chen et al. 2024). Maize root tip ATAC-seq data were aligned to the B73 reference genome (RefGen v5.0), while Medicago root tip datasets were mapped to the A17 genome (v5.0). Sorghum root H3K27ac ChIP-seq controls were aligned to the BTx623 genome. To ensure consistency between the gold-standard networks and the snRNA-seq datasets, we mapped the prior networks gene IDs to the genome versions used for the snRNA-seq analysis. For maize, RefGen v3 gene IDs were mapped to B73 RefGen v5 using a cross-reference table, Grassius (https://grassius.org/translation_tool), and Gramene (https://ensembl.gramene.org/Zea_mays/Info/Index). For sorghum, BTx623 gene IDs were mapped to RTx430 v2.1 using a mapping table from Phytozome (https://phytozome-next.jgi.doe.gov/). For Medicago truncatula, Mt4 gene IDs were mapped to A17 Mt5.0 using an Mt4-to-Mt5 mapping table (https://medicago.toulouse.inra.fr/MtrunA17r5.0-ANR/). Only transcription factors that could be mapped to the genome annotations used for the snRNA-seq analysis were retained in the corresponding gold-standard networks. Reads were downloaded using the SRA Toolkit-3.0.2, mapped with Bowtie2-2.2.1, and processed with Samtools-1.9 to retain reads with mapping quality ≥3, remove PCR duplicates, and generate sorted, indexed BAM files. Peaks were called with MACS2-v2.2.6 in paired-end mode (-f BAMPE) using effective genome sizes of 2.12×10⁹ bp (maize), 6.6×10⁸ bp (sorghum), and 4.2×10⁸ bp (Medicago). We linked regulatory elements to target genes by intersecting peak regions with transcription factor (TF) motifs and promoters. Motif information from the Cis-BP database was scanned across both genome strands using the PIQ toolkit (v1.3; pwmmatch.exact.r). We defined promoters as ±5 kb around transcription starts sites (TSS), with additional ±1–5 kb windows used for comparison. We merged overlapping motif–peak and peak–TSS regions to generate TF–gene connections. Prior networks from biological replicates were merged by averaging confidence scores of overlapping edges, percentile-normalized using rankedges, and the top 20% of edges were retained as the gold standard networks **(Supplemental Table 2).** The accuracy of MERLIN- and scMTNI-inferred networks was evaluated against these gold standards using the area under the precision–recall curve (AUPR), computed with the *AUCCalculator* package (Davis and Goadrich 2006). Precision was defined as the proportion of predicted edges found in the gold standard and recall as the fraction of gold standard edges recovered. We also calculated early precision, recall, and F-scores using fixed confidence thresholds (0.9–0.6) and edge cutoffs (top 1,000–30,000) to assess high-confidence predictions. Randomized networks were generated by shuffling edge scores within each method’s consensus network to establish baseline enrichment. Biological relevance was further assessed by identifying predictable TFs, regulators whose predicted targets significantly overlapped with gold standard targets using a hypergeometric test with FDR correction (Q < 0.05) (Marbach et al., 2016). Predictable TFs were evaluated in the top 30,000 and 50,000 edges for both MERLIN and aggregated scMTNI networks, as well as within key scMTNI clusters: maize (QC1 [Z2], Cortex/Endodermis [Z6], QC2 [Z7], BCs [Z9]); *Medicago* (Cortex [M1], Root Cap/Columella [M3], Root Cap/BCs [M4], Endodermis [M8]); and sorghum (Cortex/Endodermis [S4], QC [S7+S9+S13+S10], Root Cap [S11], BCs [S12]).

### Checking the ortholog expression in pseudo-bulk form

Orthogroups ID (OGID) were analyzed to investigate cross-species expression. We built a matrix to identify orthogroups with low duplication levels: those containing only the “_0” suffix, or both “_0” and “_1” (duplication level ≤1). We retained orthogroups containing genes from all three species or missing genes from only one species, resulting in four orthogroup (OG) sets: (1) no duplications, no missing values (2,938 shared genes); (2) no duplications, one missing value (6,009 genes); (3) up to one duplication, no missing values (3,990 genes); and (4) up to one duplication, one missing value (8,533 genes). For each gene in each OG set, we computed average log-transformed expression across all cells within each cluster to generate pseudo-bulk expression profiles. We then grouped genes into 20 clusters based on their pseudo-bulk expression patterns using k-means clustering, and generated heatmaps with clusters grouped either by species or by shared annotations.

To assess expression conservation, we computed Pearson correlation coefficients between pseudo-bulk profiles of all clusters in an all-by-all comparison, producing symmetric correlation matrices. Each matrix was visualized in two orderings: (i) rows and columns grouped by species and (ii) by shared cluster annotations.

### RNA velocity analysis

Unspliced (pre-mature) and spliced (mature) transcripts were quantified using velocyto-v0.17.17 (https://velocyto.org/velocyto.py/index.html), executed directly on each Cell Ranger output directory for both watered and dried root samples, and the resulting count matrices were merged across replicates. RNA velocity and pseudotime analyses were performed using the scVelo-v0.3.1 dynamical model framework (Bergen et al. 2020). Following standard preprocessing, we selected the 2,000 most variable genes by dispersion, fitted the dynamical model to estimate transcriptional kinetics, and reconstructed latent temporal trajectories. Automatic root detection failed in maize (“identified 0 region of root cells and 1 region of endpoints”) and produced multiple ambiguous roots in sorghum. Therefore, root cells were manually assigned as quiescent-center (QC) clusters (Z2 in maize and S7+S9+S10+S13 in sorghum) before recomputing latent time.

Genes were retained as velocity genes if they met quality thresholds (coefficient of determination ≥ 0.01, fit likelihood ≥ 0.001) and had sufficient spliced/unspliced counts, resulting in 1,068 velocity genes in Medicago, 1,047 in maize, and 912 in sorghum. We identified candidate driver genes based on higher fit likelihood and partial gene likelihoods computed within each cluster. Finally, we intersected the 2,000 highly variable genes with curated marker sets and inferred GRNs to link transcriptional dynamics with known cell-type markers and GRN-predicted regulators.

### Plant material and growing conditions for maize mutants

Seeds from the following maize genotypes were requested from germplasm banks (accession numbers provided): (i) a zmbzip89::mu1038804 mutant (stock UFMu-03763, mu1038804, Maize Genetics Cooperation Stock Center) in the W22 inbred background, carrying a Mu transposon insertion within an exon of Zm00001eb238570 (zmbzip89), with a 9-bp target-site duplication (Springer et al. 2018); (ii) its near-isogenic parental line W22 (PI 693394 and NSL 30053, United States Department of Agriculture National Plant Germplasm System, USDA-NPGS), hereafter referred to as W22-wt, which carries the functional wild-type ZmbZIP89 allele.

The experiment was carried out in the Walnut Street Greenhouses (University of Wisconsin-Madison, Madison, WI, USA). Thirty pots (15 replicates per accession), each measuring 35 cm in diameter by 28 cm in height, were filled with a previously prepared substrate consisting of one 107-liter bag of BM6 All Purpose Mix (Berger Growing Mixes), a standard mix of coarse-grade peat moss, perlite, vermiculite, non-ionic wetting agent, and dolomitic and calcitic limestone, combined with one 50-lb (22.7 kg) bag of Turface Athletics MVP (PROFILE Products), a calcined clay amendment added to improve soil drainage and aeration. The substrate was homogenized and fertilized with slow-release granular Nutricote Total (NPK 13-11-11 plus micronutrients) at 10 g per pot. Three seeds per pot were sown; thinning was performed at the V2-V3 developmental stage to maintain one plant per pot. Soil moisture was maintained by watering 2-3 times per week, keeping the substrate moist but never waterlogged. Greenhouse temperatures were maintained at 27 ± 1°C (daytime) and 19 ± 1°C (nighttime), with a relative humidity of 80-85%. Plants were grown under a 16 h/8 h light/dark photoperiod with supplemental lighting providing a minimum photosynthetic photon flux density of 250 µmol m⁻² s⁻¹.

### BC counting and aerial root phenotyping in maize W22 wild-type line and *bzip89* mutant

Eight weeks after planting, 3 cm-long healthy aerial roots were excised from plants and placed into 2 mL microtubes containing 1 mL of 1× Phosphate-Buffered Saline (PBS, pH ≈ 7.4) to stimulate mucilage release (Pankievicz et al. 2022; Fathinejad et al. 2026) without causing osmotic stress in the BCs. After 48 hours, each solution (PBS + mucilage) was gently homogenized; 50 µL was then collected, mixed in 0.2 mL tubes with 20 µL of 0.4% Trypan blue dye, incubated for 1-2 minutes at room temperature, and pipetted into both sides of the cell counting chamber slides (C10283, Invitrogen). Each side was read three times using the countess 3 FL Automated Cell Counter (AMQAF2001, Invitrogen); the dye dilution ratio was adjusted in the cell counter settings, and the mean of six readings per aerial root at each time point was used as input for the statistical analyses. In addition, the plants were phenotyped for the following traits: number of nodes forming aerial roots, height of the top node forming aerial roots, number of aerial roots per node, length and diameter of aerial roots (means of six roots per plant), and stalk diameter (Laspisa et al. 2025). In all analyses, the Scott-Knott clustering algorithm at the 5% significance level was applied for mean-comparison tests (Jelihovschi et al. 2014).

### Phylogenetic analyses

Protein sequences of the C2H2 zinc finger orthologs of Medicago, maize and sorghum, together with sequences of *Arabidopsis thaliana*, *Solanum lycopersicum* and *Physcomitrium patens*, were aligned with CLC Genomics Workbench (Ver 8.0) using default setting gap open cost 15.0, and gap extension cost 1.0. The phylogeny was inferred with CLC Genomics Workbench (Ver 8.0) under the Jukes-Cantor model. Node support was estimated from 100 bootstrap replicates. The tree was rooted using *Physcomitrium patens* as an outgroup, based on its more distant phylogenetic, and branch lengths represented as 1 unit.

## AUTHOR CONTRIBUTIONS

R.E.V., M. Kotvanova., and C.K. contributed equally to this work. M.Kotvanova, C.K, P.G. and S.P. contributed to the computational analyses. R.D. and T.K. contributed to the greenhouse experiments. C.D., W.J.P., and M. Kirst. contributed with experimental tools. S.R. and J.M.A. designed the research, analyzed the data, and wrote the manuscript. All authors read and approved the final version of the manuscript.

## ACKNOWLEDGMENTS

The authors acknowledge the Maize Genetics Cooperation Stock Center and the USDA National Plant Germplasm System for providing material. During the preparation of this manuscript, Claude (Claude Opus 4.8, Anthropic, PBC; available at Claude AI) was used to improve the clarity and readability of the text. After using this tool, the authors reviewed and edited the content as needed, and they take full responsibility for the content of this publication.

## DATA AVAILABILITY

The single-nucleus RNA-sequencing data generated in this study have been deposited in the NCBI Gene Expression Omnibus (GEO) under accession number GSE348174 and in the NCBI Sequence Read Archive (SRA) under BioProject accession number PRJNA1532956. The data will be publicly available upon publication. The plasmids generated in this study are available upon request. The code used for clustering, RNA velocity, and network inference is available at https://github.com/Roy-lab/SoNar-data-analysis. Previously published datasets used to build the gold standard networks are available under GSE101482 and GSE154845 (Medicago ATAC-seq), GSE152046 (maize ATAC-seq), and PRJNA952350 (sorghum H3K27ac ChIP-seq).

## SUPPLEMENTAL DATA SETS

**Supplemental data set 1:** Cell type specific DEG list from each species

**Supplemental data set 2:** Cell-cluster annotations.

**Supplemental data set 3:** scVelo dynamical latent time

**Supplemental data set 4:** MERLIN module statistics

**Supplemental data set 5:** Topological properties of inferred networks

**Supplemental data set 6:** Top Regulators from kmeans and LDA topics

## REFERENCES

AUCCalculator 0.2. https://mark.goadrich.com/programs/AUC/. Retrieved September 20, 2026

Awika JM. Major Cereal Grains Production and Use around the World. . In. Advances in Cereal Science: Implications to Food Processing and Health Promotion, ACS Symposium Series. (American Chemical Society), pp. 1–13. 10.1021/bk-2011-1089.ch001

Barkas N, Petukhov V, Kharchenko P, and Biederstedt E. Pagoda2: Single cell analysis and differential expression. R package version. 2021.

Bennett AB, Pankievicz VCS, and Ané J-M. A model for nitrogen fixation in cereal crops. Trends Plant Sci. 2020:25(3):226–235. 10.1016/j.tplants.2019.12.004

Bergen V, Lange M, Peidli S, Wolf FA, and Theis FJ. Generalizing RNA velocity to transient cell states through dynamical modeling. Nat Biotechnol. 2020:38(12):1408–1414. 10.1038/s41587-020-0591-3

Boisson-Dernier A, Chabaud M, Garcia F, Bécard G, Rosenberg C, and Barker DG. Agrobacterium rhizogenes-transformed roots of Medicago truncatula for the study of nitrogen-fixing and endomycorrhizal symbiotic associations. Mol Plant Microbe Interact. 2001:14(6):695–700. 10.1094/mpmi.2001.14.6.695

Brinkmann V, Reichard U, Goosmann C, Fauler B, Uhlemann Y, Weiss DS, Weinrauch Y, and Zychlinsky A. Neutrophil extracellular traps kill bacteria. Science. 2004:303(5663):1532–1535. 10.1126/science.1092385

Burdo B, Gray J, Goetting-Minesky MP, Wittler B, Hunt M, Li T, Velliquette D, Thomas J, Gentzel I, dos Santos Brito M, et al. The Maize TFome--development of a transcription factor open reading frame collection for functional genomics. Plant J. 2014:80(2):356–366. 10.1111/tpj.12623

Cannesan MA, Gangneux C, Lanoue A, Giron D, Laval K, Hawes M, Driouich A, and Vicré-Gibouin M. Association between border cell responses and localized root infection by pathogenic Aphanomyces euteiches. Ann Bot. 2011:108(3):459–469. 10.1093/aob/mcr177

Cervantes-Pérez SA, Thibivilliers S, Laffont C, Farmer AD, Frugier F, and Libault M. Cell-specific pathways recruited for symbiotic nodulation in the Medicago truncatula legume. Mol Plant. 2022:15(12):1868–1888. 10.1016/j.molp.2022.10.021

Chen M, Ji M, Wen B, Liu L, Li S, Chen X, Gao D, and Li L. GOLDEN 2-LIKE transcription factors of plants. Front Plant Sci. 2016:7:1509. 10.3389/fpls.2016.01509

Chen X, He C, Xu H, Zeng G, Huang Q, Deng Z, Qin X, Shen X, and Hu Y. Characterization of the SWI/SNF complex and nucleosome organization in sorghum. Front Plant Sci. 2024:15:1430467. 10.3389/fpls.2024.1430467

Conde D, Triozzi PM, Balmant KM, Doty AL, Miranda M, Boullosa A, Schmidt HW, Pereira WJ, Dervinis C, and Kirst M. A robust method of nuclei isolation for single-cell RNA sequencing of solid tissues from the plant genus Populus. PLoS One. 2021:16(5):e0251149. 10.1371/journal.pone.0251149

Crisp PA, Marand AP, Noshay JM, Zhou P, Lu Z, Schmitz RJ, and Springer NM. Stable unmethylated DNA demarcates expressed genes and their cis-regulatory space in plant genomes. Proc Natl Acad Sci U S A. 2020:117(38):23991–24000. 10.1073/pnas.2010250117

Davis J and Goadrich M. The relationship between Precision-Recall and ROC curves. . In. Proceedings of the 23rd international conference on Machine learning - ICML ’06. (ACM Press: New York, New York, USA). 10.1145/1143844.1143874

Denyer T, Ma X, Klesen S, Scacchi E, Nieselt K, and Timmermans MCP. Spatiotemporal developmental trajectories in the Arabidopsis root revealed using high-throughput single-cell RNA sequencing. Dev Cell. 2019:48(6):840–852.e5. 10.1016/j.devcel.2019.02.022

D’haeseleer K, Den Herder G, Laffont C, Plet J, Mortier V, Lelandais-Brière C, De Bodt S, De Keyser A, Crespi M, Holsters M, et al. Transcriptional and post-transcriptional regulation of a NAC1 transcription factor in Medicago truncatula roots. New Phytol. 2011:191(3):647–661. 10.1111/j.1469-8137.2011.03719.x

Driouich A, Durand C, Cannesan M-A, Percoco G, and Vicré-Gibouin M. Border cells versus border-like cells: are they alike? J Exp Bot. 2010:61(14):3827–3831. 10.1093/jxb/erq216

Driouich A, Gueye M-LF, Vicré M, and Moore JP. The root extracellular trap; a complex and dynamic biomatrix network essential for plant protection. Curr Opin Plant Biol. 2024:82(102656):102656. 10.1016/j.pbi.2024.102656

Driouich A, Smith C, Ropitaux M, Chambard M, Boulogne I, Bernard S, Follet-Gueye M-L, Vicré M, and Moore J. Root extracellular traps versus neutrophil extracellular traps in host defence, a case of functional convergence? Biol Rev Camb Philos Soc. 2019:94(5):1685–1700. 10.1111/brv.12522

Emms DM and Kelly S. OrthoFinder: phylogenetic orthology inference for comparative genomics. Genome Biol. 2019:20(1):238. 10.1186/s13059-019-1832-y

FAO Cereal Supply and Demand Brief. WorldFoodSituation. https://www.fao.org/worldfoodsituation/csdb/en. Retrieved December 1, 2024

Fathinejad A, Sobhtalab Z, Mohammadi MH, and Nazari M. Root type governs mucilage secretion and border cell release in maize. Rhizosphere. 2026:38(101387):101387. 10.1016/j.rhisph.2026.101387

Fendrych M, Van Hautegem T, Van Durme M, Olvera-Carrillo Y, Huysmans M, Karimi M, Lippens S, Guérin CJ, Krebs M, Schumacher K, et al. Programmed cell death controlled by ANAC033/SOMBRERO determines root cap organ size in Arabidopsis. Curr Biol. 2014:24(9):931–940. 10.1016/j.cub.2014.03.025

Feng Q, Cubría-Radío M, Vavrdová T, De Winter F, Schilling N, Huysmans M, Nanda AK, Melnyk CW, and Nowack MK. Repressive ZINC FINGER OF ARABIDOPSIS THALIANA proteins promote programmed cell death in the Arabidopsis columella root cap. Plant Physiol. 2023:192(2):1151–1167. 10.1093/plphys/kiad130

Fortier M, Lemaitre V, Gaudry A, Pawlak B, Driouich A, Follet-Gueye M-L, and Vicré M. A fine-tuned defense at the pea root caps: Involvement of border cells and arabinogalactan proteins against soilborne diseases. Front Plant Sci. 2023:14:1132132. 10.3389/fpls.2023.1132132

Frank M, Fechete LI, Tedeschi F, Nadzieja M, Nørgaard MMM, Montiel J, Andersen KR, Schierup MH, Reid D, and Andersen SU. Single-cell analysis identifies genes facilitating rhizobium infection in Lotus japonicus. Nat Commun. 2023:14(1):7171. 10.1038/s41467-023-42911-1

Gao J, Feng P, Zhang J, Dong C, Wang Z, Chen M, Yu Z, Zhao B, Hou X, Wang H, et al. Enhancing maize’s nitrogen-fixing potential through ZmSBT3, a gene suppressing mucilage secretion. J Integr Plant Biol. 2023:65(12):2645–2659. 10.1111/jipb.13581

Gautam V, Singh A, Yadav S, Singh S, Kumar P, Sarkar Das S, and Sarkar AK. Conserved LBL1-ta-siRNA and miR165/166-RLD1/2 modules regulate root development in maize. Development. 2021:148(1):dev190033. 10.1242/dev.190033

Gul H, Gul S, and Shahzad A. Rooting for survival: bZIP89 tunes cell-specific root plasticity and drought adaptation in maize. Plant Sci. 2026:362(112774):112774. 10.1016/j.plantsci.2025.112774

Hawes M, Allen C, Turgeon BG, Curlango-Rivera G, Minh Tran T, Huskey DA, and Xiong Z. Root border cells and their role in plant defense. Annu Rev Phytopathol. 2016:54:143–161. 10.1146/annurev-phyto-080615-100140

Hawes MC, Bengough G, Cassab G, and Ponce G. Root caps and rhizosphere. J Plant Growth Regul. 2002:21(4):352–367. 10.1007/s00344-002-0035-y

Hawes MC, Curlango-Rivera G, Xiong Z, and Kessler JO. Roles of root border cells in plant defense and regulation of rhizosphere microbial populations by extracellular DNA “trapping.” Plant Soil. 2012:355(1):1–16. 10.1007/s11104-012-1218-3

Hawes MC, Gunawardena U, Miyasaka S, and Zhao X. The role of root border cells in plant defense. Trends Plant Sci. 2000:5(3):128–133. 10.1016/s1360-1385(00)01556-9

Hawes MC and Pueppke SG. Sloughed peripheral root cap cells: Yield from different species and callus formation from single cells. Am J Bot. 1986:73(10):1466–1473. 10.1002/j.1537-2197.1986.tb10892.x

Heim MA, Jakoby M, Werber M, Martin C, Weisshaar B, and Bailey PC. The basic helix-loop-helix transcription factor family in plants: a genome-wide study of protein structure and functional diversity. Mol Biol Evol. 2003:20(5):735–747. 10.1093/molbev/msg088

Jaiswal V, Kakkar M, Kumari P, Zinta G, Gahlaut V, and Kumar S. Multifaceted roles of GRAS transcription factors in growth and stress responses in plants. iScience. 2022:25(9):105026. 10.1016/j.isci.2022.105026

Javelle M, Vernoud V, Depège-Fargeix N, Arnould C, Oursel D, Domergue F, Sarda X, and Rogowsky PM. Overexpression of the epidermis-specific homeodomain-leucine zipper IV transcription factor Outer Cell Layer1 in maize identifies target genes involved in lipid metabolism and cuticle biosynthesis. Plant Physiol. 2010:154(1):273–286. 10.1104/pp.109.150540

Jelihovschi E, Faria JC, and Allaman IB. ScottKnott: A package for performing the Scott-Knott clustering algorithm in R. TEMA Tend Mat Apl Comput. 2014:15(1):003. 10.5540/tema.2014.015.01.0003

Kim D, Tran A, Kim HJ, Lin Y, Yang JYH, and Yang P. Gene regulatory network reconstruction: harnessing the power of single-cell multi-omic data. NPJ Syst Biol Appl. 2023:9(1):51. 10.1038/s41540-023-00312-6

Kim H and Park H. Nonnegative matrix factorization based on alternating nonnegativity constrained least squares and active set method. SIAM J Matrix Anal Appl. 2008:30(2):713–730. 10.1137/07069239x

Knaack SA, Conde D, Chakraborty S, Balmant KM, Irving TB, Maia LGS, Triozzi PM, Dervinis C, Pereira WJ, Maeda J, et al. Temporal change in chromatin accessibility predicts regulators of nodulation in Medicago truncatula. BMC Biol. 2022:20(1):252. 10.1186/s12915-022-01450-9

Kranawetter C and Sumner LW. Differential root zone secretions and the role of root border cells in rhizosphere manipulation. Phytochem Rev. 2025:24(6):5639–5658. 10.1007/s11101-025-10084-y

Kranawetter C, Zeng S, Joshi T, and Sumner LW. A Medicago truncatula Metabolite Atlas enables the visualization of differential accumulation of metabolites in root tissues. Metabolites. 2021:11(4):238. 10.3390/metabo11040238

Larrainzar E, Riely BK, Kim SC, Carrasquilla-Garcia N, Yu H-J, Hwang H-J, Oh M, Kim GB, Surendrarao AK, Chasman D, et al. Deep sequencing of the Medicago truncatula root transcriptome reveals a massive and early interaction between Nodulation factor and ethylene signals. Plant Physiol. 2015:169(1):233–265. 10.1104/pp.15.00350

Laspisa D, Venado RE, Diogo R, Ane J-M, and Wallace JG. Genetic determinants of aerial root traits that support biological nitrogen fixation in maize. bioRxiv. 2025. 10.1101/2025.05.30.657053

Lin Y, Zhang C, Lan H, Gao S, Liu H, Liu J, Cao M, Pan G, Rong T, and Zhang S. Validation of potential reference genes for qPCR in maize across abiotic stresses, hormone treatments, and tissue types. PLoS One. 2014:9(5):e95445. 10.1371/journal.pone.0095445

Li P, Zhu T, Wang Y, Zhang X, Yang X, Fang S, Li W, Rui W, Yang A, Duan Y, et al. Natural variation in a cortex/epidermis-specific transcription factor bZIP89 determines lateral root development and drought resilience in maize. Sci Adv. 2025:11(15):eadt1113. 10.1126/sciadv.adt1113

Lopez L, Perrella G, Calderini O, Porceddu A, and Panara F. Genome-wide identification of histone modification gene families in the model legume Medicago truncatula and their expression analysis in nodules. Plants. 2022:11(3):322. 10.3390/plants11030322

Lu Z, Liu H, Kong Y, Wen L, Zhao Y, Zhou C, and Han L. Late elongated hypocotyl positively regulates salt stress tolerance in Medicago truncatula. Int J Mol Sci. 2023:24(12):9948. 10.3390/ijms24129948

Iyer NR, Shin J, Cuskey S, Tian Y, Nicol NR, Doersch TE, Seipel F, McCalla SG, Roy S, and Ashton RS. Modular derivation of diverse, regionally discrete human posterior CNS neurons enables discovery of transcriptomic patterns. Sci Adv. 2022:8(39):eabn7430.

Maher KA, Bajic M, Kajala K, Reynoso M, Pauluzzi G, West DA, Zumstein K, Woodhouse M, Bubb K, Dorrity MW, et al. Profiling of accessible chromatin regions across multiple plant species and cell types reveals common gene regulatory principles and new control modules. Plant Cell. 2018:30(1):15–36. 10.1105/tpc.17.00581

Ma H, Liu C, Li Z, Ran Q, Xie G, Wang B, Fang S, Chu J, and Zhang J. ZmbZIP4 contributes to stress resistance in maize by regulating ABA synthesis and root development. Plant Physiol. 2018:178(2):753–770. 10.1104/pp.18.00436

Marbach D, Lamparter D, Quon G, Kellis M, Kutalik Z, and Bergmann S. Tissue-specific regulatory circuits reveal variable modular perturbations across complex diseases. Nat Methods. 2016:13(4):366–370. 10.1038/nmeth.3799

Maxwell CA, Hartwig UA, Joseph CM, and Phillips DA. A chalcone and two related flavonoids released from alfalfa roots induce nod genes of Rhizobium meliloti. Plant Physiol. 1989:91(3):842–847. 10.1104/pp.91.3.842

Pankievicz VCS, Delaux P-M, Infante V, Hirsch HH, Rajasekar S, Zamora P, Jayaraman D, Calderon CI, Bennett A, and Ané J-M. Nitrogen fixation and mucilage production on maize aerial roots is controlled by aerial root development and border cell functions. Front Plant Sci. 2022:13:977056. 10.3389/fpls.2022.977056

Pereira WJ, Boyd J, Conde D, Triozzi PM, Balmant KM, Dervinis C, Schmidt HW, Boaventura-Novaes C, Chakraborty S, Knaack SA, et al. The single-cell transcriptome program of nodule development cellular lineages in Medicago truncatula. Cell Rep. 2024:43(2):113747. 10.1016/j.celrep.2024.113747

Riely BK, He H, Venkateshwaran M, Sarma B, Schraiber J, Ané J-M, and Cook DR. Identification of legume RopGEF gene families and characterization of a Medicago truncatula RopGEF mediating polar growth of root hairs: Medicago RopGEF gene family. Plant J. 2011:65(2):230–243. 10.1111/j.1365-313X.2010.04414.x

Ropitaux M, Bernard S, Schapman D, Follet-Gueye M-L, Vicré M, Boulogne I, and Driouich A. Root border cells and mucilage secretions of soybean, Glycine max (Merr) L.: Characterization and role in interactions with the Oomycete Phytophthora parasitica. Cells. 2020:9(10):2215. 10.3390/cells9102215

Roy S, Lagree S, Hou Z, Thomson JA, Stewart R, and Gasch AP. Integrated module and gene-specific regulatory inference implicates upstream signaling networks. PLoS Comput Biol. 2013:9(10):e1003252. 10.1371/journal.pcbi.1003252

Rudrappa T, Czymmek KJ, Paré PW, and Bais HP. Root-secreted malic acid recruits beneficial soil bacteria. Plant Physiol. 2008:148(3):1547–1556. 10.1104/pp.108.127613

Schindelin J, Arganda-Carreras I, Frise E, Kaynig V, Longair M, Pietzsch T, Preibisch S, Rueden C, Saalfeld S, Schmid B, et al. Fiji: an open-source platform for biological-image analysis. Nat Methods. 2012:9(7):676–682. 10.1038/nmeth.2019

Shulse CN, Cole BJ, Ciobanu D, Lin J, Yoshinaga Y, Gouran M, Turco GM, Zhu Y, O’Malley RC, Brady SM, et al. High-throughput single-cell transcriptome profiling of plant cell types. Cell Rep. 2019:27(7):2241–2247.e4. 10.1016/j.celrep.2019.04.054

Springer NM, Anderson SN, Andorf CM, Ahern KR, Bai F, Barad O, Barbazuk WB, Bass HW, Baruch K, Ben-Zvi G, et al. The maize W22 genome provides a foundation for functional genomics and transposon biology. Nat Genet. 2018:50(9):1282–1288. 10.1038/s41588-018-0158-0

Stein-O’Brien GL, Arora R, Culhane AC, Favorov AV, Garmire LX, Greene CS, Goff LA, Li Y, Ngom A, Ochs MF, et al. Enter the matrix: Factorization uncovers knowledge from omics. Trends Genet. 2018:34(10):790–805. 10.1016/j.tig.2018.07.003

Sudhakar Reddy P, Srinivas Reddy D, Sivasakthi K, Bhatnagar-Mathur P, Vadez V, and Sharma KK. Evaluation of sorghum [sorghum bicolor (L.)] reference genes in various tissues and under abiotic stress conditions for quantitative real-time PCR data normalization. Front Plant Sci. 2016:7:529. 10.3389/fpls.2016.00529

Tran TM, MacIntyre A, Hawes M, and Allen C. Escaping underground nets: Extracellular DNases degrade plant extracellular traps and contribute to virulence of the plant pathogenic bacterium Ralstonia solanacearum. PLoS Pathog. 2016:12(6):e1005686. 10.1371/journal.ppat.1005686

Van Deynze A, Zamora P, Delaux P-M, Heitmann C, Jayaraman D, Rajasekar S, Graham D, Maeda J, Gibson D, Schwartz KD, et al. Nitrogen fixation in a landrace of maize is supported by a mucilage-associated diazotrophic microbiota. PLoS Biol. 2018:16(8):e2006352. 10.1371/journal.pbio.2006352

Venado RE, Wilker J, Infante V, McLimans C, Robbins F, Phillips C, Calderón CI, Wallace JG, and Ané J-M. Aerial root formation in Oaxacan maize (Zea mays) landraces persists into the adult phase and is minimally affected by soil nitrogen and ambient humidity. Front Plant Sci. 2025a:16(1607733):1607733. 10.3389/fpls.2025.1607733

Venado RE, Wilker J, Pankievicz VCS, Infante V, MacIntyre A, Wolf ESA, Vela S, Robbins F, Fernandes-Júnior PI, Vermerris W, et al. Mucilage produced by aerial roots hosts diazotrophs that provide nitrogen in Sorghum bicolor. PLoS Biol. 2025b:23(3):e3003037. 10.1371/journal.pbio.3003037

Venado RE, Wilker J, Pankievicz V, Infante V, MacIntyre A, Wolf E, Vela S, Robbins F, Fernandes-Júnior PI, Vermerris W, et al. Mucilage produced by sorghum (Sorghum bicolor) aerial roots supports a nitrogen-fixing community. bioRxiv. 2023:2023.08.05.552127. 10.1101/2023.08.05.552127

Watson BS, Bedair MF, Urbanczyk-Wochniak E, Huhman DV, Yang DS, Allen SN, Li W, Tang Y, and Sumner LW. Integrated metabolomics and transcriptomics reveal enhanced specialized metabolism in Medicago truncatula root border cells. Plant Physiol. 2015:167(4):1699–1716. 10.1104/pp.114.253054

Webb BA, Compton KK, Del Campo JSM, Taylor D, Sobrado P, and Scharf BE. Sinorhizobium meliloti chemotaxis to multiple amino acids is mediated by the chemoreceptor McpU. Mol Plant Microbe Interact. 2017:30(10):770–777. 10.1094/MPMI-04-17-0096-R

Wei K, Chen J, Wang Y, Chen Y, Chen S, Lin Y, Pan S, Zhong X, and Xie D. Genome-wide analysis of bZIP-encoding genes in maize. DNA Res. 2012:19(6):463–476. 10.1093/dnares/dss026

Wen F, White GJ, VanEtten HD, Xiong Z, and Hawes MC. Extracellular DNA is required for root tip resistance to fungal infection. Plant Physiol. 2009:151(2):820–829. 10.1104/pp.109.142067

Wolf ESA, Vela S, Wilker J, Davis A, Robert M, Infante V, Venado RE, Voiniciuc C, Ané J-M, and Vermerris W. Identification of genetic and environmental factors influencing aerial root traits that support biological nitrogen fixation in sorghum. G3. 2024:14(3). 10.1093/g3journal/jkad285

Wolf FA, Angerer P, and Theis FJ. SCANPY: large-scale single-cell gene expression data analysis. Genome Biol. 2018:19(1):15. 10.1186/s13059-017-1382-0

Zhang S, Pyne S, Pietrzak S, Halberg S, McCalla SG, Siahpirani AF, Sridharan R, and Roy S. Inference of cell type-specific gene regulatory networks on cell lineages from single cell omic datasets. Nat Commun. 2023:14(1):3064. 10.1038/s41467-023-38637-9

Zhang T, Lv W, Zhang H, Ma L, Li P, Ge L, and Li G. Genome-wide analysis of the basic Helix-Loop-Helix (bHLH) transcription factor family in maize. BMC Plant Biol. 2018:18(1):235. 10.1186/s12870-018-1441-z

Zhu T, Wang H, Wang Y, Yang X, Fang S, Li W, Rui W, Wang B, Tian Y, Sun D, et al. Transcriptome-wide association study identifies the ZmbZIP89-ZmSAUR21 module as a key regulator of root growth and drought tolerance in maize. J Integr Plant Biol. 2026:(jipb.70220). 10.1111/jipb.70220

