## Supplementary Text for "Single-Cell Omics to Uncover Cell-Type-Specific Gene Regulatory Networks Driving Mucilage Production in Specialized Roots"

### Supplemental Figures

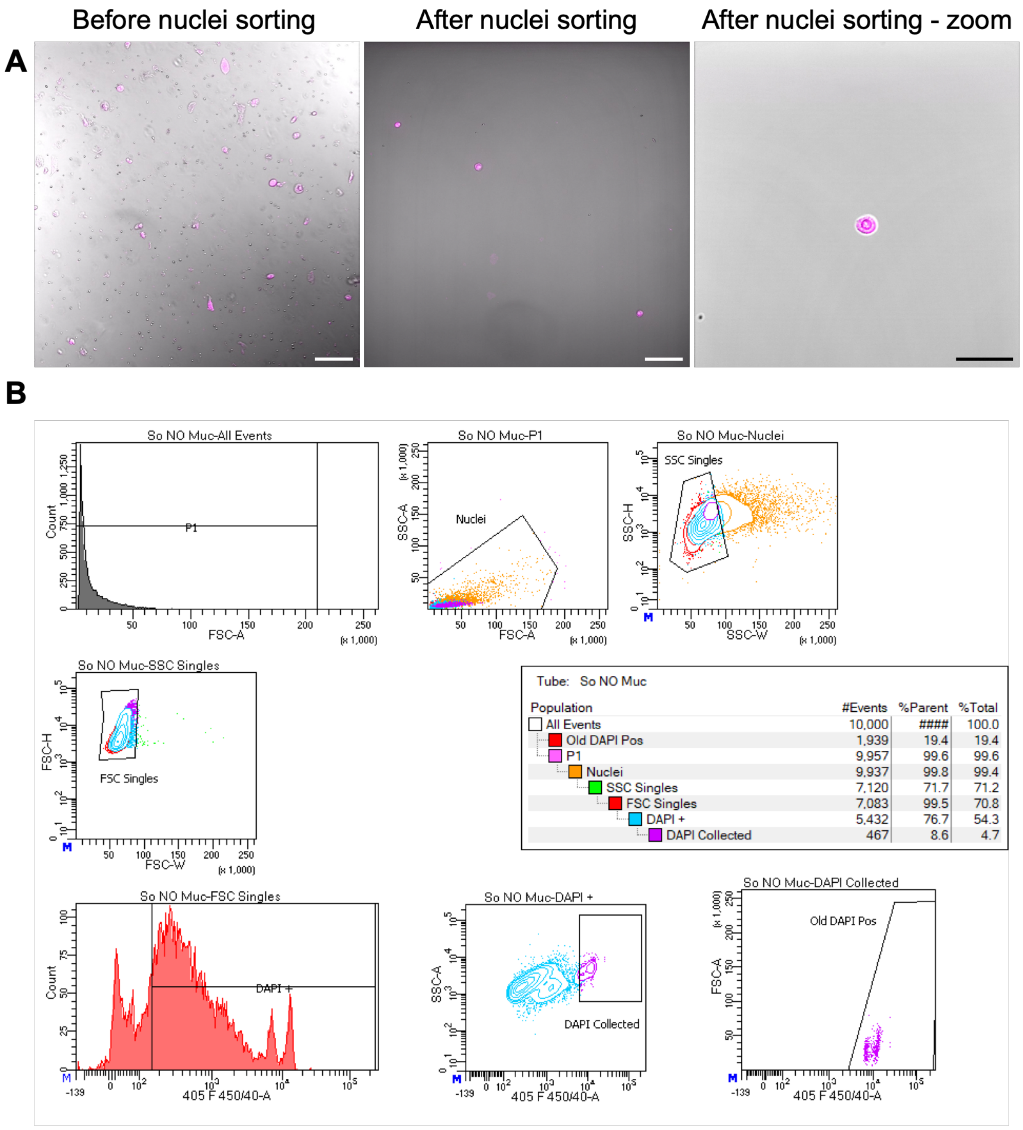

**Supplemental Figure 1: Fluorescence-Activated Nuclei Sorting (FANS).** Isolation of nuclei requires the removal of all plant debris co-isolated with the nuclei, such as chloroplasts and other particles, especially considering the nature of aerial roots. A) After nuclei isolation, the samples contained various plant particles that were removed using FANS. After nuclei isolation, samples were stained with DAPI (magenta) and sorted using FANS. Finally, a zoomed-in panel shows nuclear integrity, a critical parameter for successful snRNA-seq. Scale bars is 50 µm and 25 µm for the zoomed-in panel. B) FANS used different filters to select single nuclei. First, the total number of cells was quantified. Next, all DAPI-stained particles were selected, followed by light scattering analysis using side scatter (SSC) and forward scatter (FSC). Finally, a total of 40,000 nuclei were selected for processing with the 10× Genomics microfluidic chips.

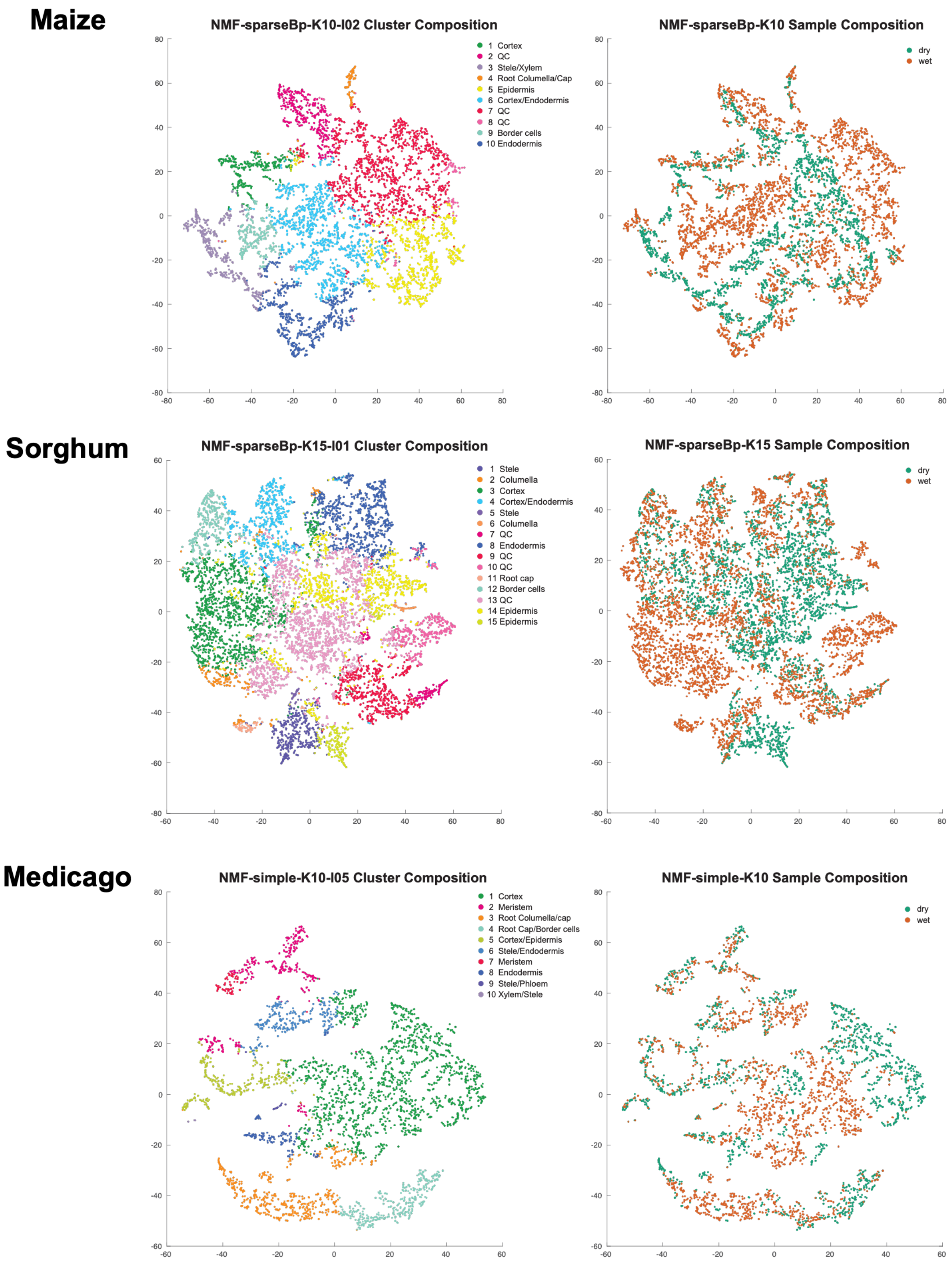

**Supplemental Figure 2: Clusters identified using NMF in maize, sorghum, and Medicago.** The snRNA-seq data were visualized using a t-SNE plot generated from the combined dataset, with the proportion of cells from each condition displayed alongside. We identified ten clusters in maize, fifteen in sorghum, and ten in Medicago. In the distribution plots, dry conditions are shown in green and wet conditions in brown.

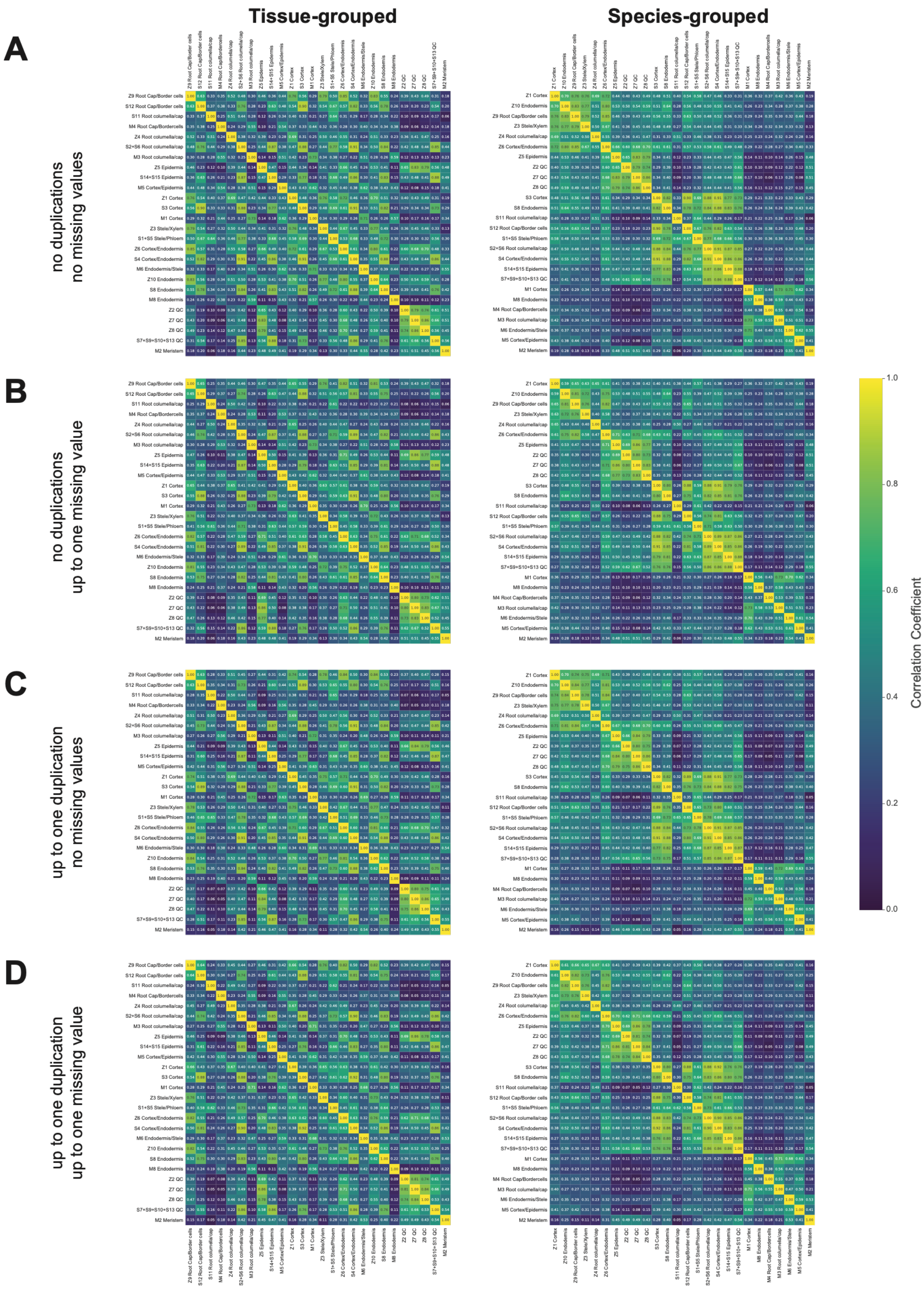

**Supplemental Figure 3: Cross-species comparison of pseudo-bulk gene expression profiles across cell clusters.** Pearson correlation matrices comparing pseudo-bulk expression profiles of cell clusters from Medicago (M), maize (Z), and sorghum (S). For each orthogroup (OG) set: A) no duplications, no missing values; B) no duplications, up to one missing value; C) up to one duplication, no missing values; D) up to one duplication, up to one missing value, pseudo-bulk profiles were generated by averaging expression across all cells assigned to each cluster. Correlations were then computed in an all-by-all manner between clusters using vectors containing the pseudo-bulk expression values of all genes in the OG set. Each correlation matrix is shown with two orderings: tissue-grouped (left), where clusters are arranged by shared cell-type annotations across species, and species-grouped (right), where clusters are arranged by species.

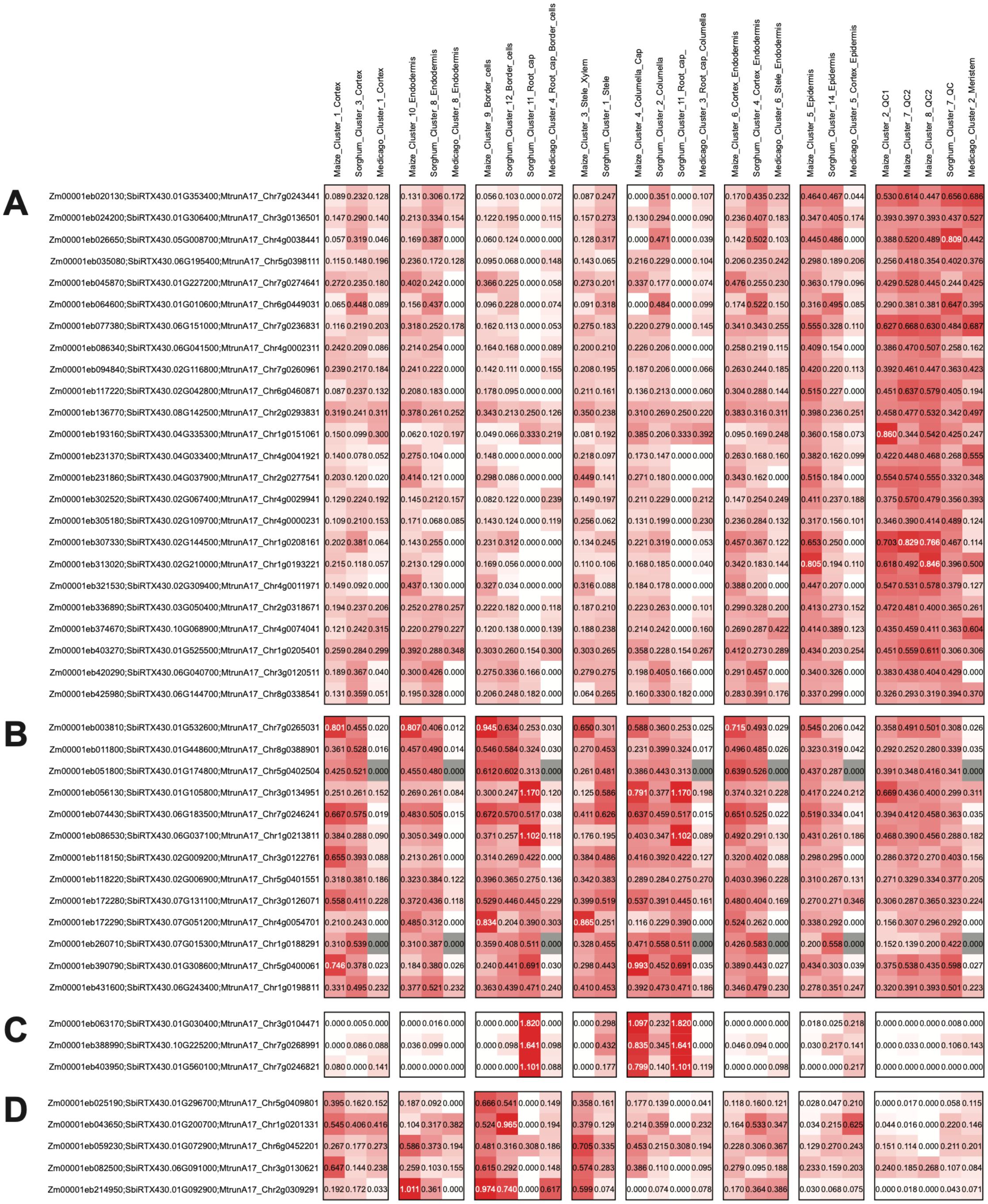

**Supplemental Figure 4: Cross-species conservation of orthogroup pseudo-bulk expression patterns.** Heatmaps show pseudo-bulk expression profiles of selected orthogroups from the set without duplications and without missing values across Medicago, maize, and sorghum cell clusters. Genes were grouped according to their cross-species pseudo-bulk expression patterns using k-means clustering, and four representative clusters with distinct expression patterns are shown in panels A) – D). Columns represent cell clusters from all three species, grouped by similar cell-type annotations; rows correspond to orthogroups. Each row label lists the corresponding orthologous genes from maize, sorghum, and Medicago.

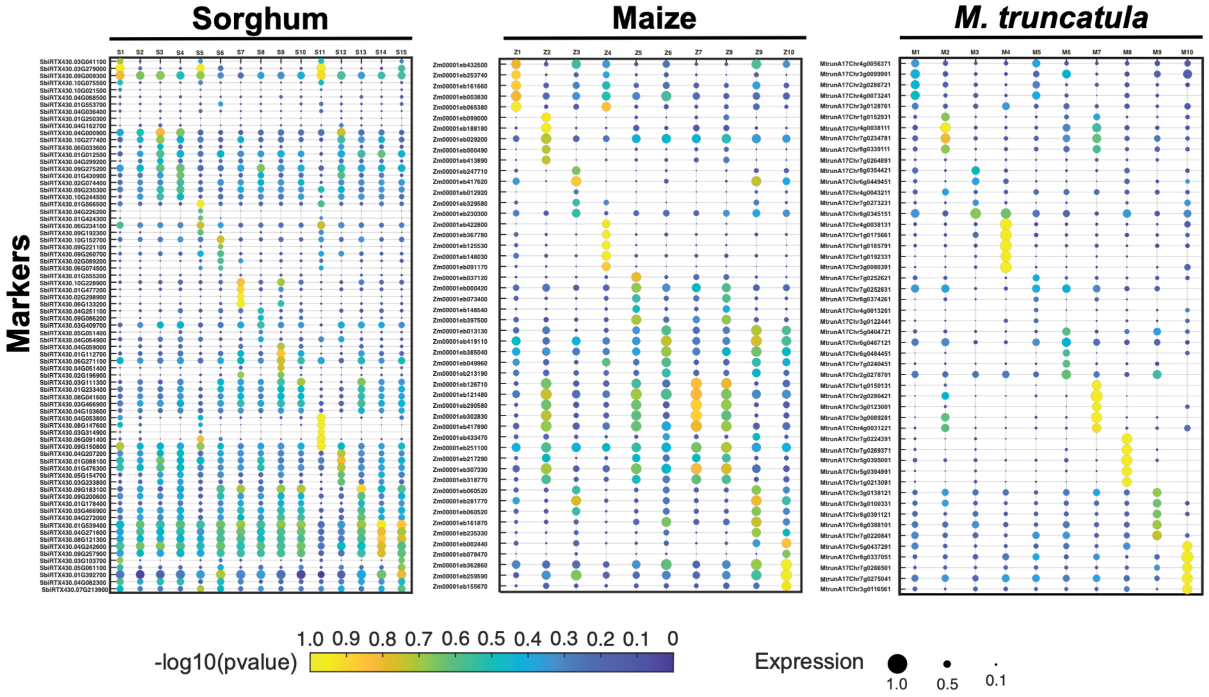

**Supplemental Figure 5: snRNA-seq cluster-specific markers. A)** Top differentially expressed genes (DEGs) identified for each cluster, showing their relative expression across all clusters.

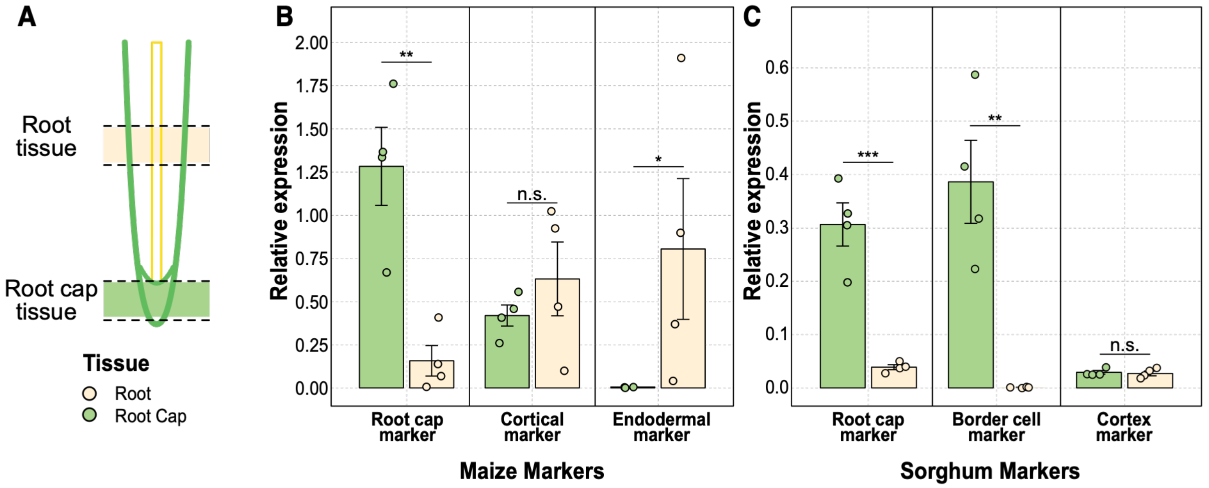

**Supplemental Figure 6:** Gene expression analysis of selected markers was performed to validate cluster annotation in maize and sorghum using RT-qPCR to quantify transcript abundance. For maize, the markers analyzed included: root cap - Zm00001eb163560, cortex - Zm0001eb027760, and endodermis - Zm00001eb005460. For sorghum, the markers tested were root cap - SbiRTX430.04G115800, border cells - SbiRTX430.05G002900, and cortex - SbiRTX430.01G385000. A) Total RNA was extracted from either the aerial root cap or the elongation zone of aerial roots. B) Relative expression of cluster markers in two distinct aerial root tissues of maize. Expression was normalized using housekeeping genes ZmActin and ZmUbiquitin. C) Relative expression of cluster markers in two distinct aerial root tissues of sorghum. Expression was normalized using SbTubulin and SbGAPDH. Each dot represents an independent biological replicate. Statistical significance between tissue types was assessed using Student's t-test in R (v4.4.0). Significant levels * < 0.05, ** <0.01, *** < 0.001 and n.s non-significant.

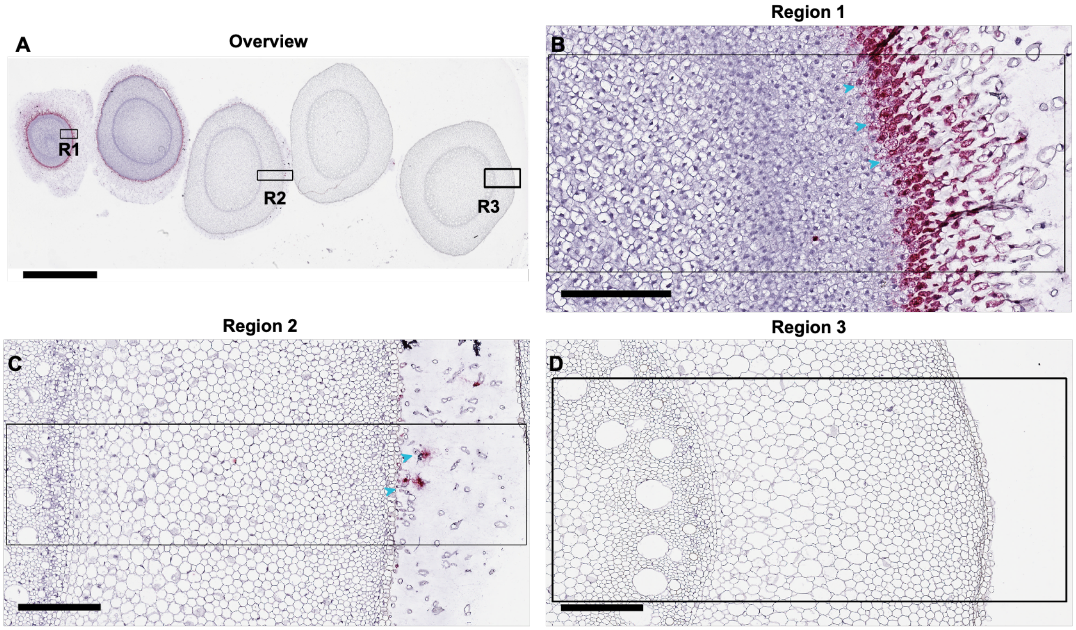

**Supplemental Figure 7*: Maize root cap marker.*** *In situ* hybridization of the root cap marker Zm00001eb163560 in various sections of the aerial root of maize. A) Overview image showing the hybridization signal across different root sections. Boxes indicate regions shown in higher magnification. Scale bar: 4 mm. B) Magnified view of the root tip (first region); arrows indicate marker expression in root cap cells, especially those becoming border cells. Scale bar: 200 µm C) Marker expression remains detectable in the third root section (blue arrows). Scale bar: 200 µm D) In the fifth section, signal is no longer detected in any cell layers. Scale bar: 200 µm

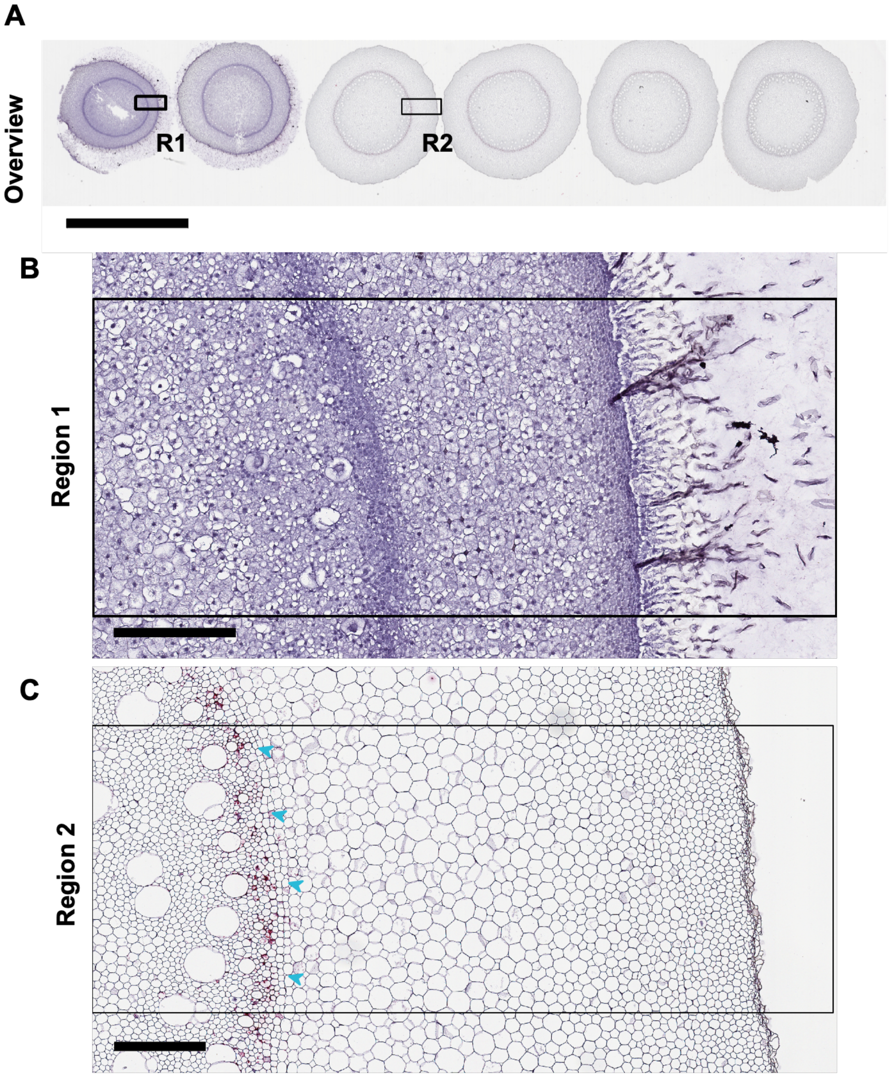

**Supplemental Figure 8: Supplemental Figure 8. Maize endodermis marker**. In situ hybridization of the endodermis marker Zm00001eb005460 in various sections of the aerial root of maize. A) Overview image showing the hybridization signal across different root sections. Boxes indicate regions shown in higher magnification. Scale bar: 5 mm. B) Magnified view of the root tip (first region) without probe hybridization. Scale bar: 200 µm. C) Marker expression is observed in the elongation zone, specifically localized to the endodermal cell layer. (blue arrows). Scale bar: 200 µm.

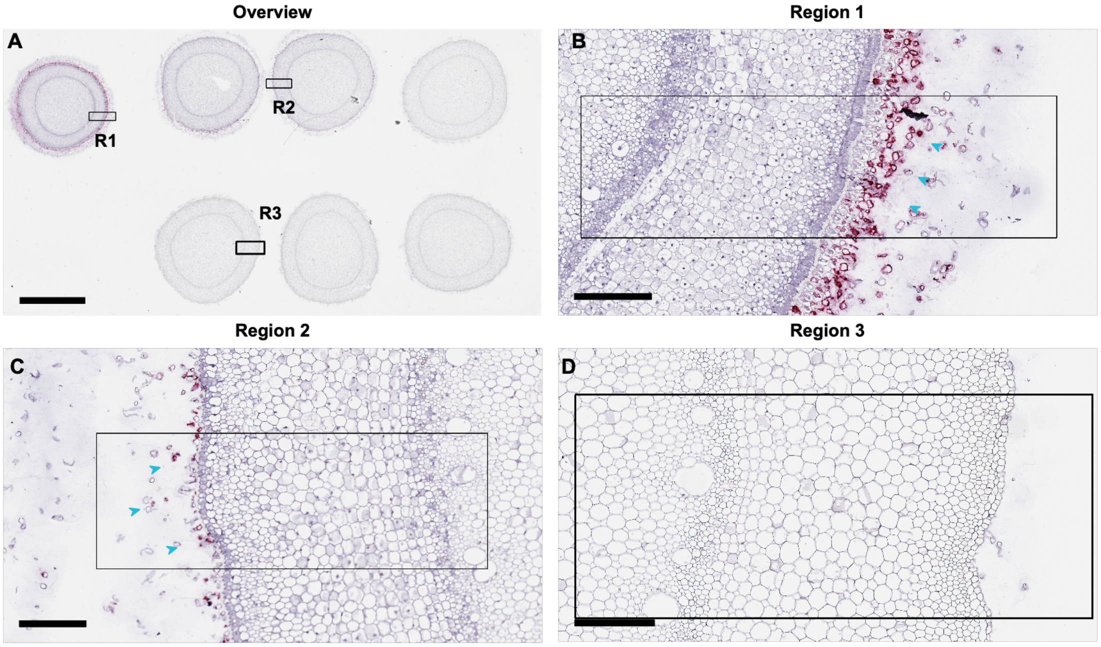

**Supplemental Figure 9: Sorghum border cell marker.** In situ hybridization of the border cell marker SbiRTX430.05G002900 in various sections of the aerial root of sorghum. A) Overview image showing the hybridization signal across different root sections. Boxes indicate regions shown in higher magnification. Scale bar: 3 mm. B) Magnified view of the root tip (first region); arrows indicate marker expression in border cells. Scale bar: 200 µm. C) Marker expression remains detectable in the third root section (blue arrows). Scale bar: 200 µm D) In the fifth section, border cells are no longer present, and no signal is detected. Scale bar: 200 µm

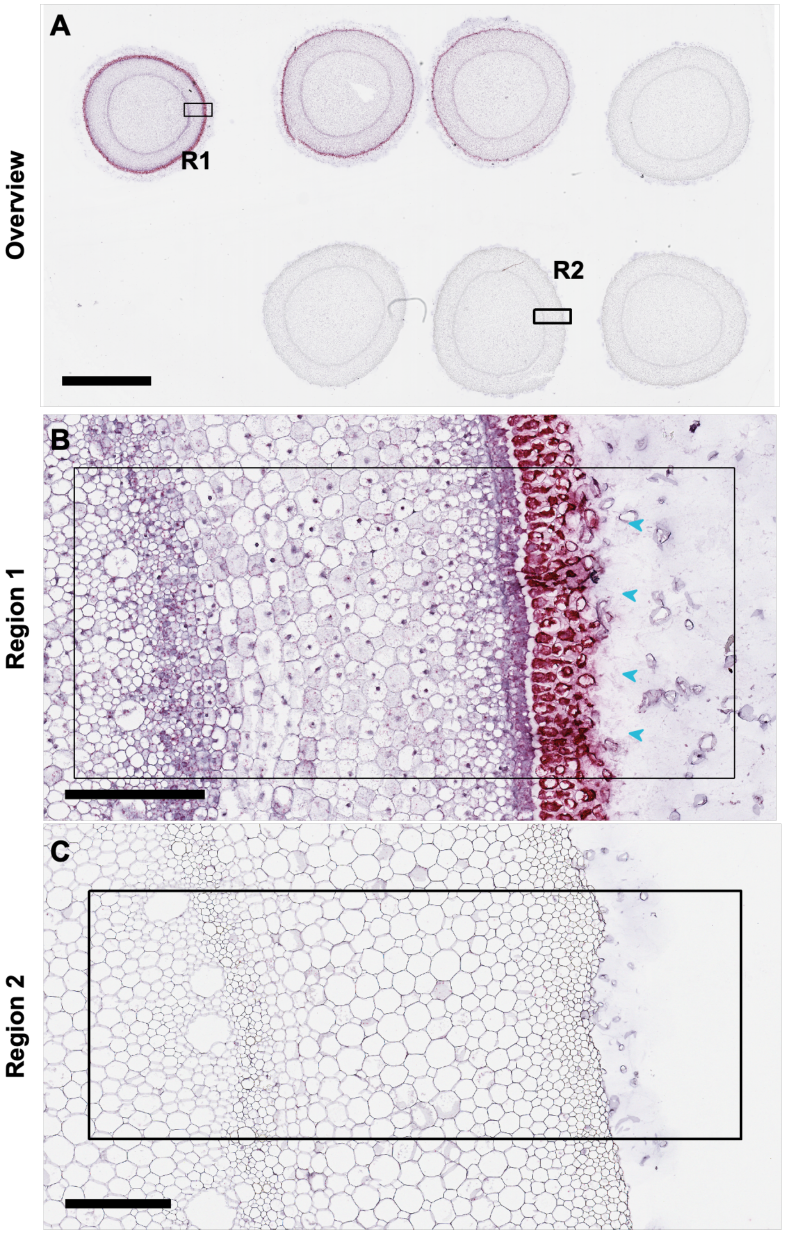

**Supplemental Figure 10:** In situ hybridization of the root cap marker SbiRTX430.03G383000 in various sections of the aerial root of sorghum. A) Overview image showing the hybridization signal across different root sections. Boxes indicate regions shown in higher magnification. Scale bar: 5 mm. B) Magnified view of the root tip (first region) with high expression in the root cap cells (blue arrows). Hybridization signal is no longer detected beyond the fourth tissue section. Scale bar: 200 µm.

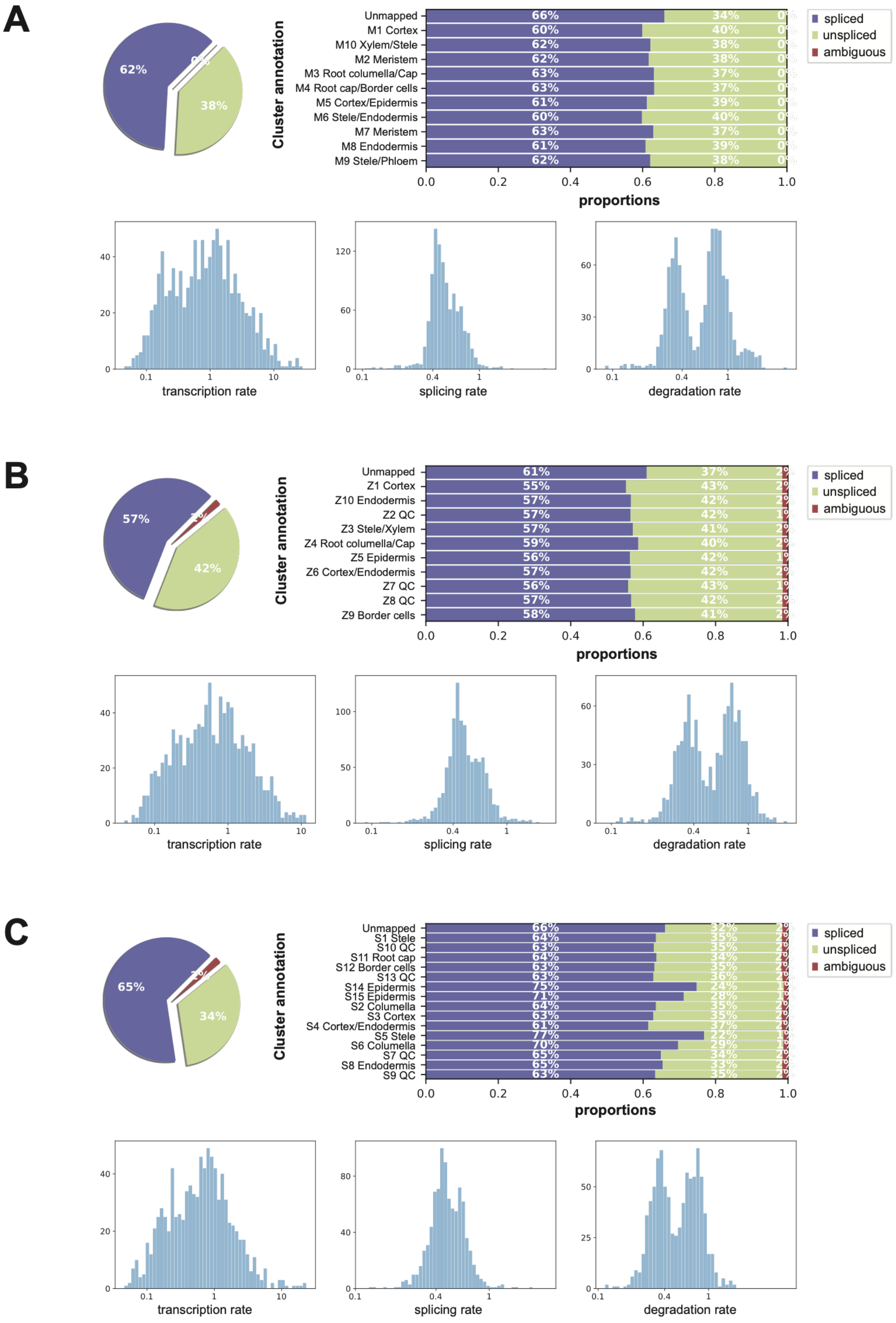

**Supplemental Figure 11: Summary statistics of RNA velocity model parameters.** For each species, **A)** Medicago, **B)** maize**, C)** sorghum, the upper panels show the proportion of spliced, unspliced, and ambiguous transcript counts across all cells (left) and across annotated cell clusters (right). The lower panels show the distributions of kinetic parameters inferred by the scVelo dynamical RNA velocity model, including transcription rate, splicing rate, and degradation rate across genes.

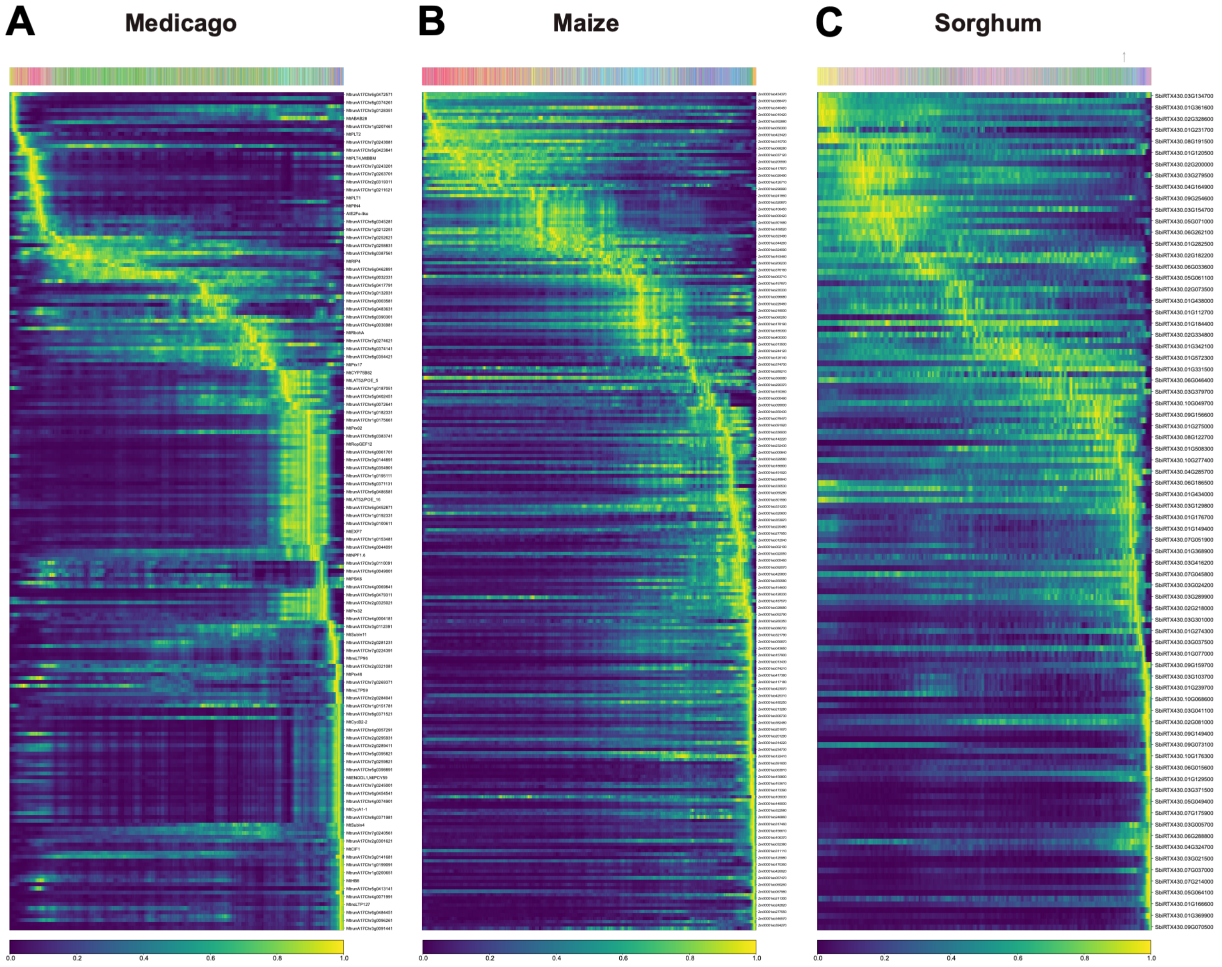

**Supplemental Figure 12: Expression dynamics of known marker genes along latent time.** A–C) Heatmaps showing the expression of known marker genes ordered by latent time in (A) Medicago, (B) maize, and (C) sorghum. Columns represent individual cells arranged from early to late latent time, and rows represent genes. Expression values are scaled from low to high as indicated by the color bar. The annotation bar above each heatmap indicates cluster identity.

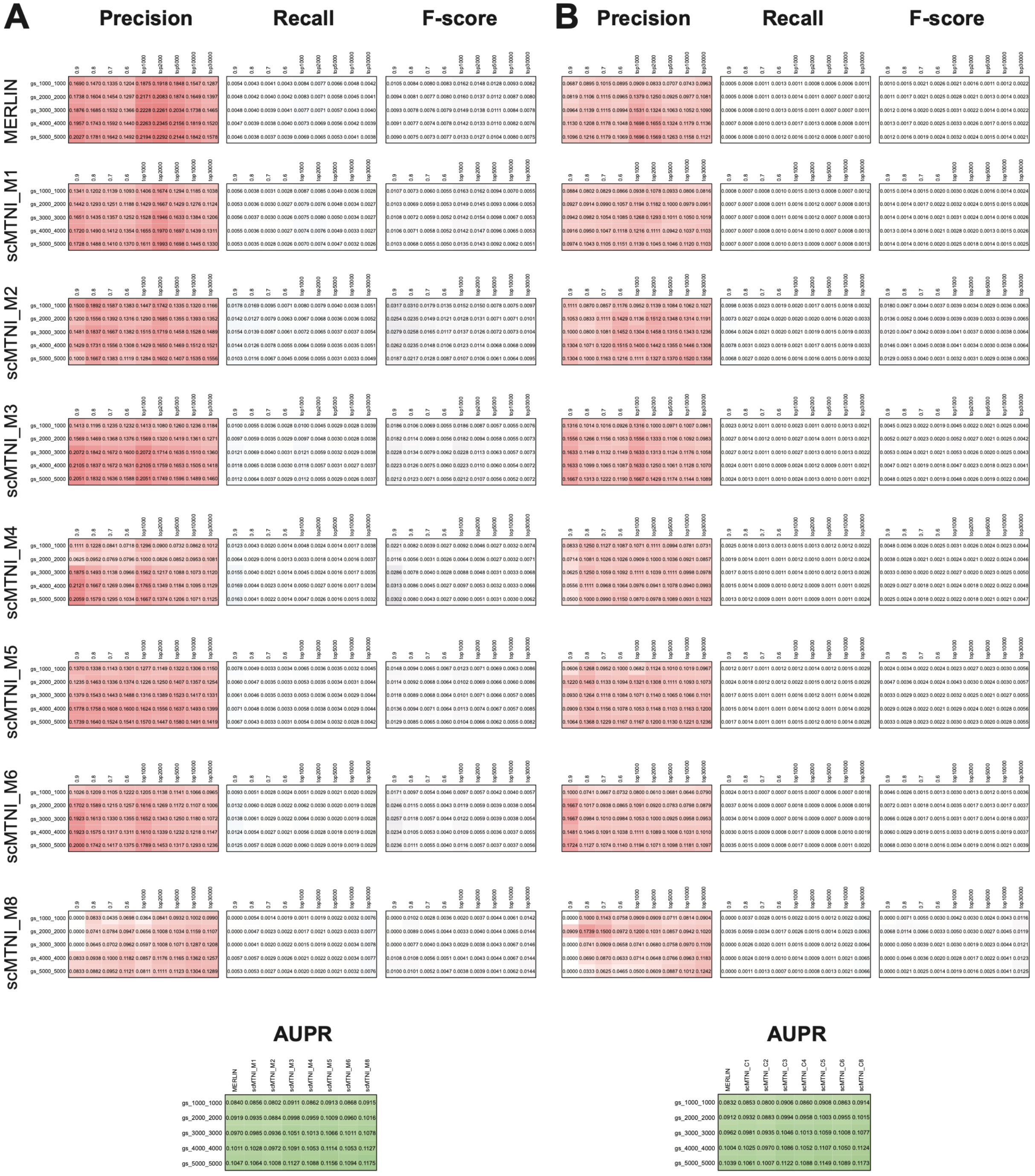

**Supplemental Figure 13: Evaluation of Medicago MERLIN and scMTNI cluster-specific networks against ATAC-seq derived regulatory gold standards.** Each row shows early precision, recall, F-score, and area under the precision-recall (AUPR) for Medicago regulatory networks inferred using MERLIN and scMTNI (cluster-specific networks). Precision, recall, and F-score were computed for the top inferred edges selected either by confidence thresholds (0.6, 0.7, 0.8, 0.9) or by the number of highest-scoring edges (top 1000, 2000, 5000, 10000, and 30000). Performance was evaluated against ATAC-seq-derived gold standards from GSE101482 using promoter regions defined as ±1 kb, ±2 kb, ±3 kb, ±4 kb, and ±5 kb around transcription starts sites (TSS). AUPR values were computed using the complete inferred networks. **A)** Evaluation of the inferred regulatory networks. **B)** Evaluation of randomized networks (edge-shuffled) used as a baseline control.

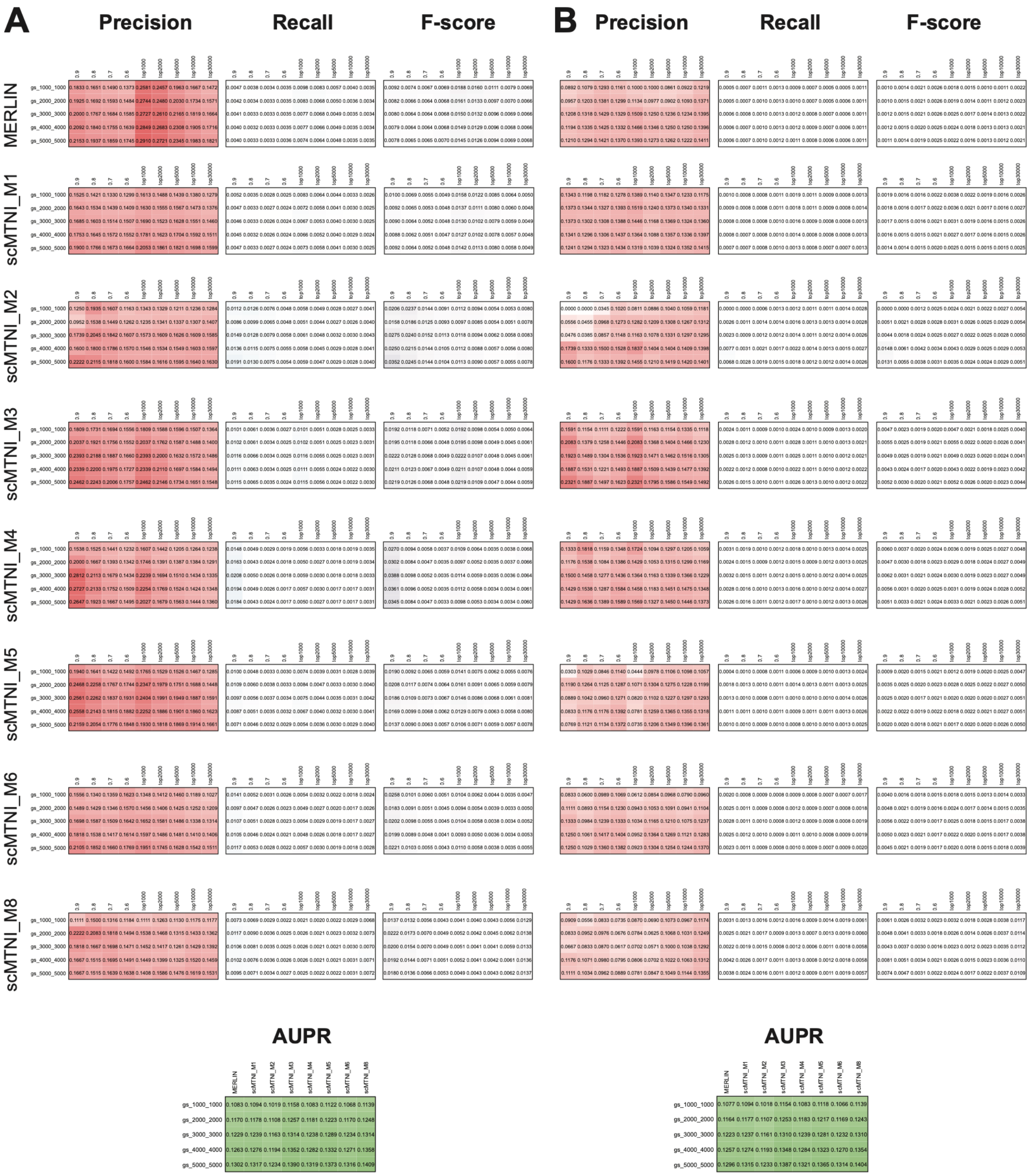

**Supplemental Figure 14: Evaluation of Medicago MERLIN and scMTNI cluster-specific networks against ATAC-seq derived regulatory gold standards.** Performance was evaluated against ATAC-seq-derived gold standards from GSE154845.

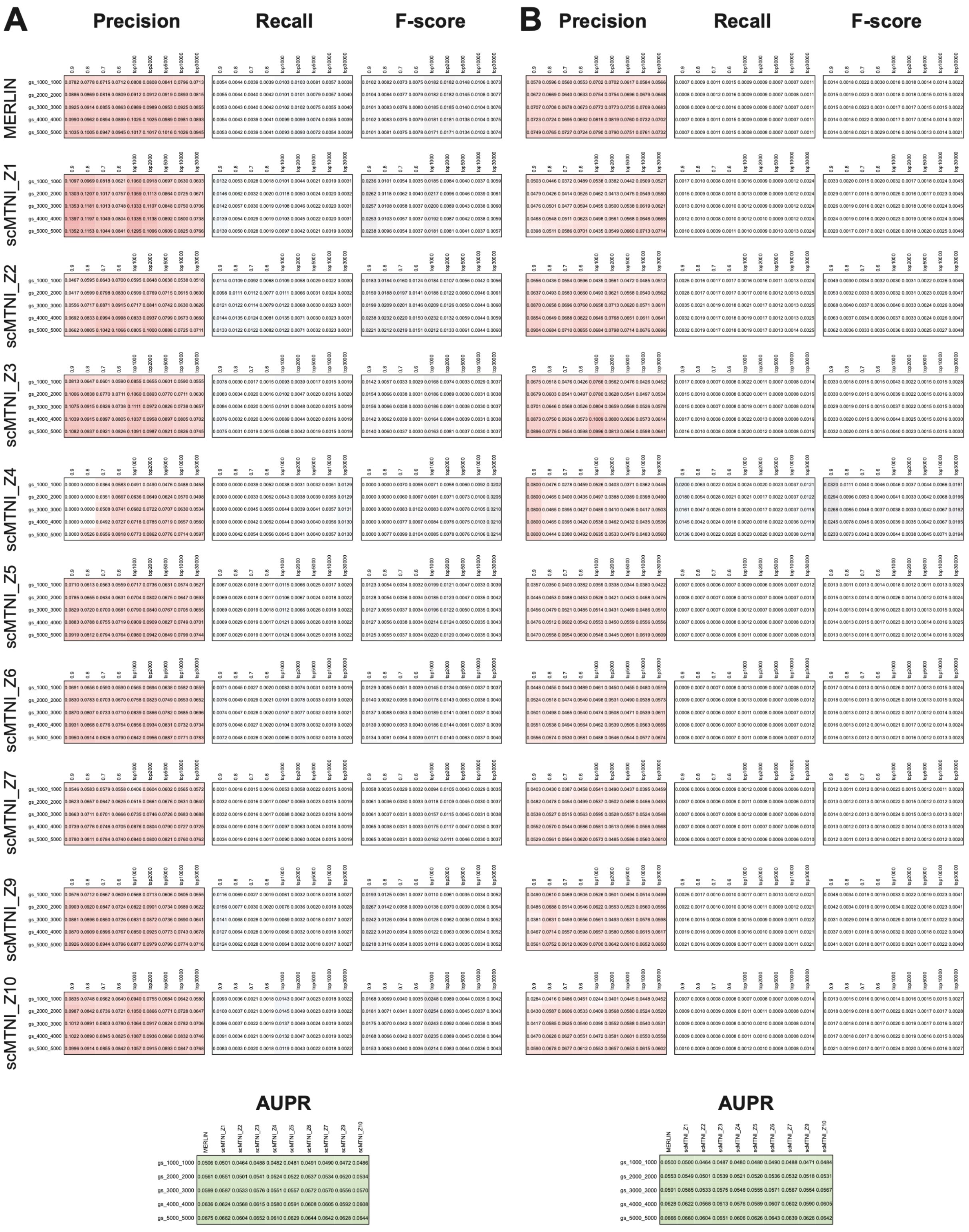

**Supplemental Figure 15: Evaluation of maize MERLIN and scMTNI cluster-specific networks against ATAC-seq derived regulatory gold standard.** Each row shows early precision, recall, F-score, and area under the precision-recall (AUPR) for maize regulatory networks inferred using MERLIN and scMTNI (cluster-specific networks). Precision, recall, and F-score were computed for the top inferred edges selected either by confidence thresholds (0.6, 0.7, 0.8, 0.9) or by the number of highest-scoring edges (top 1000, 2000, 5000, 10000, and 30000). Performance was evaluated against an ATAC-seq-derived gold standard from GSE152046, using promoter regions defined as ±1 kb, ±2 kb, ±3 kb, ±4 kb, and ±5 kb around transcription starts sites (TSS). AUPR values were computed using the complete inferred networks**. A)** Evaluation of the inferred regulatory networks. **B)** Evaluation of randomized networks (edge-shuffled) used as a baseline control.

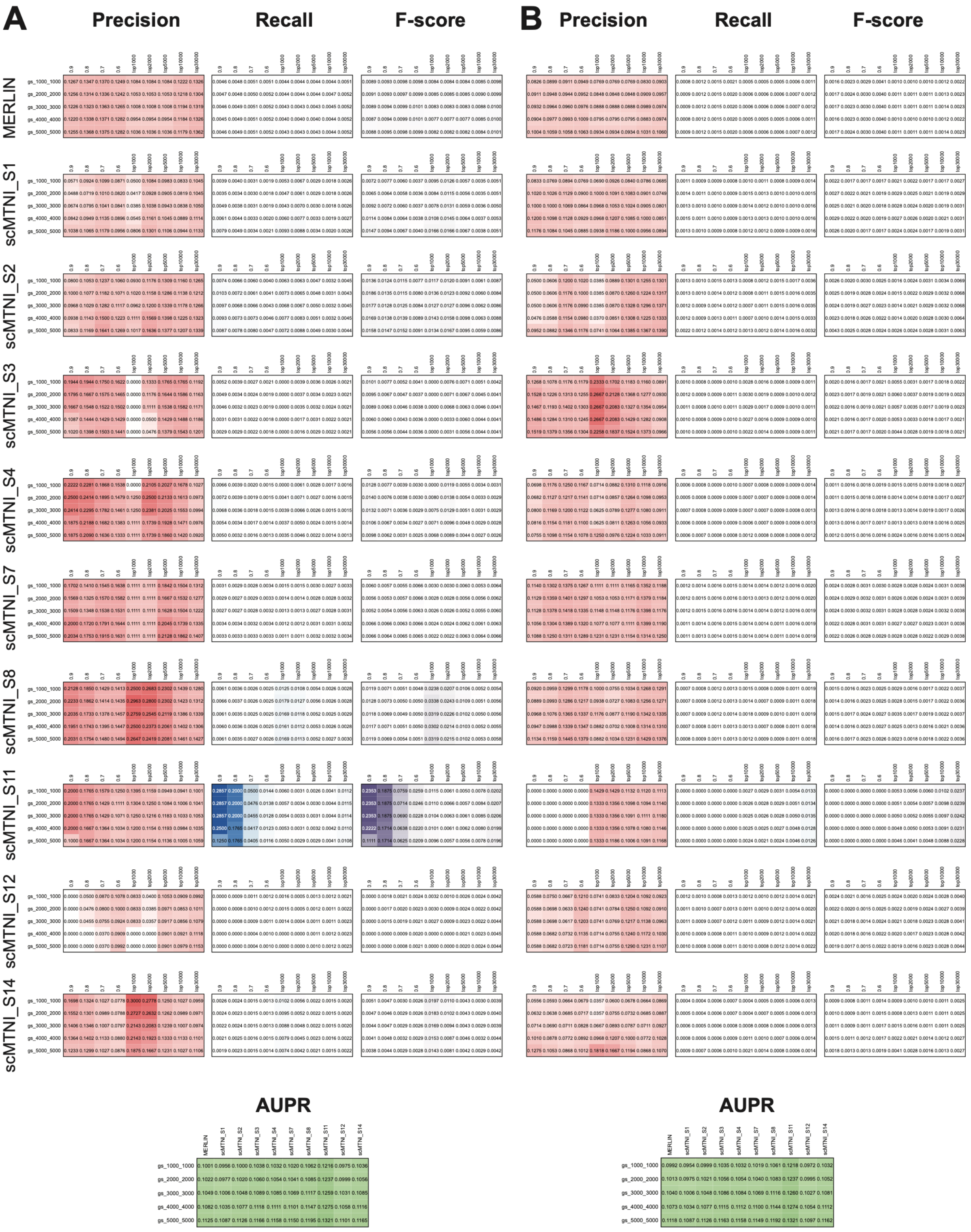

**Supplemental Figure 16: Evaluation of sorghum MERLIN and scMTNI cluster-specific networks against a ChIP-seq-derived regulatory gold standard.** Each row shows early precision, recall, F-score, and area under the precision-recall (AUPR) for sorghum regulatory networks inferred using MERLIN and scMTNI (cluster-specific networks). Precision, recall, and F-score were computed for the top inferred edges selected either by confidence thresholds (0.6, 0.7, 0.8, 0.9) or by the number of highest-scoring edges (top 1000, 2000, 5000, 10000, and 30000). Performance was evaluated against an H3K27ac ChIP-seq-derived gold standard from PRJNA952350, using promoter regions defined as ±1 kb, ±2 kb, ±3 kb, ±4 kb, and ±5 kb around transcription starts sites (TSS). AUPR values were computed using the complete inferred networks. **A)** Evaluation of the inferred regulatory networks. **B)** Evaluation of randomized networks (edge-shuffled) used as a baseline control.

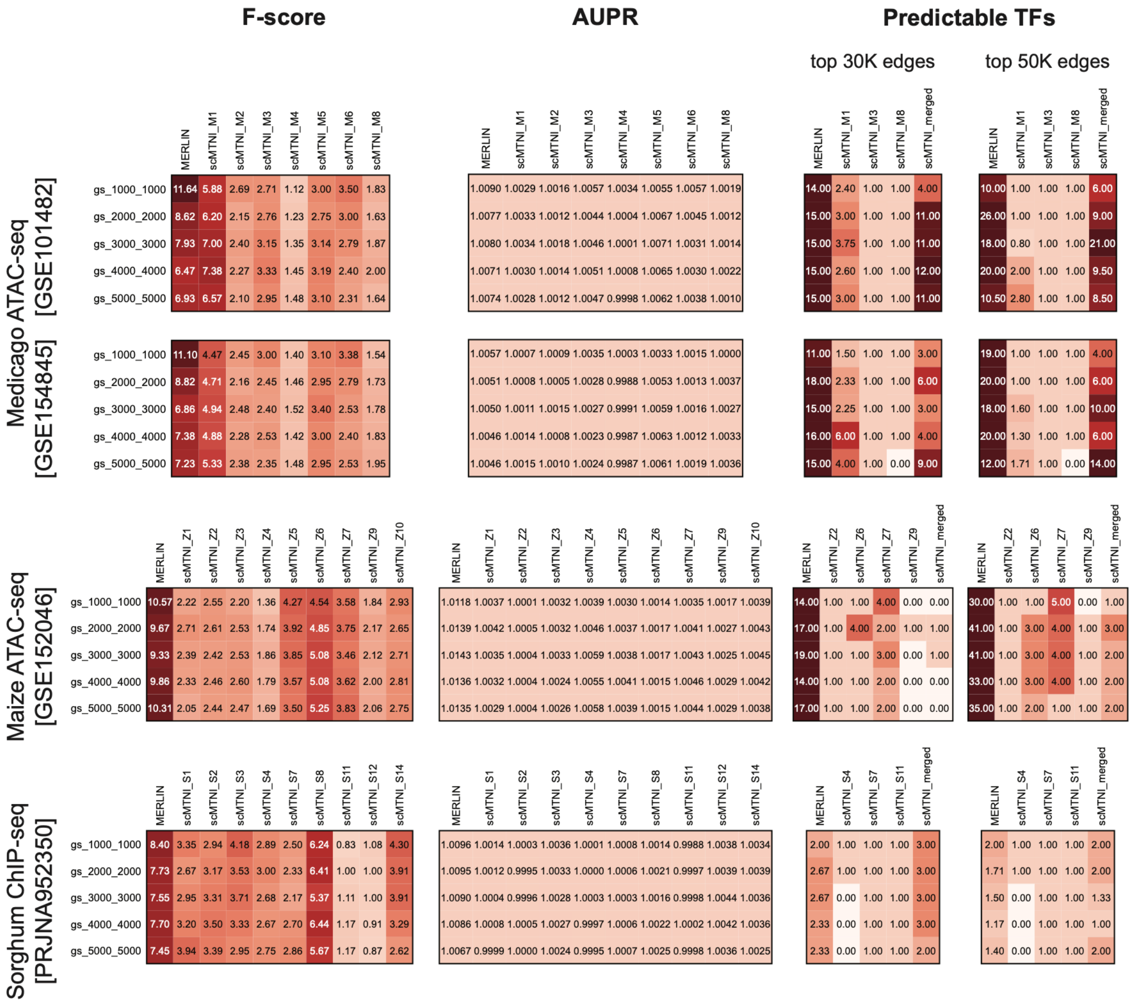

***Supplemental Figure 17:*** **Summary evaluation of inferred regulatory networks across species.** Heatmaps show fold enrichment of network performance metrics relative to randomized networks for MERLIN and scMTNI cluster-specific models. For F-score and AUPR, enrichment was calculated using the top 5000 inferred edges from each network against gold standards from Medicago ATAC-seq (GSE101482, GSE154845), maize ATAC-seq (GSE152046), and sorghum ChIP-seq (PRJNA952350). The Predictable TFs panels show fold enrichment in the number of transcription factors with non-zero enrichment scores compared to randomized networks, using the top 30000 edges (left panel) and top 50000 edges (right panel) from each inferred network. Rows correspond to gold standards and promoter region definitions used to construct them, and columns correspond to the evaluated networks (MERLIN, cluster-specific and merged scMTNI networks).

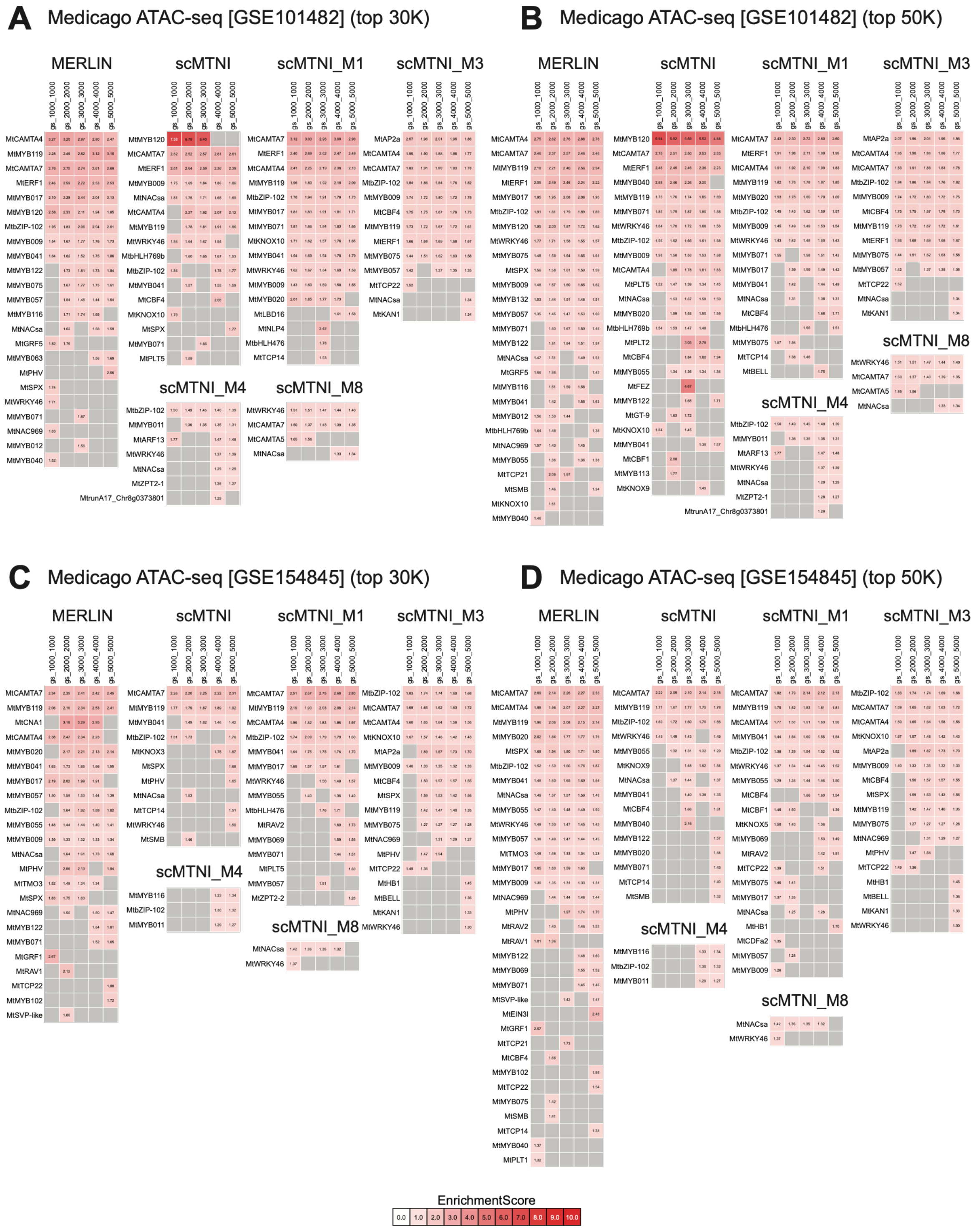

***Supplemental Figure 18:*** **Predictable transcription factors in Medicago MERLIN and scMTNI cluster-specific networks evaluated using ATAC-seq-derived gold standards.** Heatmaps show the enrichment of ATAC-seq-derived target sets for individual transcription factors (TFs) in the inferred regulatory networks. Rows correspond to TFs, and columns represent promoter regions used to construct the gold standard, defined as ±1 kb, ±2 kb, ±3 kb, ±4 kb, and ±5 kb around transcription start sites (TSS). Color intensity indicates the enrichment score. Grey cells indicate TFs not predicted by the corresponding network. **A, B)** Networks constructed from the A) top 30000 and B) top 50000 edges evaluated against the ATAC-seq gold standard from GSE101482. **C, D)** Networks constructed from the **C)** top 30000 and **D)** top 50000 edges evaluated against the ATAC-seq gold standard from GSE154845.

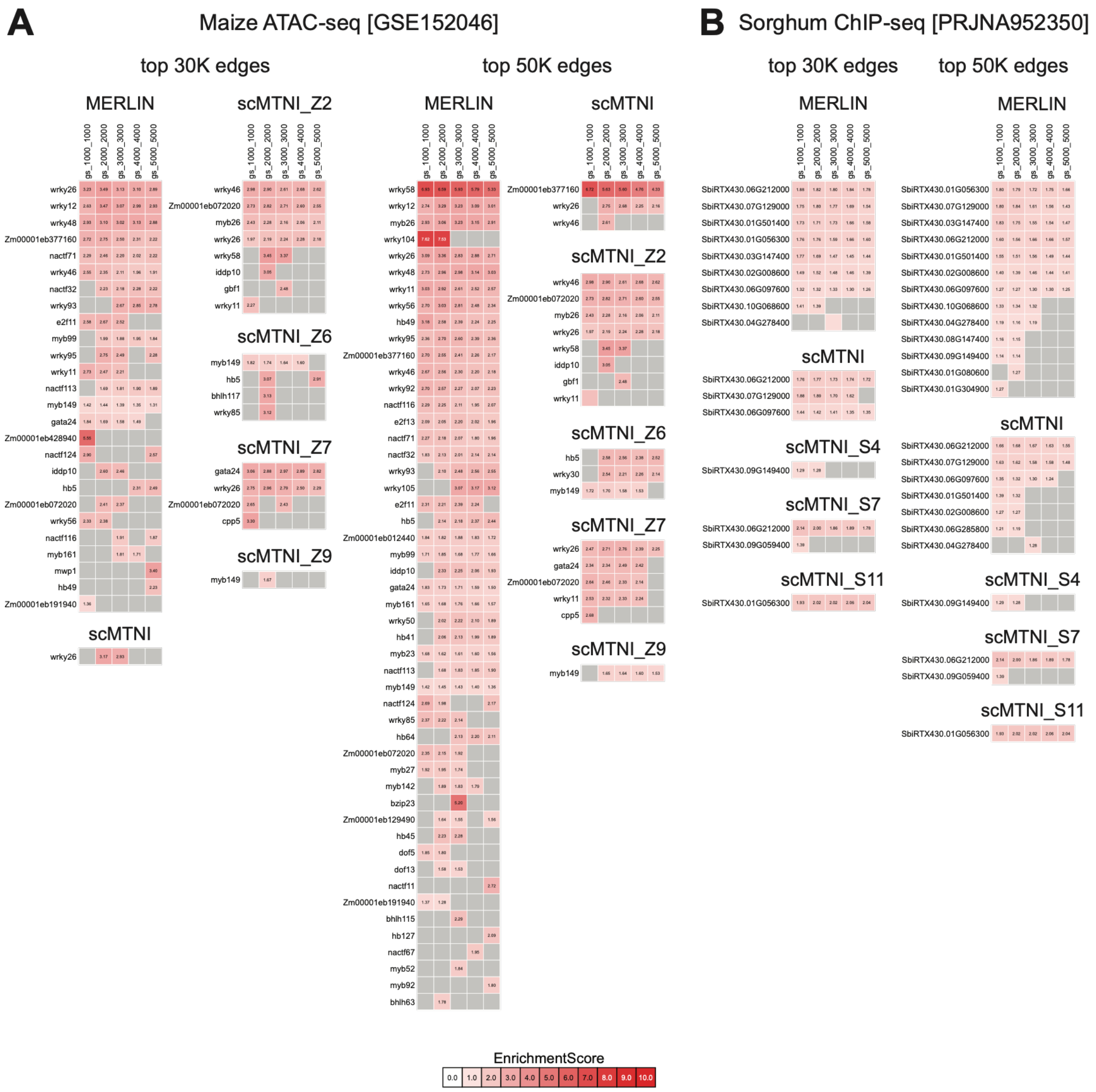

**Supplemental Figure 19:** ***Predictable transcription factors in maize and sorghum MERLIN and scMTNI cluster-specific networks evaluated using gold standards. A, B)*** *Heatmaps show enrichment of gold-standard target sets for individual transcription factors (TFs) in the inferred regulatory networks. Rows correspond to TFs, and columns represent promoter regions used to construct the gold standard, defined as ±1 kb, ±2 kb, ±3 kb, ±4 kb, and ±5 kb around transcription start sites (TSS). Color intensity indicates the enrichment score. Grey cells indicate TFs that were not identified as predictable by the corresponding network.* ***A)*** *Maize networks constructed from the top 30000 and top 50000 edges evaluated against the ATAC-seq gold standard from GSE152046.* ***B)*** *Sorghum networks constructed from the top 30000 and top 50000 edges evaluated against the ChIP-seq-derived gold standard from PRJNA952350.*

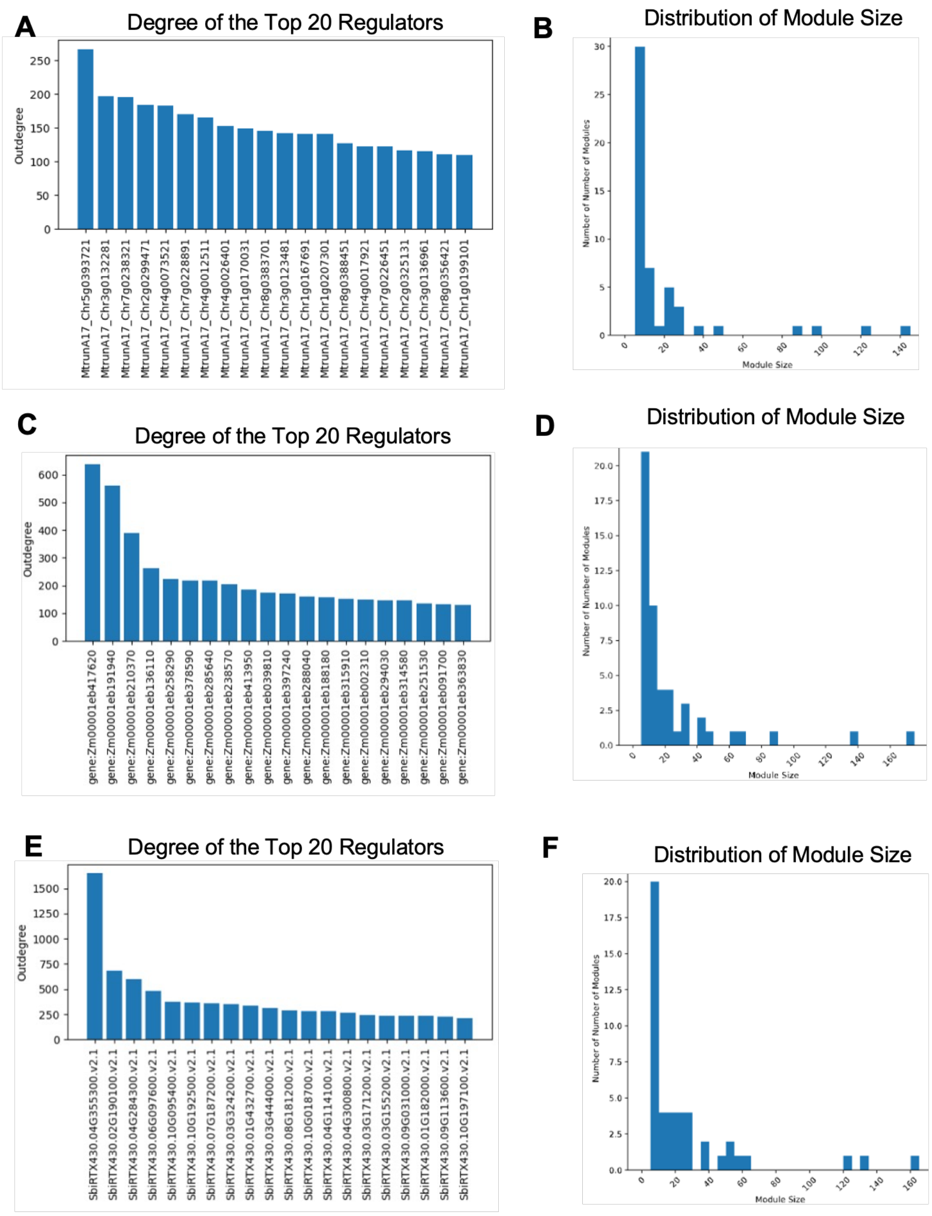

**Supplemental Figure 20: Predictable transcription factors in maize and sorghum MERLIN and scMTNI cluster-specific networks evaluated using gold standards. A, B)** Heatmaps show enrichment of gold-standard target sets for individual transcription factors (TFs) in the inferred regulatory networks. Rows correspond to TFs, and columns represent promoter regions used to construct the gold standard, defined as ±1 kb, ±2 kb, ±3 kb, ±4 kb, and ±5 kb around transcription start sites (TSS). Color intensity indicates the enrichment score. Grey cells indicate TFs that were not identified as predictable by the corresponding network. **A)** Maize networks constructed from the top 30000 and top 50000 edges evaluated against the ATAC-seq gold standard from GSE152046. **B)** Sorghum networks constructed from the top 30000 and top 50000 edges evaluated against the ChIP-seq-derived gold standard from PRJNA952350.

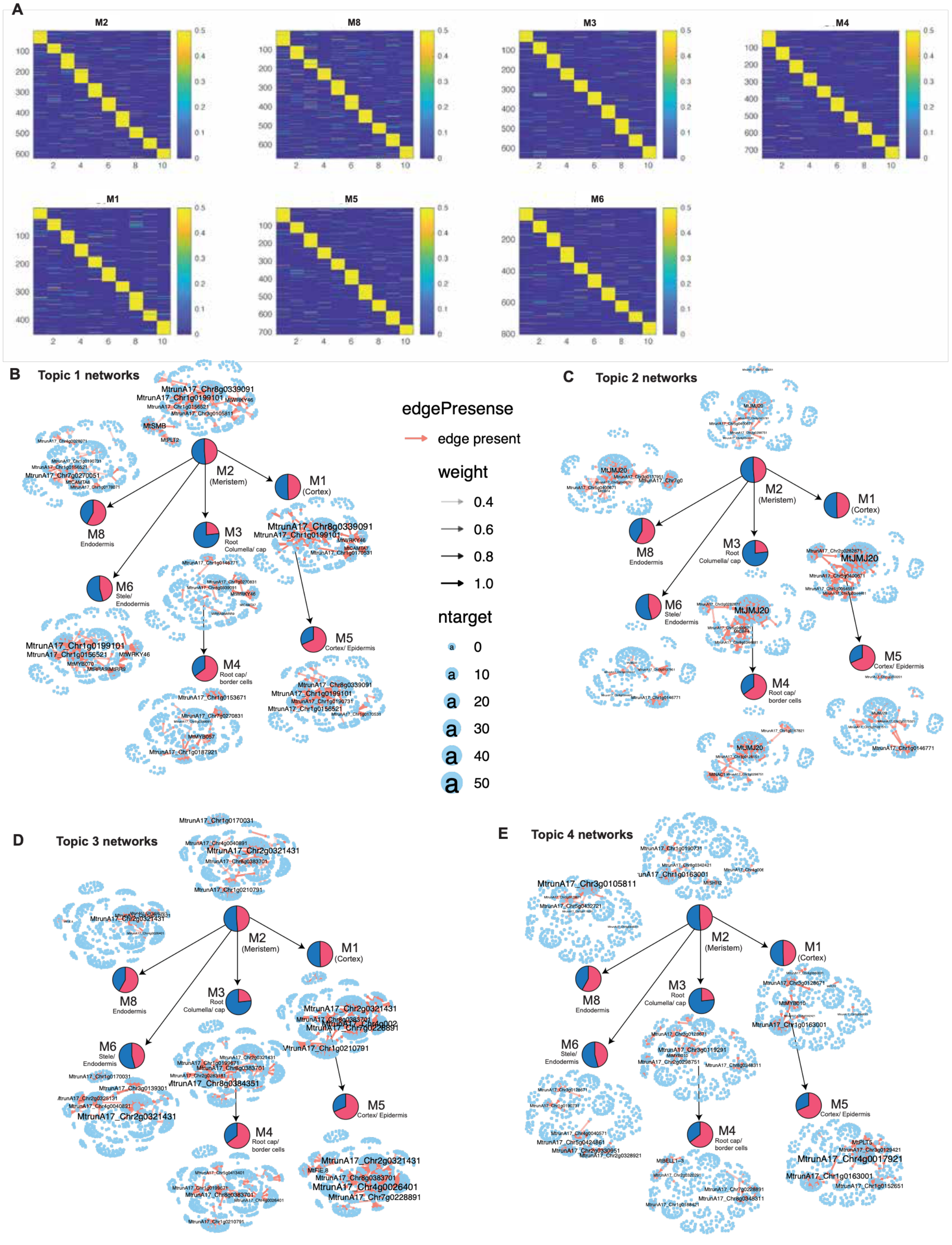

**Supplemental Figure 21: Summary statistics of MERLIN-inferred gene regulatory networks. A, C, E)** Predicted major regulators ordered by their number of targets for **A)** Medicago, **C)** maize, and **E)** sorghum. **B, D, F)** The distribution of MERLIN module sizes (number of genes per module) for **B)** Medicago, **D)** maize, and **F)** sorghum.

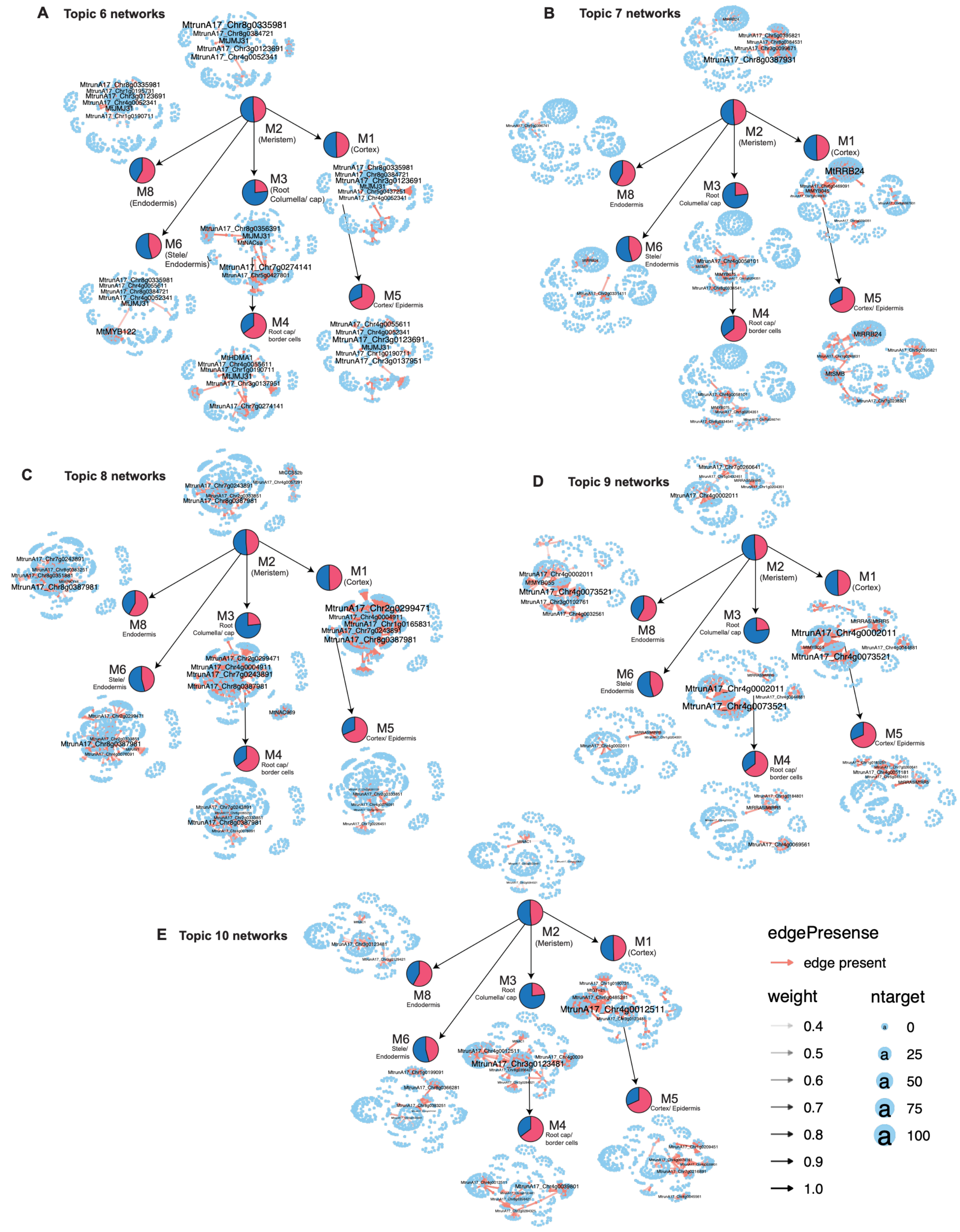

**Supplemental Figure 22: Topic-specific networks for each cell cluster** for **A)** topic 6, **B)** topic 7, **C)** topic 8, **D)** topic 9, **E)** topic 10

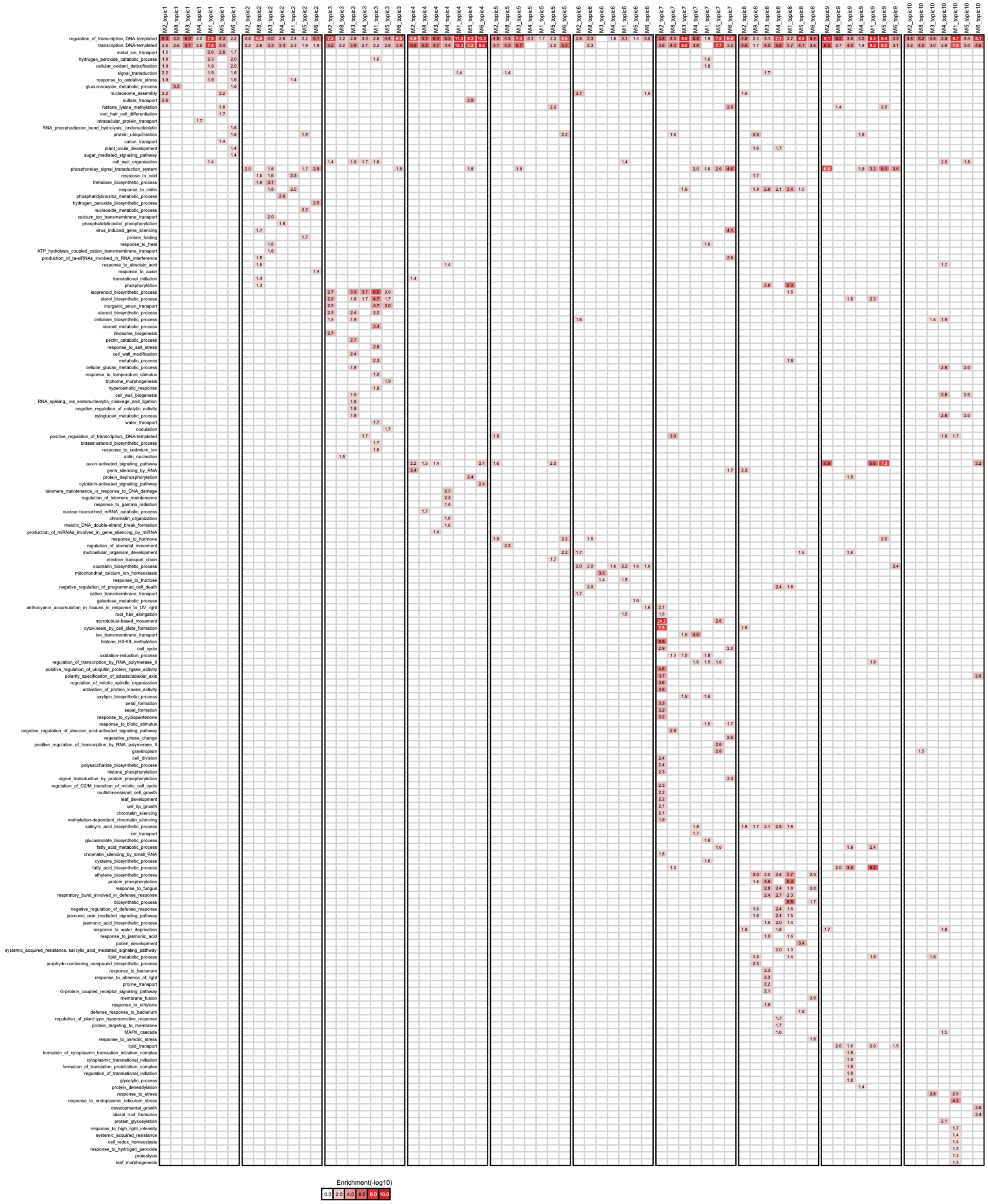

**Supplemental Figure 23: Gene Ontology (GO) process enrichment for genes associated with each LDA topic-specific network identified from Medicago.** We used FDR<0.01 to determine significantly enriched terms. The red intensity of the heatmap corresponds to −log10(FDR).

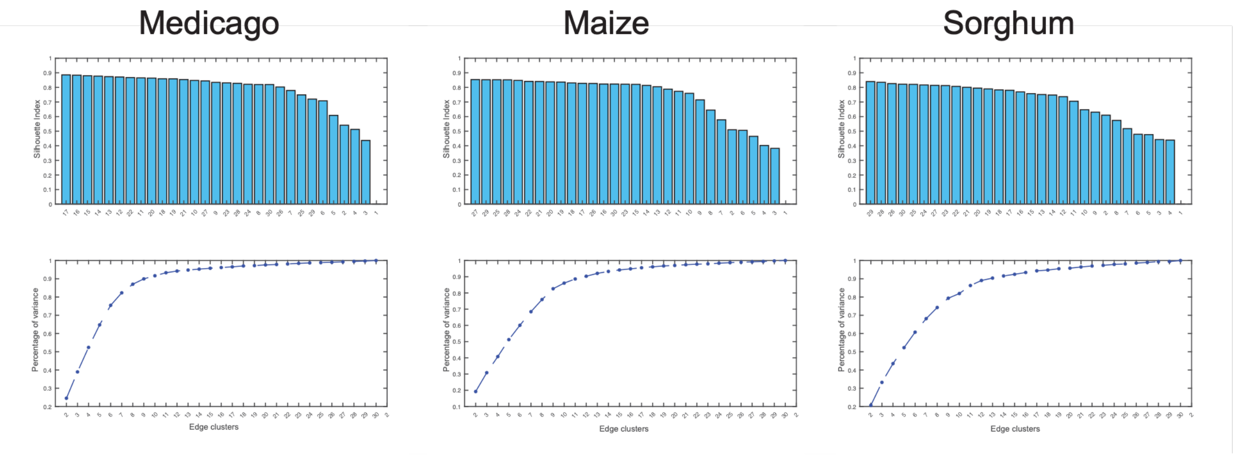

**Supplemental Figure 24: Selection of the number of edge clusters for each species.** Silhouette index (top) and cumulative percentage of variance explained (bottom) for different numbers of edge clusters in Medicago, maize, and sorghum. The candidate numbers of clusters are ordered according to their silhouette index in the top panels. Higher silhouette-index values indicate better separation and cohesion of edge clusters.

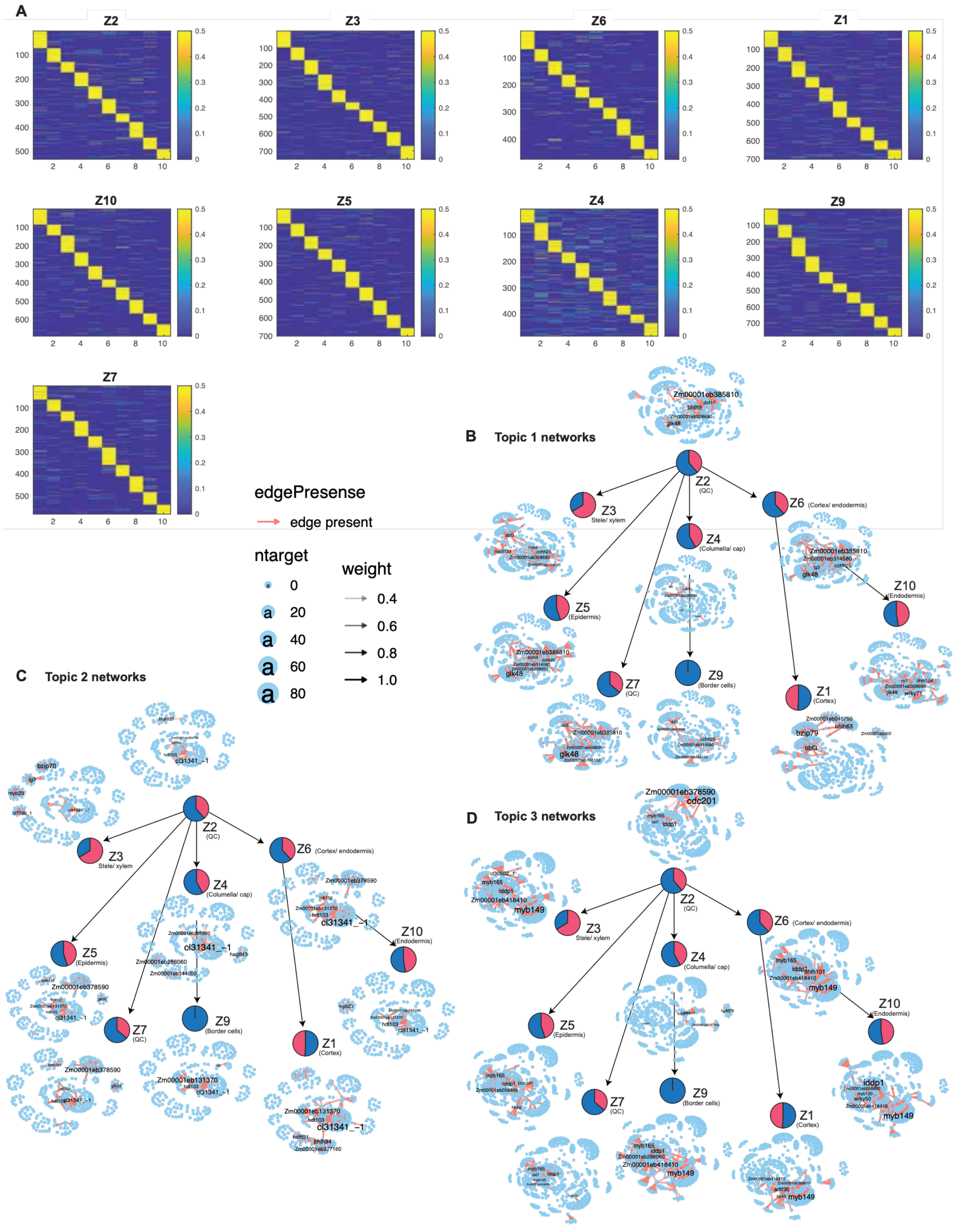

**Supplemental Figure 25: LDA analysis of scMTNI inferred networks of maize. A)** The document-topic weight matrix for all genes for all cell clusters with 10 topics. Topic-specific networks for each cell cluster for topic 1 **(B)**, topic 2 **(C)**, and topic 3 **(D)**.

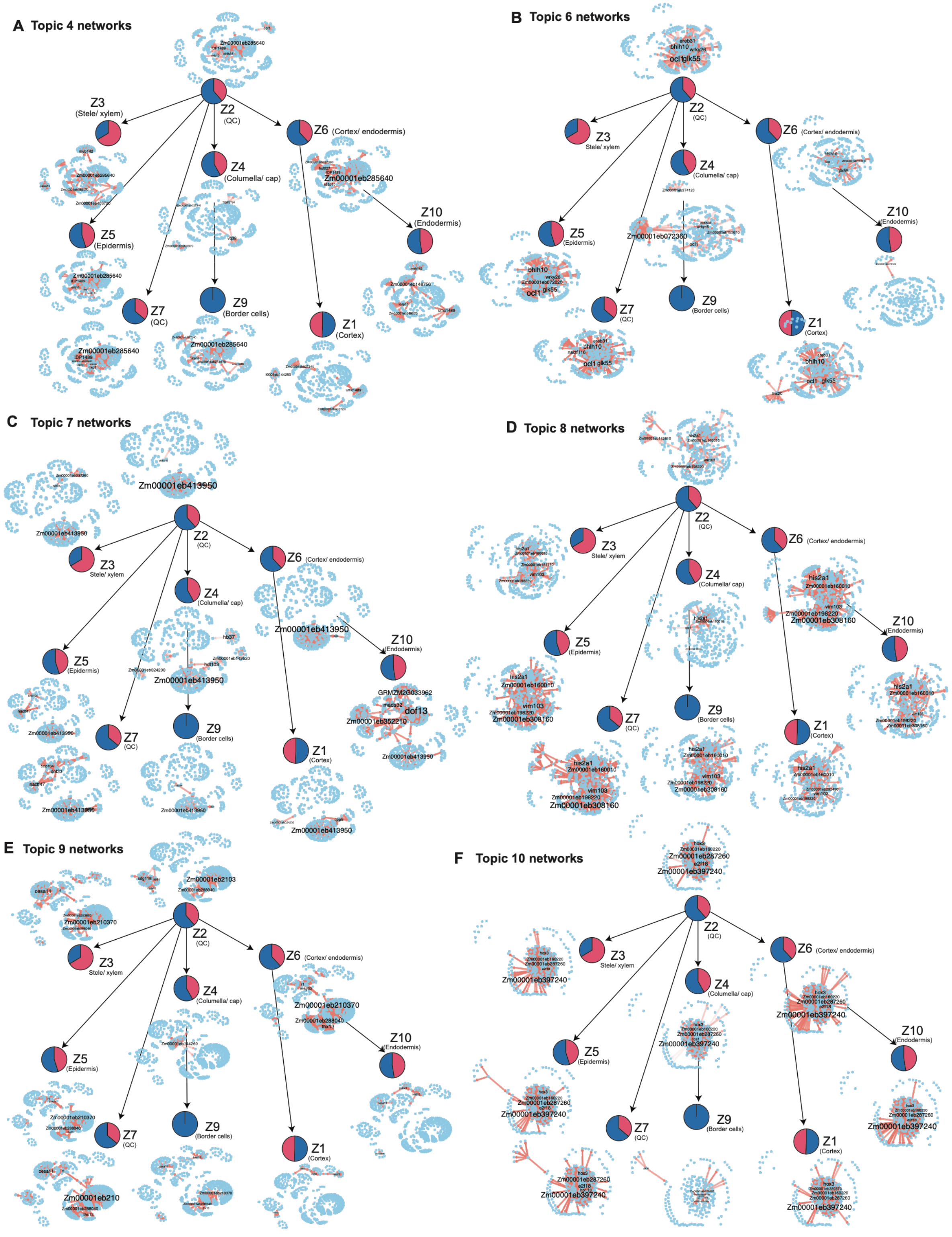

**Supplemental Figure 26: Topic-specific networks obtained from scMTNI-inferred networks for maize.** Shown are topic-specific networks for topic 4 **(A)**, topic 6 **(B)**, topic 7 **(C)**, topic 8 **(D)**, topic 9 **(E)**, and topic 10 **(F)**

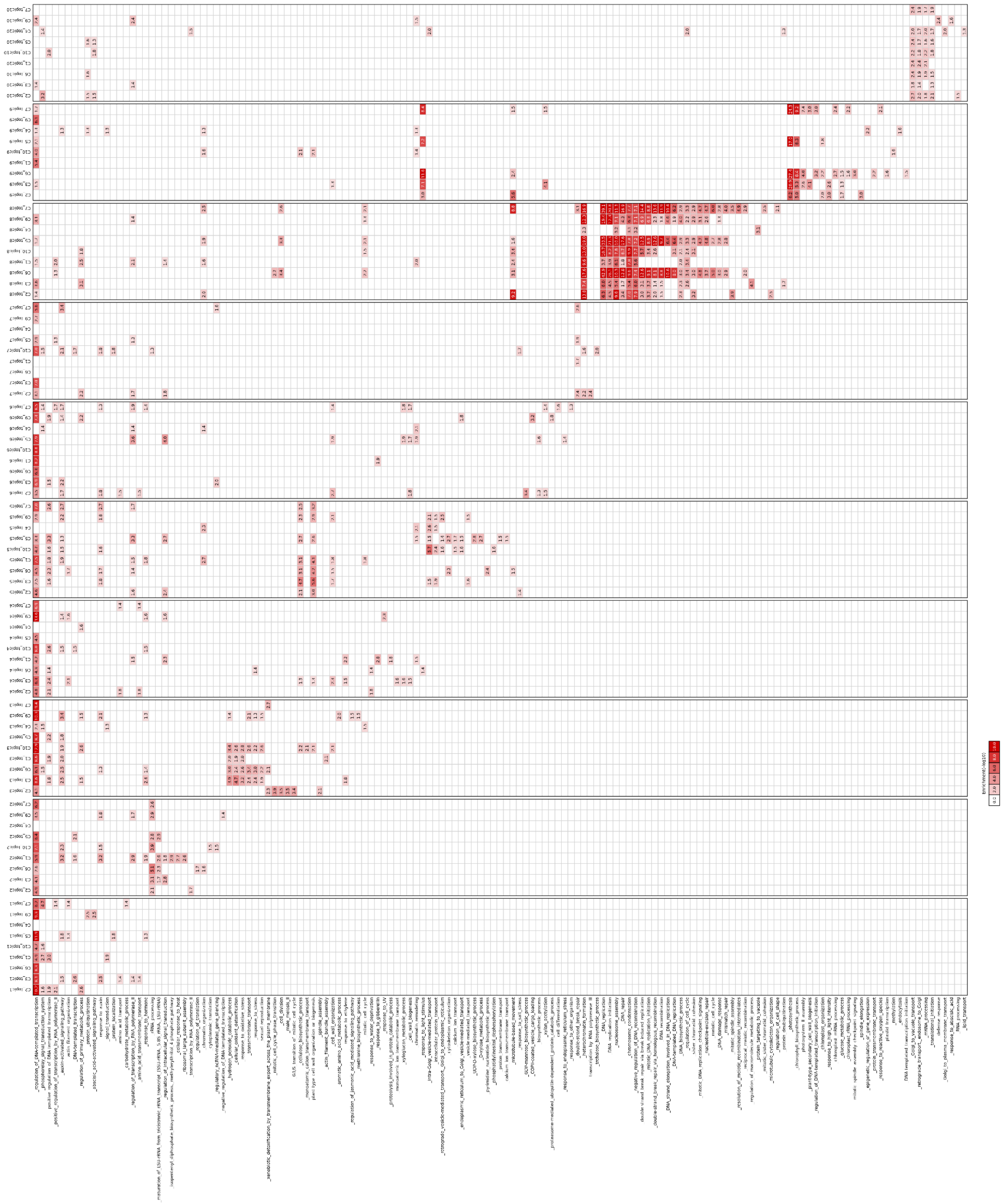

***Supplemental Figure 27:* GO enrichment for genes in each LDA topic-specific network inferred by scMTNI in maize.** We used FDR <0.01 to determine significantly enriched terms. The red intensity of the heat map is proportional to −log10(FDR) of enrichment.

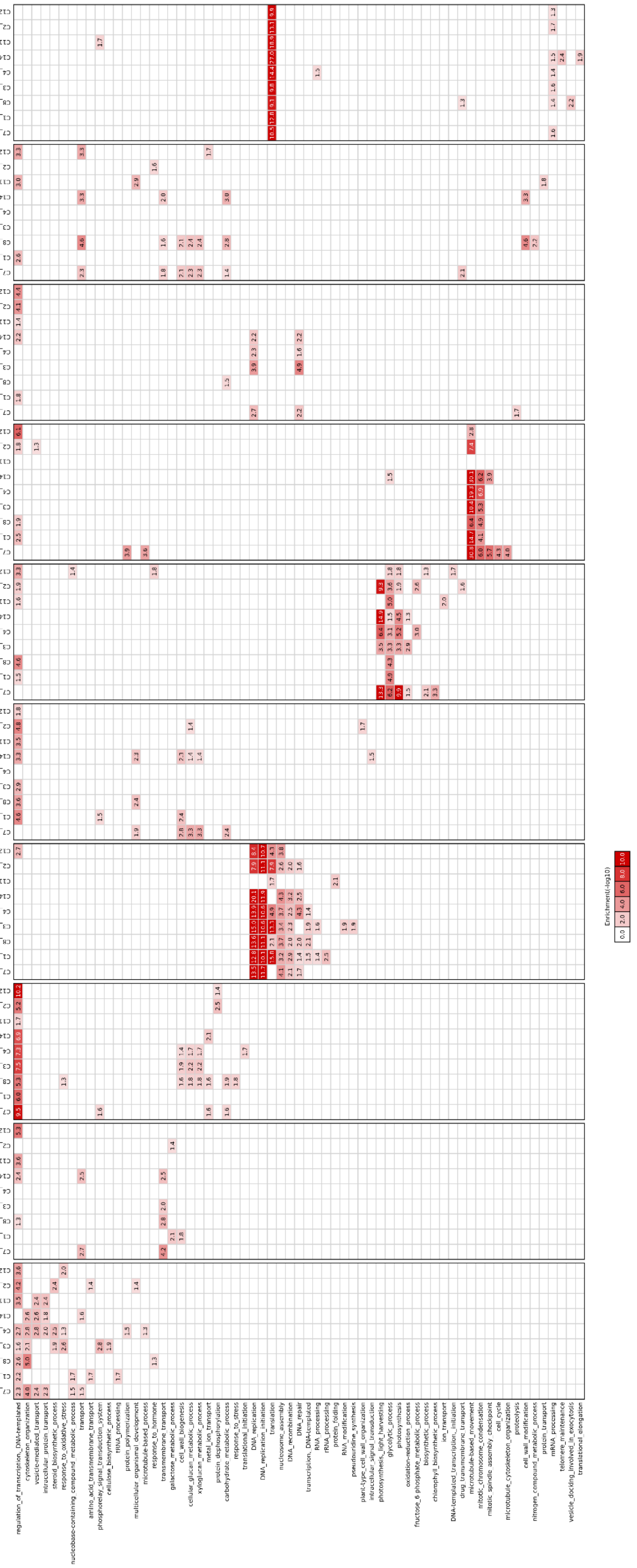

**Supplemental Figure 28: GO enrichment for genes in each LDA topic-specific network inferred by scMTNI on sorghum.** We used FDR <0.01 to determine significantly enriched terms. The red intensity of the heat map is proportional to −log10(FDR) of enrichment.

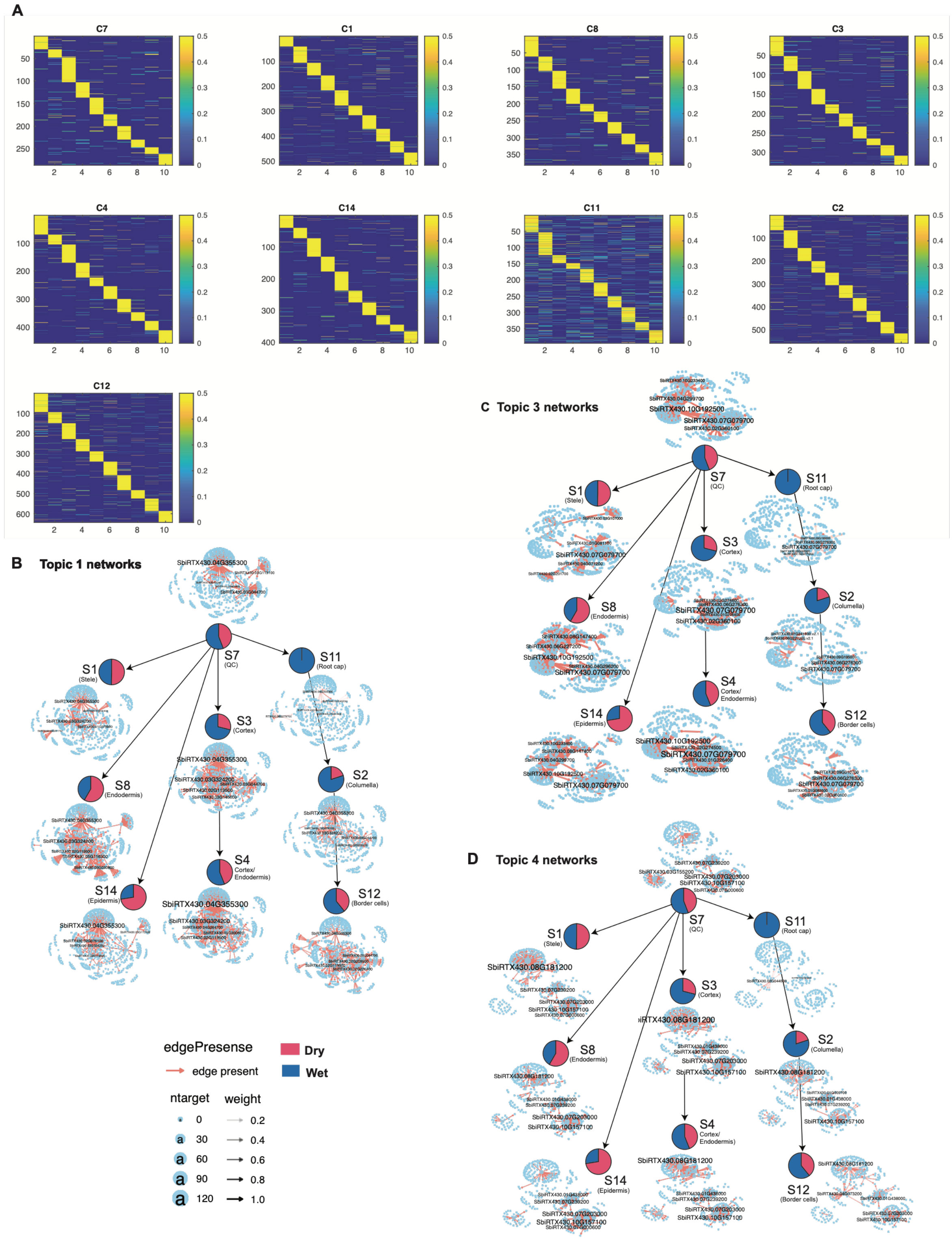

**Supplemental Figure 29: LDA analysis of scMTNI-inferred networks for sorghum. A. Document-topic weight matrix for all regulators across all cell clusters with 10 topics.** Topic-specific networks across each cell cluster for topic 1 **(B)**, topic 2 **(C)**, and topic 4 **(D)**.

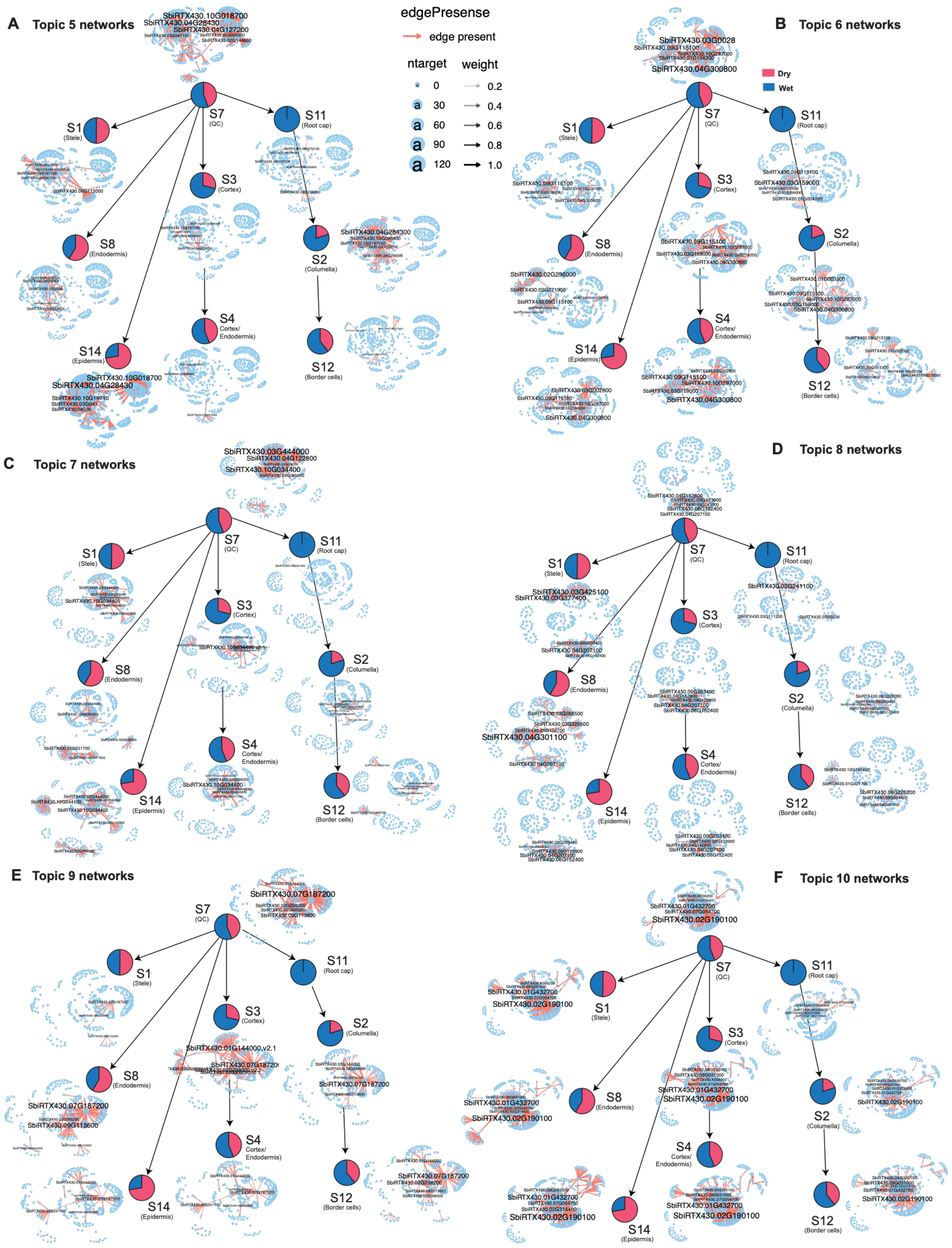

**Supplemental Figure 30: Topic-specific networks obtained from scMTNI-inferred networks for sorghum.** Shown are topic-specific networks for topic 5 **(A)**, topic 6 **(B)**, topic 7 **(C)**, topic 8 **(D)**, topic 9 **(E)**, and topic 10 **(F)**.

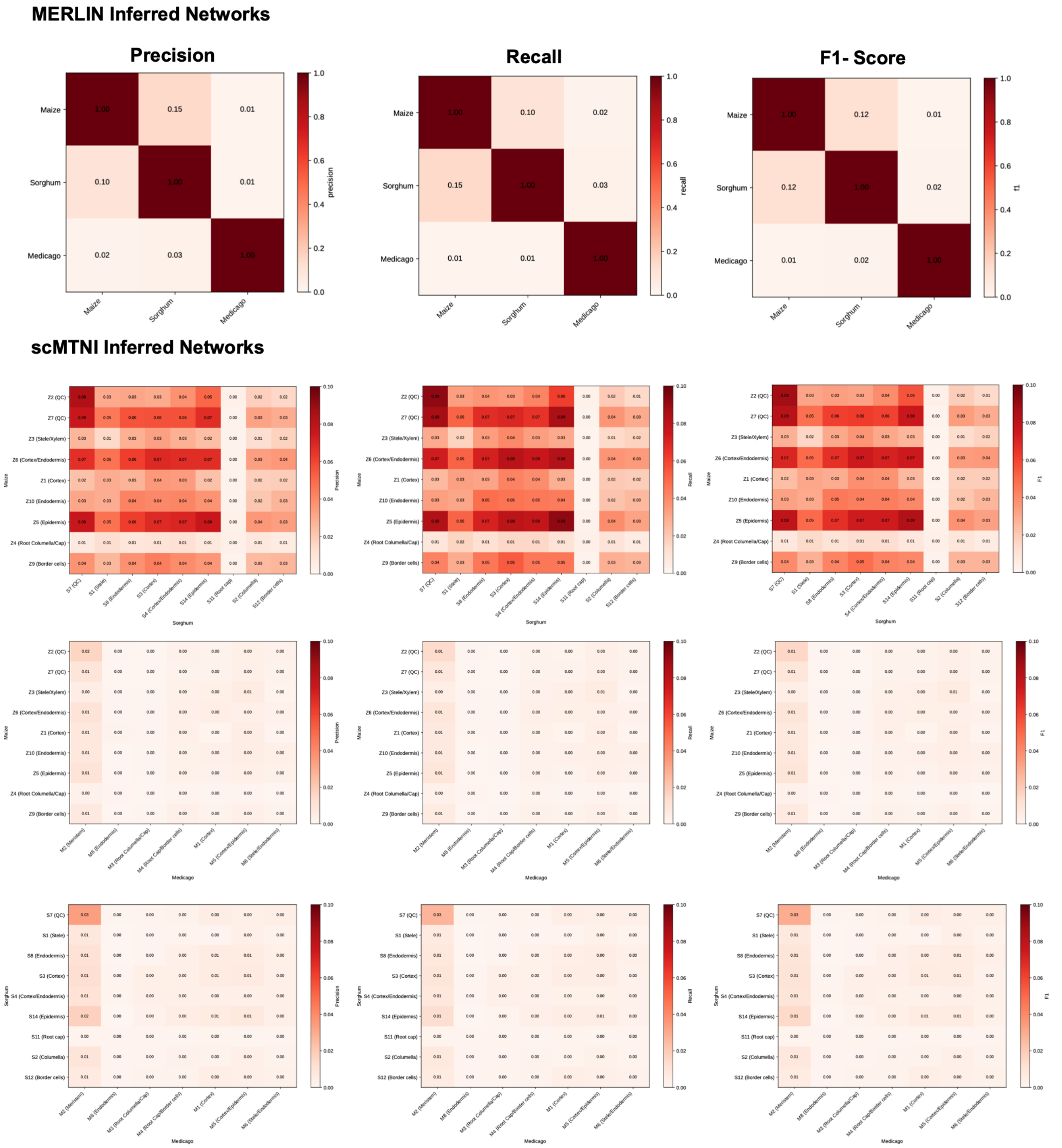

**Supplemental Figure 31: Comparison of inferred gene regulatory networks across species.** Precision, recall, and F1-score were used to quantify network similarity among maize, sorghum, and Medicago. The top panels show pairwise comparisons of species-level networks inferred using MERLIN. The lower panels show comparisons of cell-type-specific networks inferred using scMTNI for maize–sorghum, maize–Medicago, and sorghum–Medicago. Rows and columns represent the corresponding species or cell types, and darker colors indicate greater similarity between the inferred networks.

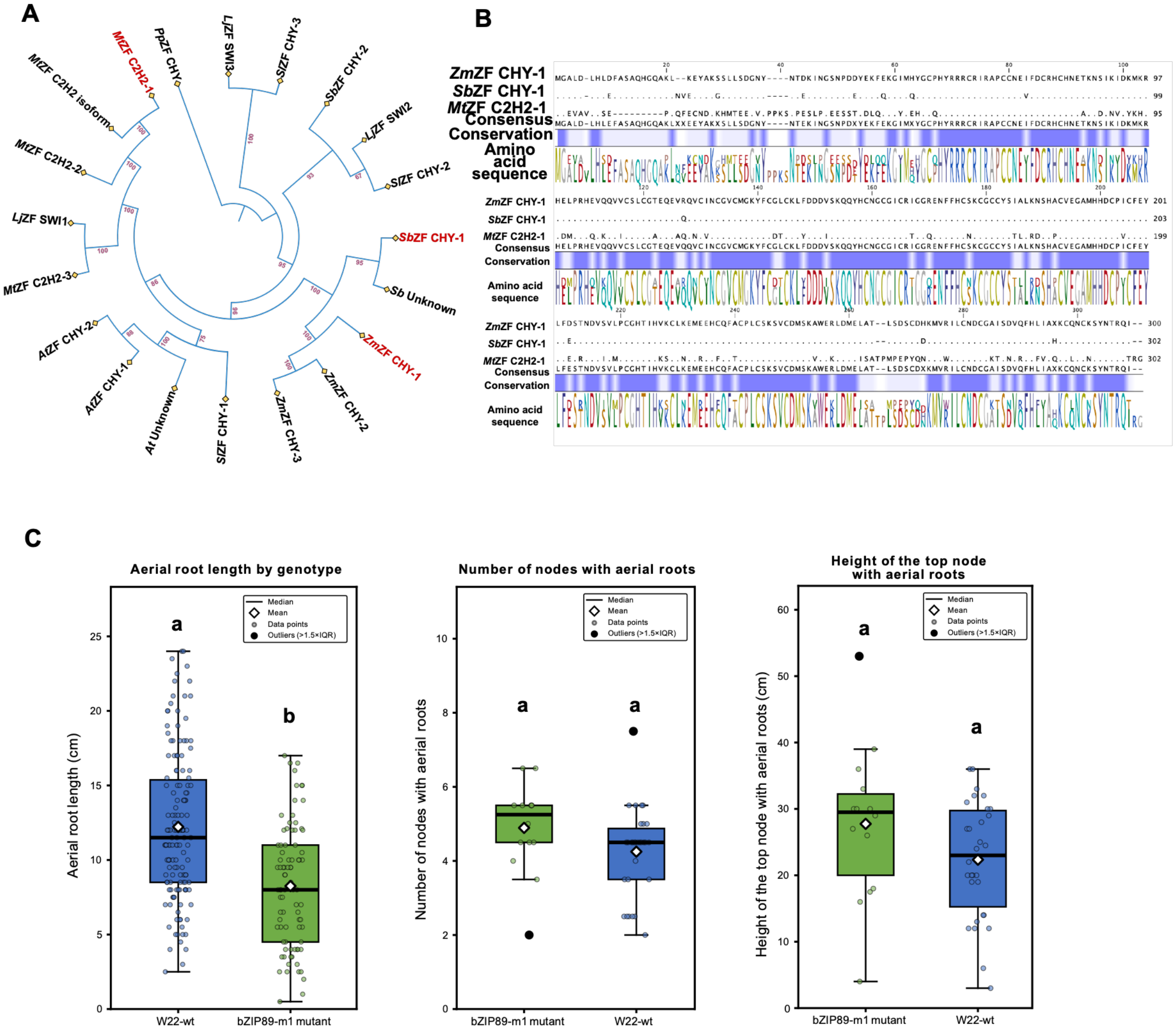

**Supplemental Figure 32: Phylogeny and protein alignment of the transcription factor C2H2.** The protein sequence from Medicago (Mt), maize (Zm), and sorghum (Sb), along with other plant species, was used to build a maximum-likelihood phylogenetic tree and a protein alignment. A). In addition to the three species, Arabidopsis thaliana (At), Solanum lycopersicum (Sl), and Physcomitrium patens (Pp) were included. Bootstrap values are included. **B)** Protein alignment showing the conserved amino acids and conserved domains (darker blue) in the three orthologs.

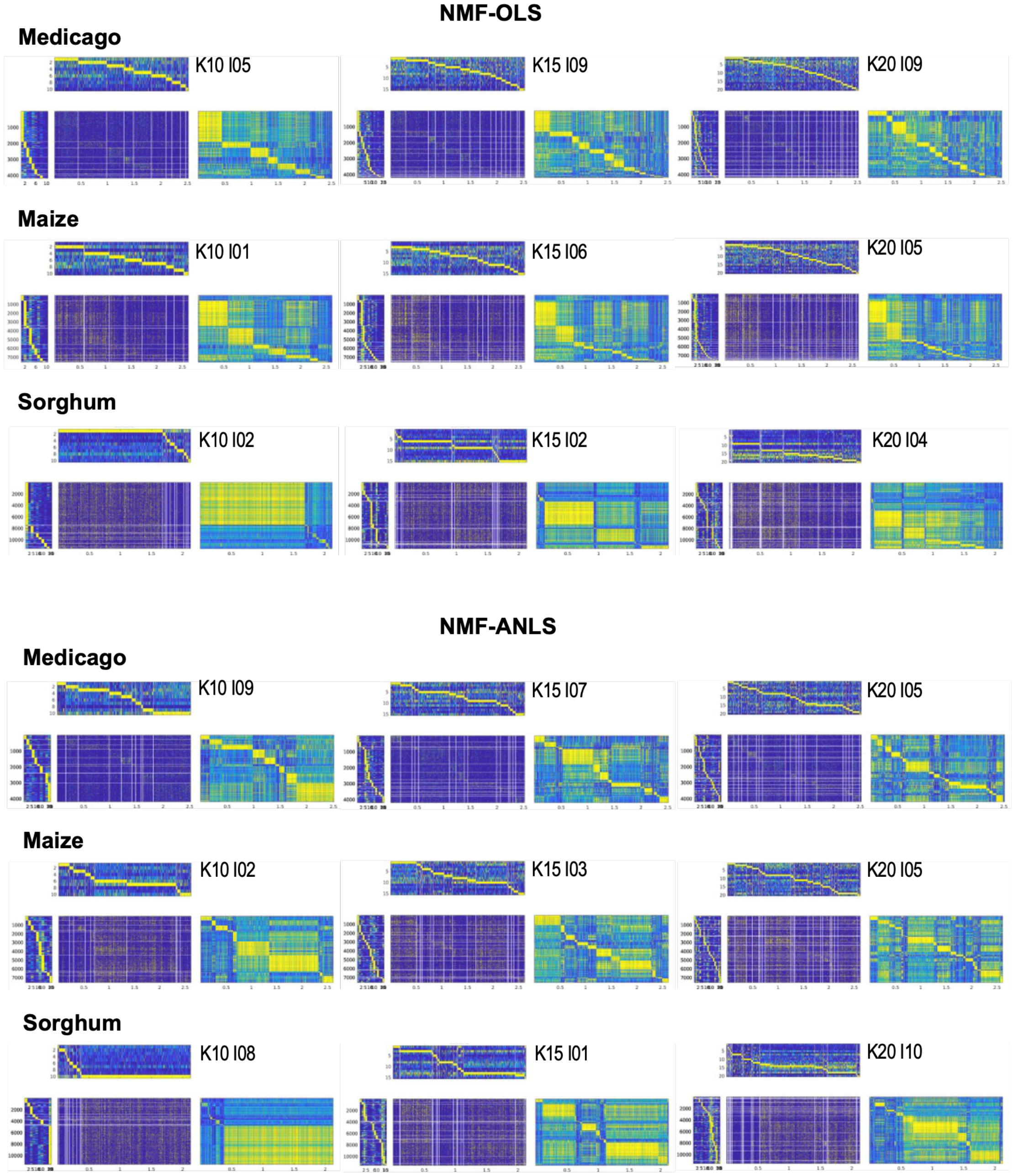

**Supplemental Figure 33: Comparison of NMF factorization methods across species. Non-negative matrix factorization (NMF) results for the gene expression matrices from Medicago, maize, and sorghum.** Two factorization approaches are shown: NMF-OLS (ordinary least squares; top) and NMF-ANLS (alternating non-negativity-constrained least squares; bottom). For each species, results are shown for three values of K (number of factors/clusters; K = 10, 15, and 20). For each configuration, the most stable random initialization is displayed. Each set of plots shows the factorization components: the original expression matrix (O), its NMF approximation (A), and the corresponding factor matrices U (cells × K) and V (K × genes). Color intensity indicates relative expression values from low (blue) to high (yellow).

### Supplemental Tables

**Supplemental Table 1:** snRNAseq cell summary in the dry and wet conditions in maize, sorghum, and Medicago.

| **Maize sample:** | **Dry** | **Wet** |
|---|---|---|
| Number of cells | 3,159 | 4,558 |
| Valid UMI | 99.90% | 99.90% |
| Valid Barcodes | 94.30% | 95.70% |
| Fraction read in cells | 76.10% | 82.70% |
| Median Genes per Cell | 1,806 | 2,808 |
| Mean Reads per Cell | 78,524 | 56,958 |
| Sequencing saturation | 80.60% | 72.80% |
| Reads Mapped to Genome | 46.00% | 63.40% |
| **Sorghum sample:** | | |
| Number of cells | 5,506 | 6,412 |
| Valid UMI | 100.00% | 100.00% |
| Valid Barcodes | 97.00% | 97.10% |
| Fraction read in cells | 80.40% | 86.30% |
| Median Genes per Cell | 2,694 | 2,564 |
| Mean Reads per Cell | 52,410 | 23,830 |
| Sequencing saturation | 81.80% | 67.90% |
| Reads Mapped to Genome | 87.20% | 90% |
| **Medicago sample:** | | |
| Number of cells | 2,129 | 2,563 |
| Valid UMI | 99.90% | 99.90% |
| Valid Barcodes | 97.30% | 95.80% |
| Fraction read in cells | 77.20% | 55.70% |
| Median Genes per Cell | 2,604 | 1,770 |
| Mean Reads per Cell | 57,256 | 42,221 |
| Sequencing saturation | 81.70% | 65.60% |
| Reads Mapped to Genome | 87.30% | 90.20% |

**Supplemental Table 2:** Network statistics for gold standard datasets from three species, generated using ATAC-seq for maize and Medicago, and ChIP-seq for sorghum.

| Medicago | | | |
|---|---|---|---|
| **Network** | **# regulators** | **# targets** | **# edges** |
| ATAC_GSE101482(U1_D1) | 350 | 13, 985 | 379,688 |
| ATAC_GSE101482(U2_D2) | 350 | 17, 714 | 514,240 |
| ATAC_GSE101482(U3_D3) | 350 | 20, 815 | 631,182 |
| ATAC_GSE101482(U4_D4) | 350 | 23, 559 | 742,305 |
| ATAC_GSE101482(U5_D5) | 350 | 25, 948 | 848,485 |

| Medicago | | | |
|---|---|---|---|
| **Network** | **# regulators** | **# targets** | **# edges** |
| ATAC_GSE154845(U1_D1) | 350 | 14,714 | 530,712 |
| ATAC_GSE154845(U2_D2) | 350 | 18,435 | 699,724 |
| ATAC_GSE154845(U3_D3) | 350 | 21,522 | 847,361 |
| ATAC_GSE154845(U4_D4) | 350 | 24,259 | 981,138 |
| ATAC_GSE154845(U5_D5) | 350 | 26,691 | 1,110,641 |

| Maize | | | |
|---|---|---|---|
| **Network** | **# regulators** | **# targets** | **# edges** |
| ATAC_GSE152046(U1_D1) | 1095 | 22, 727 | 1,275,366 |
| ATAC_GSE152046(U2_D2) | 1095 | 25, 417 | 1,559,944 |
| ATAC_GSE152046(U3_D3) | 1095 | 27, 284 | 1,773,442 |
| ATAC_GSE152046(U4_D4) | 1095 | 28, 622 | 1,964,352 |
| ATAC_GSE152046(U5_D5) | 1095 | 29, 681 | 2,142,743 |

| Sorghum | | | |
|---|---|---|---|
| **Network** | **# regulators** | **# targets** | **# edges** |
| ChipSeq_PRJNA952350(U1_D1) | 102 | 16, 757 | 143, 850 |
| ChipSeq_PRJNA952350(U2_D2) | 102 | 18, 054 | 157, 945 |
| ChipSeq_PRJNA952350(U3_D3) | 102 | 19, 269 | 173, 306 |
| ChipSeq_PRJNA952350(U4_D4) | 102 | 20, 460 | 190, 577 |
| ChipSeq_PRJNA952350(U5_D5) | 102 | 21, 534 | 210, 512 |

**Supplemental Table 3:** Candidate regulators selected for the experimental validation from medicago and maize

| **Medicago candidate regulators** | | | | | | | | | | |
|---|---|---|---|---|---|---|---|---|---|---|
| **Common name** | **Medicago validated regulator** | **MERLIN module if any** | **scMTNI kmeans** | **scMTNI LDA** | **Sorghum ortholog** | **Sorghum kmeans** | **Sorghum topic** | **Maize ortholog** | **Maize kmeans** | **Maize scMTNI LDA** |
| C2H2 zinc finger | MtrunA17_Chr1g0148421 | | E14 | - | SbiRTX430.03G300600 | E8 | topic1 | Zm00001eb154580 (umc1489) | - | topic4 |
| WOX Homobox | MtrunA17_Chr5g0400781 | module18236 | | 1,2,3,9 | SbiRTX430.01G292900 | - | - | Zm00001eb025650 | - | - |
| TUBBY | MtrunA17_Chr1g0201891 | module18227 | - | 5 | SbiRTX430.07G002500 | - | - | Zm00001eb198820 (tubtf7) | - | - |
| | | | | | SbiRTX430.06G296200 | - | - | Zm00001eb076160 (tubtf4) | - | - |
| | | | | | SbiRTX430.09G159400 | - | - | Zm00001eb146720 (tubtf2) | - | - |
| | | | | | SbiRTX430.03G396900 | - | - | Zm00001eb407680 (tubtf14) | - | - |
| | | | | | SbiRTX430.08G050400 | - | - | Zm00001eb202420 (tubtf3) | - | - |
| | | | | | SbiRTX430.05G046700 | - | - | | - | - |
| MYB | MtrunA17_Chr3g0111941 | | | 1,3,6 | - | - | - | - | - | - |
| Kruppel like | MtrunA17_Chr1g0207451 | module18227 | E14 | 5 | - | - | - | - | - | - |
| **Maize candidate regulator** | | | | | | | | | | |
| **Common name** | **maize validated regulator** | **MERLIN module if any** | **scMTNI kmeans** | **scMTNI LDA** | **Sorghum ortholog** | **Sorghum kmeans** | **Sorghum topic** | **Medicago ortholog** | **Medicago kmeans** | **Medicago scMTNI LDA** |
| ***Zmbzip89*** | Zm00001eb238570 | - | - | Topic 5 | - | - | - | - | - | - |

**Supplemental Table 4:** List of plasmids used in this study

| Plasmid | Description | Reference |
|---|---|---|
| pJET1.2 | Vector for blunt end cloning, $Amp^R$ | Thermofisher Scientific (Cat K240020) |
| LIII-TM- lacZdy NLS2xYFP | Binary expression vector for promoter NLS-2xYFP fusions with NLS-2xmCherry transformation marker, KmR | |
| pENTR™/D-TOPO™ | Directional cloning vector for entry to the Gateway System | Thermofisher Scientific (Cat K240020) |
| pK7GW1GW2 – II – RedRoot | RNAi expression vector with TdTomato as transformation marker and the RNAi sequence is under the *Mt*Ubiquitin promoter $Spec^R$ | |
| pJET1.2-$RootCap_{pro}$ | pJET1.2 carrying a 2kb fragment upstream of the gene MtrunA17_Chr4g0011791 | This study |
| pJET1.2-$RootCap\_BC_{pro}$ | pJET1.2 carrying a 2kb fragment upstream of the gene MtrunA17_Chr8g0379831 | This study |
| pJET1.2-$Endodermis_{pro}$ | pJET1.2 carrying a 2kb fragment upstream of the gene MtrunA17_Chr4g0321181 | This study |
| MtA17_$RootCap_{pro}$:NLS-2xYFP | LIII vector expressing the NLS-2xYFP reporter under the control of the 2kb MtA17_$RootCap_{pro}$ promoter (MtrunA17_Chr4g0011791) | This study |
| MtA17_$RootCap_BC_{pro}$:NLS-2xYFP | LIII vector expressing the NLS-2xYFP reporter under the control of the 2kb MtA17_$RootCap_{pro}$ promoter (MtrunA17_Chr8g0379831) | This study |
| MtA17_$Endodermis_{pro}$:NLS-2xYFP | LIII vector expressing the NLS-2xYFP reporter under the control of the 2kb MtA17_$RootCap_{pro}$ promoter (MtrunA17_Chr4g0321181) | This study |
| pENTR – Mt0781 | Gateway vector containing a sequence of the gene MtrunA17_Chr5g0400781 to generate a knockout line | This study |
| pENTR – Mt7451 | Gateway vector containing a sequence of the gene MtrunA17_Chr1g0207451 to generate a knockout line | This study |
| pENTR – Mt1941 | Gateway vector containing a sequence of the gene MtrunA17_Chr3g0111941 to generate a knockout line | This study |
| pENTR – Mt8421 | Gateway vector containing a sequence of the gene MtrunA17_Chr1g0148421 to generate a knockout line | This study |

| | | |
|---|---|---|
| pENTR – Mt1891 | Gateway vector containing a sequence of the gene MtrunA17_Chr1g0201891 to generate a knockout line | This study |
| pK7GW1GW2 – Mt0781 | RNAi expression vector with TdTomato as transformation marker and the RNAi Mt0781 (MtrunA17_Chr5g0400781) sequence under the MtUbiquitin promoter SpecR | This study |
| pK7GW1GW2 – Mt7451 | RNAi expression vector with TdTomato as transformation marker and the RNAi Mt7451 (MtrunA17_Chr1g0207451) sequence under the MtUbiquitin promoter SpecR | This study |
| pK7GW1GW2 – Mt1941 | RNAi expression vector with TdTomato as transformation marker and the RNAi Mt1941 (MtrunA17_Chr3g0111941) sequence under the MtUbiquitin promoter SpecR | This study |
| pK7GW1GW2 – Mt8421 | RNAi expression vector with TdTomato as transformation marker and the RNAi Mt8421 (MtrunA17_Chr1g0148421) sequence under the MtUbiquitin promoter SpecR | This study |
| pK7GW1GW2 – Mt1891 | RNAi expression vector with TdTomato as transformation marker and the RNAi Mt1891 (MtrunA17_Chr1g0201891) sequence under the MtUbiquitin promoter SpecR | This study |

**Supplemental Table 5:** List of promoter, maize, and sorghum RT-qPCR and Medicago RNAi primer sequences. The Esp3I restriction site is underlined and in bold, the overhang to clone into pENTR Gateway vector

| **Primer** | **Sequence (5' – 3')** |
|---|---|
| **Promoter** | |
| MtA17_promRootCap_BC_F (MtrunA17_Chr4g0011791) | ATCGTCTCTGCGGGTTATTGAGACCGAAAGTGT |
| MtA17_promRootCap_BC_R (MtrunA17_Chr4g0011791) | ATCGTCTCTCAGAGTTATTGAGACCGAAAGTGT |
| MtA17_promRootCap_F (MtrunA17_Chr8g0379831) | ATCGTCTCAGCGGCGTTGGCCATTTATGAAACC |
| MtA17_promRootCap_R (MtrunA17_Chr8g0379831) | ATCGTCTCACAGAAGTTAATTGAAGAAAAAGAAACAATTGG |
| MtA17_promEndodermis_F (MtrunA17_Chr4g0321181) | ATCGTCTCTGCGGGATACACACGTTTACTGG |
| MtA17_promRootCap_BC_F (MtrunA17_ Chr4g0321181) | ATCGTCTCACAGATTGAAATAAAGGAACAAATT |
| **RTqPCR sorghum** | |
| Sb_BC_F (SbiRTX430.03G383000) | GGCGAGCCATTCTTTGTCCT |
| Sb_BC_R (SbiRTX430.03G383000) | TGTTGCCTCACCACCATGAC |
| Sb_RootCap_F (SbiRTX430.04G115800) | GCTGGAGATCGTCGAGGTG |
| Sb_RootCap_F (SbiRTX430.04G115800) | CCATGCTGCTGCTGATGATG |
| Sb_Cortex_F (SbiRTX430.03G38500) | ATACATCGTCAAAACCCCAA |
| Sb_Cortex_R (SbiRTX430.03G38500) | GAGACGACCTACGACTGT |
| **RTqPCR maize** | |
| Zm_Endodermis_F (Zm00001eb005460) | TCCATCGTCATCCCTCAGGT |
| Zm_Endodermis_R (Zm00001eb005460) | AGATCTTGGGCAGCAGGAAC |
| Zm_RootCap_F (Zm00001eb163560) | TGTTCTCCAACCTGTCCATCG |
| Zm_RootCap_F (Zm00001eb163560) | CTGGTGTCCCGGGAGTAGTG |
| Zm_Cortex_F (Zm00001eb027760) | GGGGCGACCATTATTCCTCTT |
| Zm_Cortex_F (Zm00001eb027760) | CGGGCGGAGGGATGACATT |
| **RNAi** | |
| Mt0781_F (MtrunA17_Chr5g0400781) | **CACC**GGATTGCGAGTTCTTG |
| Mt0781_R (MtrunA17_Chr5g0400781) | CGTACGTTTCCCTAGC |
| Mt7451_F (MtrunA17_Chr1g0207451) | **CACC**CGATCAAAACGACCGC |
| Mt7451_R (MtrunA17_Chr1g0207451) | GCGTCGGAGACTGTAA |
| Mt1941_F (MtrunA17_Chr3g0111941) | **CACC**GTACATACCGAGACTGG |
| Mt1941_R (MtrunA17_Chr3g0111941) | GTCCTTCAACTTTGGAGC |
| Mt8421_F (MtrunA17_Chr1g0148421) | **CACC**CGGGTTAGGGCAAGTAG |

| | |
|---|---|
| Mt8421_R (MtrunA17_Chr1g0148421) | CATGTTGGTGATAACCGG |
| Mt1891_F (MtrunA17_Chr1g0201891) | **CACC**CGTGACTGTTGCGTCTG |
| Mt1891_R (MtrunA17_Chr1g0201891) | GGACATTGGGAAGAGATTG |
